# Exploiting the GTEx resources to decipher the mechanisms at GWAS loci

**DOI:** 10.1101/814350

**Authors:** Alvaro N Barbeira, Rodrigo Bonazzola, Eric R Gamazon, Yanyu Liang, YoSon Park, Sarah Kim-Hellmuth, Gao Wang, Zhuoxun Jiang, Dan Zhou, Farhad Hormozdiari, Boxiang Liu, Abhiram Rao, Andrew R Hamel, Milton D Pividori, François Aguet, GTEx GWAS Working Group, Lisa Bastarache, Daniel M Jordan, Marie Verbanck, Ron Do, GTEx Consortium, Matthew Stephens, Kristin Ardlie, Mark McCarthy, Stephen B Montgomery, Ayellet V Segrè, Christopher D. Brown, Tuuli Lappalainen, Xiaoquan Wen, Hae Kyung Im

## Abstract

The resources generated by the GTEx consortium offer unprecedented opportunities to advance our understanding of the biology of human diseases. Here, we present an in-depth examination of the phenotypic consequences of transcriptome regulation and a blueprint for the functional interpretation of genome-wide association study-discovered loci. Across a broad set of complex traits and diseases, we demonstrate widespread dose-dependent effects of RNA expression and splicing. We develop a data-driven framework to benchmark methods that prioritize causal genes and find no single approach outperforms the combination of multiple approaches. Using colocalization and association approaches that take into account the observed allelic heterogeneity of gene expression, we propose potential target genes for 47% (2,519 out of 5,385) of the GWAS loci examined. Our results demonstrate the translational relevance of the GTEx resources and highlight the need to increase their resolution and breadth to further our understanding of the genotype-phenotype link.

## Introduction

In the last decade, the number of reproducible genetic associations with complex human traits that have emerged from genome-wide association studies (GWAS) has substantially grown. Many of the identified associations lie in non-coding regions of the genome, suggesting that they influence disease pathophysiology and complex traits via gene regulatory changes. Integrative studies of molecular quantitative trait loci (QTL) [Nicolae et al., 2010] have established gene expression as a key intermediate molecular phenotype, and improved functional interpretation of GWAS findings, spanning immunological diseases [Guo et al., 2015], various cancers [Wu et al., 2018; Gong et al., 2018], lipid traits [Pashos et al., 2017; Caliskan et al., 2019], and a broad array of other complex traits.

Large-scale international efforts such as the Genotype-Tissue Expression (GTEx) Consortium have provided an atlas of the regulatory landscape of gene expression and splicing variation in a broad collection of primary human tissues [Carithers et al., 2015; GTEx Consortium et al., 2017; Aguet et al., 2019]. Nearly all protein-coding genes in the genome now have at least one local variant associated with expression changes and the majority also have common variants affecting alternative splicing (FDR < 5%) [Aguet et al., 2019]. In parallel, there has been an explosive growth in the number of genetic discoveries across a large number of traits, prompting the development of integrative approaches to characterize the function of GWAS findings [Barbeira et al., 2019; Gamazon et al., 2018; Zhu et al., 2016; Gusev et al., 2016; Wen, 2016]. Nevertheless, our understanding of underlying biological mechanisms for most complex traits substantially lags behind the improved efficiency of the discovery of genetic associations, made possible by large-scale biobanks and GWAS meta-analyses.

One of the primary tools for the functional interpretation of GWAS associations has been the integrative analysis of molecular QTLs. Colocalization approaches that seek to establish shared causal variants (e.g., eCaviar [Hormozdiari et al., 2016], *enloc* [Wen et al., 2017], and *coloc* [Giambartolomei et al., 2014]), enrichment analysis (S-LDSC [Bulik-Sullivan et al., 2015] and QTLEnrich [Gamazon et al., 2018]) or mediation and association methods (SMR [Zhu et al., 2016], TWAS [Gusev et al., 2016] and PrediXcan [Gamazon et al., 2015]) have provided important insights, but they are often used in isolation, and there have been limited prior assessments of power and error rates associated with each [Wainberg et al., 2019]. Their applications often fail to provide a comprehensive, biologically interpretable view across multiple methods, traits, and tissues or offer guidelines that are generalizable to other contexts. Thus, a comprehensive assessment of expression and splicing QTLs for their contributions to disease susceptibility and other complex traits requires the development of novel methodologies with improved resolution and interpretability.

Here, we develop novel methods, approaches, and resources that elucidate how genetic variants associated with gene expression (cis-eQTLs) or splicing (cis-sQTLs) contribute to, or mediate, the functional mechanisms underlying a wide array of complex diseases and quantitative traits. Since splicing QTLs have largely been understudied, we perform a comprehensive integrative study of this class of QTLs, in a broad collection of tissues, and disease associations. We provide predictions of functional mechanisms for 74 distinct complex traits from 87 GWA study results and demonstrate independent validation and evaluation of findings using likely causal gene-disease relationships in the Online Mendelian Inheritance of Man (OMIM) database. Notably, we find widespread dose-dependent effects of cis-QTLs on traits through multiple lines of evidence. We examine the importance of considering, or correcting for, false functional links attributed to GWAS loci due to neighboring but distinct causal variants. We call this confounding LD contamination for the remainder of the paper. To identify predicted causal effects among the complex trait associated QTLs, we conduct systematic evaluation across different methods. Furthermore, we provide guidelines for employing complementary methods to map the regulatory mechanisms underlying genetic associations with complex traits.

### Mapping the regulatory landscape of complex traits

The final GTEx data release (v8) included 54 primary human tissues, 49 of which included at least 70 samples with both whole genome sequencing (WGS) and tissue-specific RNA-seq data. A total of 15,253 samples from 838 individuals were used for cis-QTL mapping (Fig. 1) [Aguet et al., 2019]. In addition to the expression quantitative trait loci (eQTL) mapping, we also evaluated genetic variation associated with alternative splicing (sQTL) and their impact on complex traits.

**Fig. 1.**
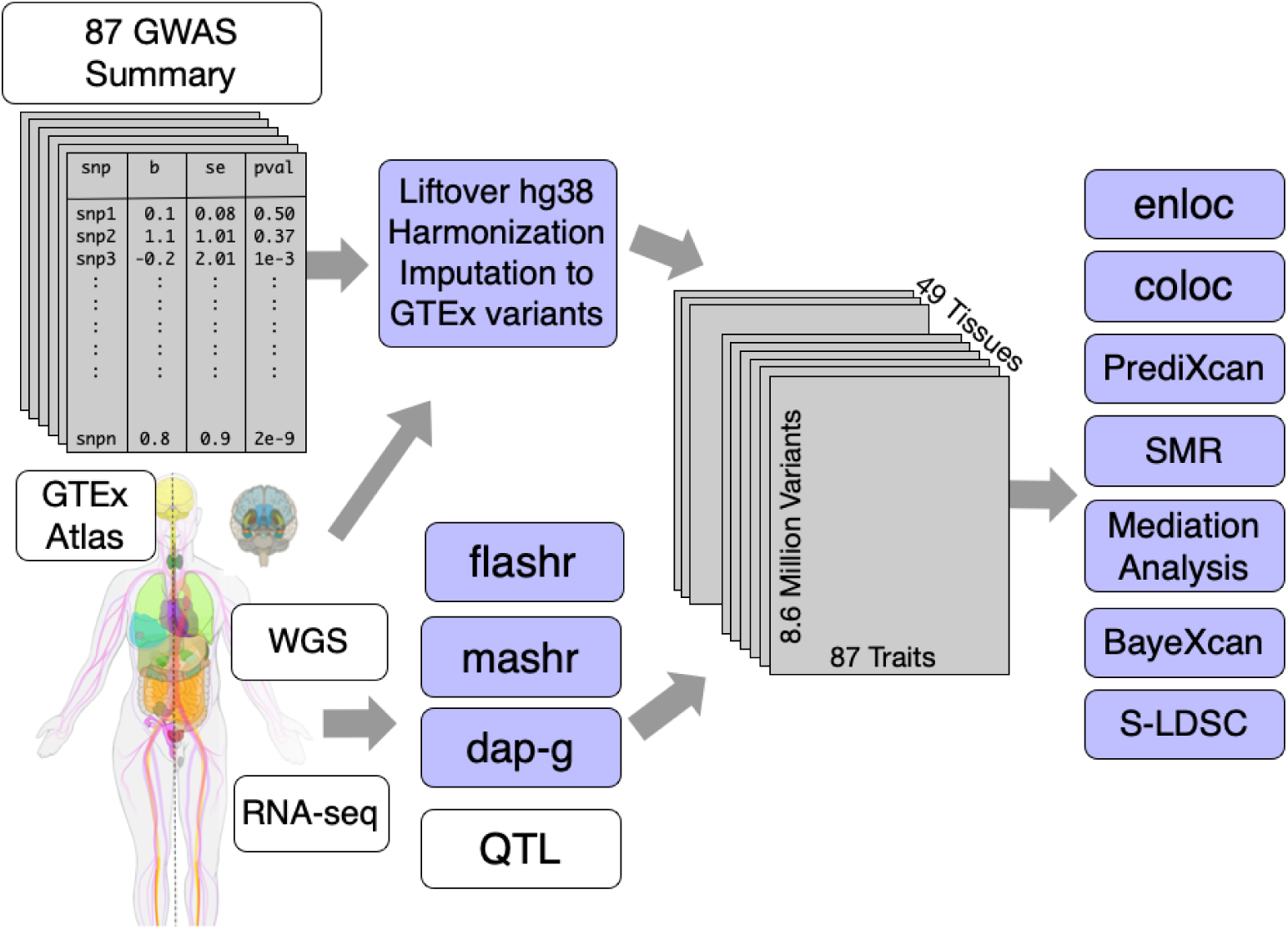
Overview of workflow for mapping complex trait associated QTLs. Full variant association summary statistics results from 114 GWAS were downloaded, standardized, and imputed to the GTEx v8 WGS variant calls (maf>0.01) for analyses. A total of 8.87 million imputed and genotyped variants were investigated to identify trait-associated QTLs. A total of 49 tissues, 87 studies (74 distinct traits), and 23,268 protein-coding genes and lncRNAs remained after stringent quality assurance protocols and selection criteria. A wide array of complex trait classes, including cardiometabolic, anthropometric, and psychiatric traits, were included.

We downloaded and processed 114 publicly available GWAS datasets with genome-wide variant association summary statistics (here onwards, summary statistics). After data harmonization, format standardization, missing data imputation and other quality assurance steps (fig. S2, fig. S3, and fig. S4), we retained 87 datasets representing 74 distinct complex traits including cardiometabolic, hematologic, neuropsychiatric and anthropometric traits (fig. S1). We provide the full list of datasets used in our study and all processing scripts as a resource to the community (table S2; see URLs).

Using these resources, we sought to identify likely causal associations among these gene- and alternatively spliced transcript-associated variants (eVariants and sVariants, respectively). For this purpose, we applied colocalization, enrichment, and association analyses, and provide a resource to enable investigations into gene prioritization approaches for disease associations (see URLs).

Gene expression and alternative splicing dysregulations have been proposed as the underlying mechanism of the association signals in many diseases [Pashos et al., 2017; Takata et al., 2017; Saferali et al., 2019; Li et al., 2016; Gamazon et al., 2018; Barbeira et al., 2018]. Similar to previous reports [GTEx Consortium et al., 2017], we observed robust and widespread enrichment of eQTLs and sQTLs among disease-associated variants (fig. S5). This observation suggests a causal role for expression and splicing regulation in complex traits.

### Dose-dependent regulatory effects of expression and alternative splicing on complex traits

Nevertheless, enrichment studies can be confounded by many unknown factors. Therefore, we sought to gather stronger evidence for a causal link by testing whether there is a dose-dependent effect of expression and splicing QTLs on complex traits. Fig. 2A illustrates schematically our approach. We examined whether expression or splicing associated variants (referred to as e/sVariants for the remainder of the paper) with higher impact on gene expression or splicing lead to higher impact on a complex trait, i.e. a larger GWAS effect (Fig. 2A). The impact of the regulation of a gene on a trait is quantified by the slope *β*_gene_. That is, a null hypothesis of no dose-dependent effect is equivalent to *β*_gene_ = 0.

**Fig. 2.**
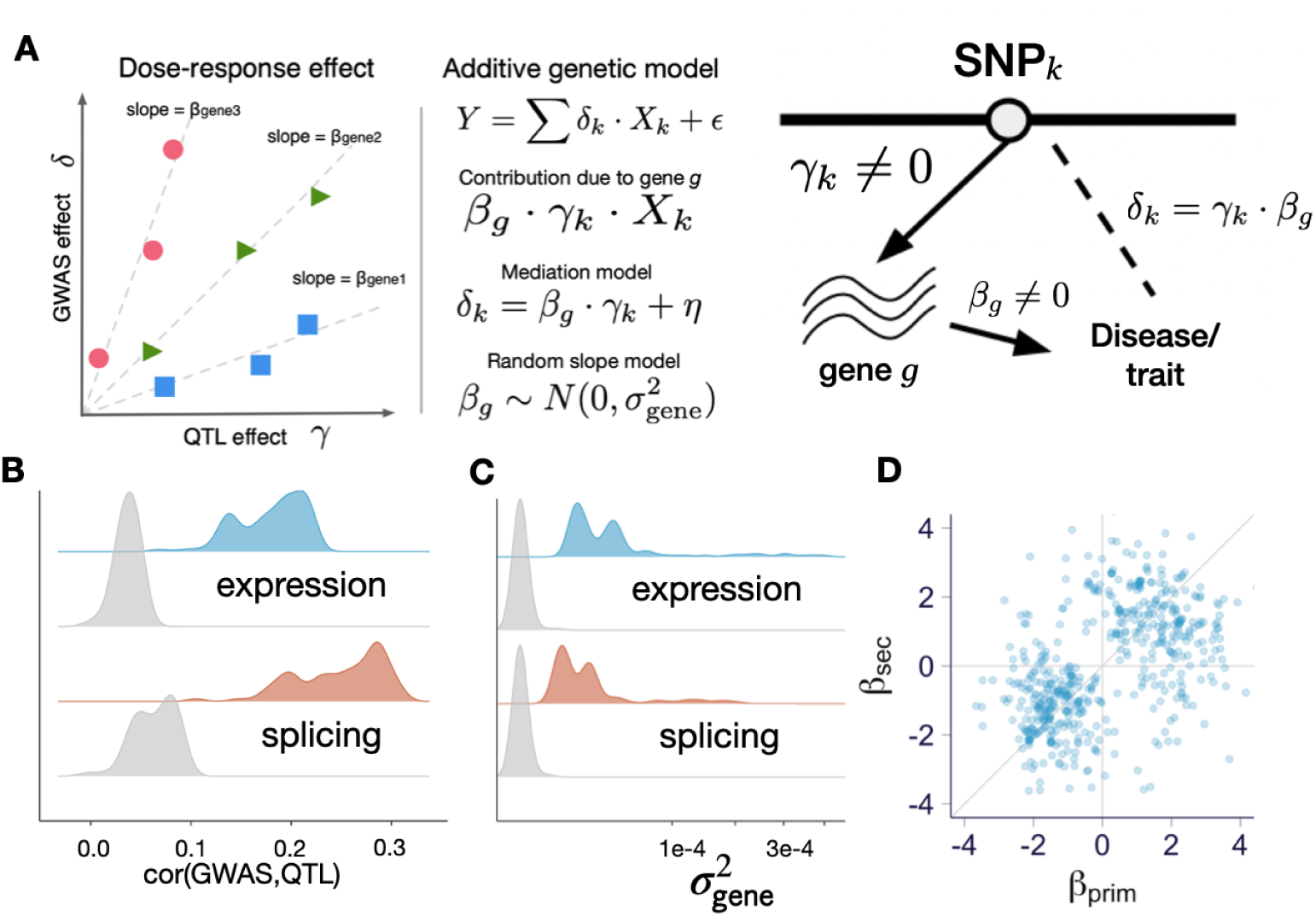
Dose-dependent effects of QTLs on complex traits. Here all analyses were performed with fine-mapped variants (QTL with highest posterior inclusion probability). **(A)** Schematic representation of dose-response model. **(B)** Correlation between QTL and GWAS effects, 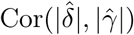. Gray distribution represents permuted null with matched local LD. Each data point corresponds to the median correlation for the trait across 49 tissues. **(C)** Average mediated effects from mediation model (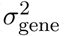, median across tissues). Gray distribution represents permuted null with matched local LD. **(E)** Mediated effects of secondary vs primary eQTLs of genes with colocalization probability (rcp) *>* 0.10. in whole blood, genes for all 87 traits are shown.

To reduce unnecessary noise in the analysis, we included only the most likely causal e/sVariant within each credible set as determined by the e/sQTL fine-mapping (denoted “fine-mapped variants” throughout the remainder of the paper. See Methods on QTL fine-mapping).

First, we quantified dose-dependent effect of expression and splicing regulation on the trait as the average mediating effect size,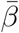. We calculated this average effect using the Pearson correlation between the absolute values of the molecular and complex trait effect sizes (cor(|*γ*|, |*δ*|)) across all fine-mapped variants (for any gene) for each trait-tissue pair. As hypothesized, we found, consistently across all tissue-trait pairs, a positive correlation between the GWAS and QTL effects, which was significantly larger than the permuted null with matched local LD. The average correlations were 0.18 (s.e. = 0.004, *p <* 1 *×* 10^−30^) and 0.25 (s.e. = 0.006, *p <* 1 *×* 10^−30^) for expression and splicing, respectively with the distribution of the median correlation across tissues for each trait shown in Fig. 2B. Averages and standard errors were calculated taking into account correlation between tissues, and p-values were calculated against permuted null with matched local LD (Supplementary Text). These results provide the first line of evidence of the dose-response effect.

To test and account for mediation effect heterogeneity (different slope/dosage sensitivity for different genes), we modeled the gene-specific mediation effect, *β*_*g*_, as a random variable following a normal distribution 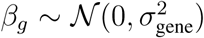. Under this random effects model, the null hypothesis can be stated as 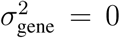 (Supplementary Text; Fig. 2C). As shown in Fig. 2C, these effects were significantly larger than expected from the permuted null (expression *p* = 1.8 *×* 10^−9^; splicing *p* = 2.5 *×* 10^−7^). These results indicate that strong genetic effects on expression or splicing are more likely to have a strong association to complex traits, adding strong support to a dose-dependent relationship between gene regulation and downstream traits.

Importantly, by averaging across all genes, the estimates, from both the average and the random-effects approach, of the mediating effect are robust to confounding due to LD, as discussed in the Supplementary Text.

Another way to account for mediation effect heterogeneity is to make use of the allelic series of independent eQTLs identified for over half of the eGenes [Aguet et al., 2019]. We examined whether the mediating effect (*β* = *δ/γ*) inferred from the primary eQTL (*β*_prim_) was consistent with the one inferred from the secondary eQTL (*β*_sec_). Among the independent eQTLs for a given gene, we called primary the one with the larger effect size. We considered only fine-mapped eQTLs given the low power to detect multiple independent sQTLs. We confirmed this concordance, as reported by the GTEx consortium [Aguet et al., 2019], demonstrating that the correlation between the primary and secondary mediating effects is larger than expected given the LD between them. To better visualize this concordance, we plotted the estimated mediating effects of primary against the secondary eQTLs (whole blood shown here but other tissues look similar, Fig. 2D and showed that they cluster near the identity line. All gene-trait pairs with relatively high regional colocalization probability (rcp>0.10, see colocalization details below) are shown here to facilitate visualization, but the clustering around the diagonal line was observed even without the filtering. This provides a third confirmatory evidence for the widespread dose-dependent effects of eQTLs on complex traits.

### Causal gene prediction and prioritization

In addition to genome-wide analyses that shed light on the molecular architecture of complex traits, QTL analysis of GWAS data can identify potential causal genes and molecular changes in individual GWAS loci. Towards this end, we performed association analysis with genetically predicted regulation and colocalization (Fig. 3A). After evaluating the performance of *coloc* and *enloc* [Wen et al., 2017; Giambartolomei et al., 2014], we chose *enloc* as our primary approach, due to its use of hierarchical models to estimate colocalization priors [Wen et al., 2017] and its ability to account for multiple causal variants. The *coloc* assumption of a single causal variant drastically reduces performance especially in large QTL datasets such as GTEx with widespread allelic heterogeneity (fig. S22). We estimated the posterior regional colocalization probability (rcp), using *enloc*, for 12,072,964 tissue-gene-GWAS locus-trait tuples and 67,943,800 tissue-splicing event-GWAS locus-trait tuples. For the tally of colocalized genes, we used rcp>0.5 as a stringent cutoff.

**Fig. 3.**
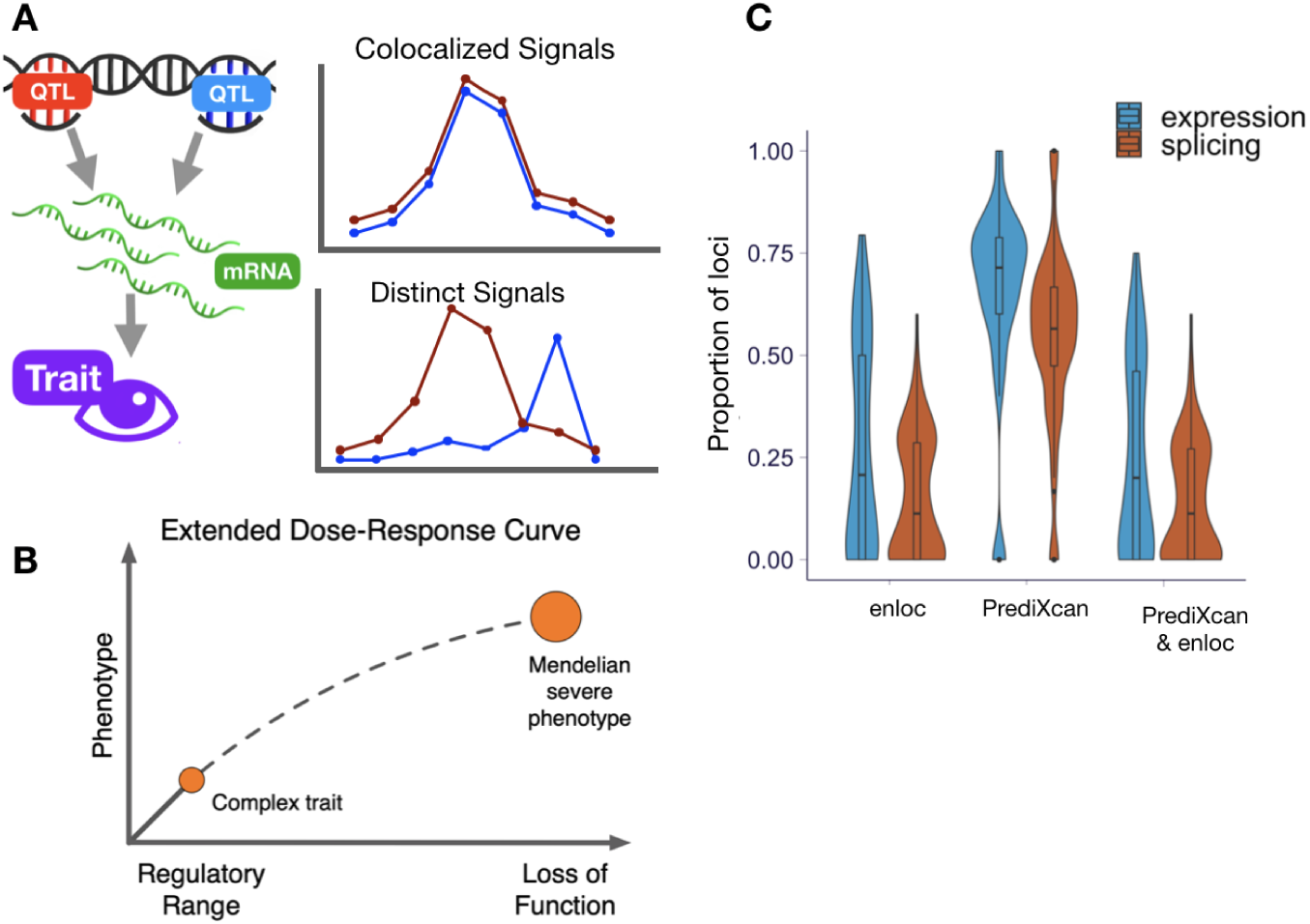
Identifying and validating predicted causal genes. **(A)** Schematic representation of association and colocalization approaches. **(B)** Schematic representation of extrapolating the dose-response curve to the Mendelian end of phenotypic variation spectrum [Plenge et al., 2013]. **(C)** Proportion of GWAS-associated loci per trait that contain colocalized and PrediXcan-associated signals for expression and splicing.

In total, we identified 3,477 (15% of 23,963) unique genes colocalizing with GWAS hits (rcp > 0.5) across all traits and tissues analyzed (fig. S9A). Similarly, 3,157 splicing events (1% out of 310,042) colocalized with GWAS hits, corresponding to 1,226 genes with at least one colocalized splicing event (5% of 23,963, fig. S9B).

Colocalization of e/sQTLs with GWAS variants provides important causal support for molecular traits. However, we found their estimates to be overly conservative. To illustrate this point, we tested the colocalization of height with itself, using two large-scale studies of individuals of European-ancestry individuals: GIANT [Wood et al., 2014] and UK Biobank. We started by performing fine-mapping of both GWAS results using *susier* [Wang et al., 2018]. Notably, only 416 (39%) of GIANT’s fine-mapped credible sets overlapped with the corresponding UK Biobank credible sets. We estimated the colocalization probability as the sum of the product of posterior inclusion probabilities of variants for each of the 1069 independent credible sets in GIANT, which is similar to the approach used by eCAVIAR [Hormozdiari et al., 2016]. Two thirds of the GIANT credible sets (66.2%) had a colocalization probability below 0.01 and about half (48.9%) had a colocalization probability below 0.001. In other words, two thirds of the loci found by GIANT would be considered not to be colocalized with UK Biobank’s loci when using a seemingly very loose colocalization probability cutoff of 0.01. Given the larger sample size of the UK Biobank GWAS (n=337,119 UKB GWAS vs. n=253,288 for GIANT), the low colocalization cannot be attributed to lack of power. This result is likely due in part to the sensitivity to small LD differences between different EUR populations that make up large GWAS meta-analysis cohorts such as GIANT. Our analysis illustrates the fact that colocalization probability estimates are highly conservative and may miss many causal genes, and low colocalization probability should not be interpreted as evidence of lack of a causal link between the molecular phenotype and the GWAS trait.

A complementary approach to colocalization is to estimate the GWAS trait association with genetically predicted gene expression or splicing [Gamazon et al., 2015]. The GTEx v8 data provides an important expansion of these analyses, allowing generation of prediction models in 49 tissues with whole genome sequencing data to impute gene expression and splicing variation. We trained prediction models using a variety of approaches and selected the top performing one based on precision, recall, and other metrics [Barbeira et al., 2020]. Briefly, the optimal model uses fine-mapping probabilities for feature selection and exploits global patterns of tissue sharing of regulation (Supplementary Text; fig. S23) to improve prediction. Multi-SNP prediction models were generated for a total of 686,241 gene-tissue and 1,816,703 splicing event-tissue pairs. The larger sample size and improved models led to an increase in the number of expression models to a median across tissues of 14,062, from a median of 4,776 GTEx v7 Elastic Net models (median increase at 191%, fig. S8). Splicing models are available only for the v8 release.

Next, we computed the association between an imputed molecular phenotype (expression or splicing) and a trait to estimate the genic effect on the trait, using the summary statistics based PrediXcan [Barbeira et al., 2018]. Given the widespread tissue-sharing of regulatory variation [GTEx Consortium et al., 2017], we also computed MultiXcan scores to integrate patterns of associations from multiple tissues and increase statistical power [Barbeira et al., 2019]. Out of the 22,518 genes tested with PrediXcan, 6,407 (28%) showed a significant association with at least one of the 87 traits at Bonferroni-corrected p-value threshold (*p <* 0.05*/*686, 241, where the denominator is the number of gene-tissue pairs tested; fig. S9). For splicing, about 15% (20,364 of 138,890) of tested splicing events showed a significant association (*p <* 0.05*/*1, 816, 703, where the denominator is the number of intron-tissue pairs tested). Nearly all traits (94%; 82 out of 87) showed at least one significant gene-level PrediXcan association in at least one tissue (fig. S14 and S15); the median number of associated genes across traits was 974. This resource of PrediXcan associations can be used to prioritize a list of putatively causal genes for follow-up studies.

To replicate the PrediXcan expression associations in an independent dataset, BioVU, which is a large-scale biobank tied to Electronic Health Records [Roden et al., 2008; Denny et al., 2013], we selected seven traits with predicted high statistical power. Out of 947 gene-tissue-trait discoveries tested, 458 unique gene-tissue-trait triplets (48%) showed replication in this independent biobank (PrediXcan association p < 0.05; see Supplementary Text).

Altogether, these results provide abundant links between gene regulation and GWAS loci. To further quantify this, we split the genome into approximately LD-independent blocks [Berisa and Pickrell, 2016] and identified blocks with a significant GWAS variant for each trait (at Bonferroni threshold adjusted for number of variants 0.05*/*8.8 *×* 10^6^ ∼ 5.7 *×* 10^−9^); we refer to any such region-trait pair by “GWAS locus”. We calculated the proportion of GWAS loci that contain a significantly associated gene via PrediXcan or a colocalized gene via *enloc* (rcp > 0.5). Briefly, the LD blocks are defined by analyzing empirical patterns of LD observed in 1000 Genomes [**?**] and variants in different regions are unlikely to be correlated, thus providing us with a data-driven criterion to distinguish independent genomic signals.

Across the traits, 72% (3,899/5,385) of GWAS loci had a PrediXcan expression association in the same LD block, of which 55% (2,125/3,899) had evidence of colocalization with an eQTL (table S4). For splicing, 62% (3,345/5,385) had a PrediXcan association of which 34% (1,135/3,345) colocalized with an sQTL (fig. S13). From the combined list of eGenes and sGenes, 47% of loci have a gene with both *enloc* and PrediXcan support. The distribution of the proportion of associated and colocalized GWAS loci across 87 traits is summarized in Fig. 3-C; for a typical complex trait, about 20% of GWAS loci contained a colocalized, significantly associated gene while 11% contained a colocalized, significantly associated splicing event. These results propose function for a large number of GWAS loci, but most loci remain without candidate genes, highlighting the need to expand the resolution of transcriptome studies.

Of note, two members of the sterolin family, *ABCG5* and *ABCG8*, showed highly significant predicted causal associations using both PrediXcan and *enloc* for LDL-C levels and self-reported high cholesterol levels. *ABCG8* showed more significant associations in both datasets (chr2: 43838964 - 43878466; UKB self-reported high cholesterol: -log10(*p*_PrediXcan_) = 38.43, rcp = 0.985; GLGC LDL-C: -log10(*p*_PrediXcan_) = 71.40, rcp = 0.789), compared to *ABCG5* (chr2: 43812472 - 43838865; -log10(*p*_PrediXcan_) = 36.85, rcp = 0.941; -log10(*p*_PrediXcan_) = 80.80, rcp = 0.705). Mutations in either of the two ATP-binding cassette (ABC) half-transporters, *ABCG5* and *ABCG8*, lead to reduced secretion of sterols into bile, and ultimately, obstruct cholesterol and other sterols exiting the body [Kidambi and Patel, 2008]. In mice with disrupted *Abcg5* and *Abcg8* (G5G8-/-), a 2- to 3-fold increase in the fractional absoprtion of dietary plan steols and extrememly low biliary cholesterol levels was observed, indicating that disrupting these genes contribute greatly to plasma cholesterol levels [Yu et al., 2002]. The overexpression of human *ABCG5* and *ABCG8* in transgenic Ldlr-/- mice resulted in 30% reduction in hepatic cholesterol levels and 70% reduced atherosclerotic legion in the aortic root and arch [Wilund et al., 2004] after 6-months on a Western diet.

Several other lipid-associated loci were also consistently predicted as causal across OMIM, the rare variant derived set, PrediXcan and *enloc*. Rare protein-truncating variants in *APOB* have been previously associated with reduced LDL-C and triglyceride levels and reduced coronary heart disease risk [Peloso et al., 2019]. Interestingly, *APOB* has been predicted as a causal gene in four related traits, coronary artery disease, LDL-C levels, triglyceride levels, and self-reported high cholesterol levels. Among the four traits, PrediXcan showed the highest association to LDL-C levels (-log10(*p*_PrediXcan_) = 130.89; rcp = 0.485) while self-reported high cholesterol showed the strongest evidence using *enloc* at nearly maximum posterior probability (-log10(*p*_PrediXcan_) = 93.66; rcp = 0.969). Although *APOB* has been suggested as a better molecular indicator of predicted cardiac events in place of LDL-C levels [Walldius and Jungner, 2004; Contois et al., 2009], its translation has been surprisingly slow in clinical practice [Leslie, 2017]. Here, we provide an additional support for the crucial role *APOB* may play in predicting lipid traits.

### Performance for identifying “ground truth” genes

To compare the ability of different approaches to identify the causal gene that mediates the association between GWAS loci and the traits, we sought to curate sets of “ground truth” genes using information that is independent of GWAS results. We call these sets “silver standards” as a reminder of their imperfect nature. The first silver standard was based on the OMIM (Online Mendelian Inheritance in Man) database [Hamosh et al., 2005] and the second one was based on publicly available rare variant tests from exome-wide association studies [Marouli et al., 2017; Liu et al., 2017; Locke et al., 2019] (fig. S16, table S6), resulting in 1,592 OMIM gene-trait pairs and 101 rare variant based gene-trait pairs (table S11, table S12, fig. S17.)

The rationale behind the choice of the OMIM database is the comorbidity among Mendelian and complex diseases suggesting that genes whose loss of function cause Mendelian diseases also manifest in milder phenotypic variation when modified to a lesser degree by regulatory variation [Lupski et al., 2011; Blair et al., 2013]. In other words, that the dose-response curve at the regulatory range may be extrapolated to the rare, loss-of-function end (Fig. 3B). The rationale behind the use of the rare variant association study results is the excess of deleterious rare variants associated with complex traits in genes that are in the vicinity of common variants associated with the same trait [Marouli et al., 2017; Fuchsberger et al., 2016; Keinan and Clark, 2012]. Note that rare variant associations are nearly independent of common variants due to the allele frequency difference between them.

For the analysis, we partitioned the genome into approximately independent LD blocks [Berisa and Pickrell, 2016] and considered all the blocks where a silver standard gene was available for the trait. Since only genes in the vicinity of an index gene can be discovered with cis-regulatory information, we only considered the LD blocks with a GWAS significant variant. This selection resulted in 228 OMIM gene-trait pairs (28 distinct traits) and 80 rare variant-associated genes-trait pairs (5 distinct traits) that are located within the same LD block as the GWAS locus for a matched trait (fig. S18).

Both PrediXcan and *enloc* based on expression and splicing showed good sensitivity and specificity for identifying the silver standard genes as demonstrated by the ROC curves in Figs. 4A-B. These are well above the gray random guess lines indicating the predictive ability of these methods to find causal genes.

For applications such as target selection for drug development or follow-up experiments, another relevant metric is the precision or, equivalently, positive predictive value (PPV) – the probability that the gene-trait link is causal given that it is called significant or colocalized. Precision recall curves for expression and splicing based predictions are shown in Fig. 4C-D. With more stringent threshold (towards the left in the recall axis), higher precision is obtained.

**Fig. 4.**
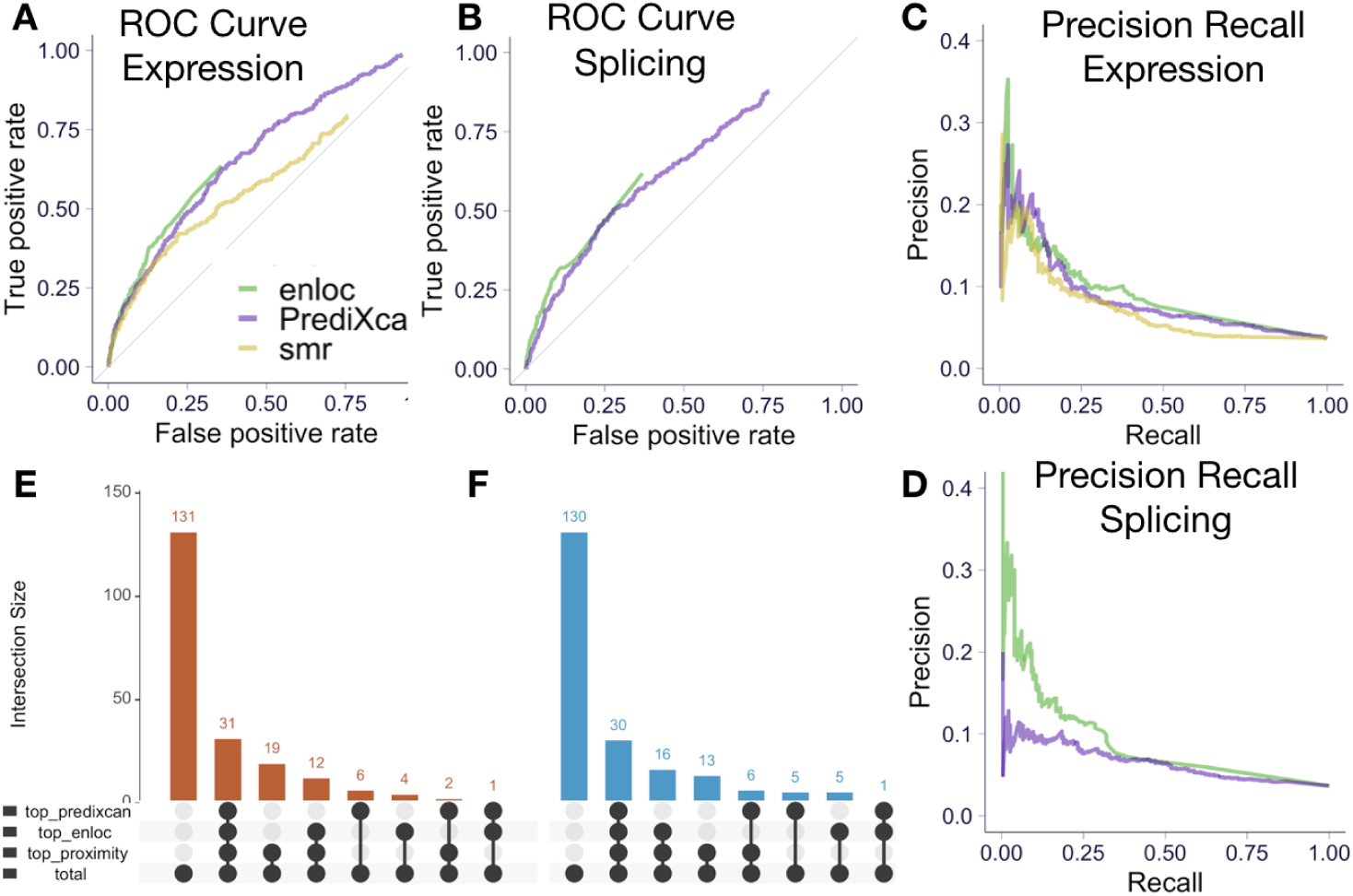
Causal gene identification performance. ROC curves of *enloc* and PrediXcan statistics to identify the ‘causal’ genes (OMIM silver standard) using expression **(A)** and splicing **(B)** are shown. Precision recall curves of *enloc* and PrediXcan to identify silver standard genes using expression **(C)** and splicing **(D)** (We show the precision in the range 0 to 0.4 to improve visualization). The number of GWAS loci (LD block-trait pairs) where the OMIM gene was ranked at the top by proximity, *enloc*, and PrediXcan using expression **(E)** and splicing **(F)**. In 131 loci out of 206 the OMIM gene was not ranked at the top by either proximity, significance, or colocalization. In thirty one of the loci, the OMIM gene was ranked first by all three criteria. In nineteen loci, the OMIM gene was closest gene (to the top GWAS variant) but not the top gene by PrediXcan significance nor *enloc*’s colocalization proability.

For example, 8.7% of genes with PrediXcan significant genes (*p <* 0.05*/*49 *×* number of gene/trait pairs) were OMIM genes and 14.8% of genes with high colocalization probability (rcp>0.5) were also OMIM genes for matched traits.

Multiple factors contribute to the rather low precision. One of them is the widespread molecular pleiotropy [Aguet et al., 2019], i.e. multiple genes affected by the same trait-associated variants. Another factor reducing the overall causal gene detection performance is the inherent bias of the OMIM gene list. Our current understanding of gene function is biased towards protein-coding variants with very large effects, as reflected in the list of OMIM genes. Genes associated to rare severe disease tend to be depleted of regulatory variation [Karczewski et al., 2019; Mohammadi et al., 2019], which will decrease the performance of a QTL-based method in a way that is unlikely to be generally applicable to GWAS genes that are more tolerant to regulatory variation [Mohammadi et al., 2019].

Among the 206 loci with at least one OMIM gene (a few loci contained multiple OMIM genes), an OMIM gene was the closest to the top GWAS SNP in 31.6% of the loci, it was the most colocalized in 24.8% of the loci, and it was the most significant in 20.4% of the loci (Fig. 4E-F).

To further investigate whether this predictive power could be improved by combining multiple criteria, we performed a joint logistic regression of OMIM gene status on 1) the proximity of the top GWAS variant to the nearest gene (distance to the gene body), 2) posterior probability of colocalization, and 3) PrediXcan association significance between QTL and GWAS variants. To make the scale of the three features more comparable, we used their respective ranking. When genes did not have an enloc or PrediXcan score, they were assigned to the last position in the ranking. All three features were significant predictors of OMIM gene status, with better ranked genes more likely to be OMIM genes (proximity *p* = 2.0 *×* 10^−2^, *enloc p* = 6.1 *×* 10^−3^, PrediXcan *p* = 2.5 *×* 10^−4^), indicating that each method provides an additional source of causal evidence even after conditioning on the others. Similar results were obtained using splicing colocalization and association scores and the rare variant based silver standard, as shown in table S8. These results provide further empirical evidence that a combination of colocalization and association methods will perform better than individual ones. The significance of the proximity score even after accounting for significance and colocalization indicates missing regulatory events, i.e. mechanisms that may be uncovered by that assays other tissue or cell type contexts, larger samples, and other molecular traits, underscoring the need to expand the size and breadth of QTL studies.

Predicted OMIM genes included well-known findings such as *PCSK9* for LDLR, with *PCSK9* significant and colocalized for relevant GWAS traits (LDL-C levels, coronary artery disease, and self-reported high cholesterol), and *Interleukins* and *HLA* subunits for asthma, both significant and colocalized for related immunological traits. Significantly associated and colocalized genes that predicted OMIM genes also included *FLG* (eczema), *TPO* (hypothyroidism), and *NOD2* (inflammatory bowel disease) (see table S11 for complete list). Analysis with rare variant-based silver standard yielded similar conclusions (Supplementary Text; fig. S21).

### Tissue enrichment of GWAS signals

A systematic survey of regulatory variation across 49 human tissues promises to facilitate the identification of the tissues of action for complex traits. However, because of the broad sharing of regulatory variation across tissues and the reduced significance of tissue-specific eQTLs, causal tissue identification has been challenging. Here we used sparse factors from *FLASH* Urbut et al. [2018] representing patterns of tissue sharing of eQTLs (Supplementary Text), to classify each gene-trait association into one of 15 tissue classes (fig. S23). Using the pattern of tissue classes of non-colocalized genes (rcp = 0) as the expected null, we assessed whether significantly associated and colocalized genes (PrediXcan significant and rcp > 0.01) were over-represented in certain tissue classes (Fig. 5). Consistent with previous reports [Gamazon et al., 2018; Ongen et al., 2017], we identified several instances in which the most significant tissue is supported by current biological knowledge. For example, blood cell count traits were enriched in whole blood, neuroticism and fluid intelligence in brain/pituitary, hypothyrodism in thyroid, coronary artery disease in artery, and cholesterol-related traits in liver. Taken together, these results show the potential of leveraging regulatory variation to help identify tissues of relevance for complex traits.

**Fig. 5.**
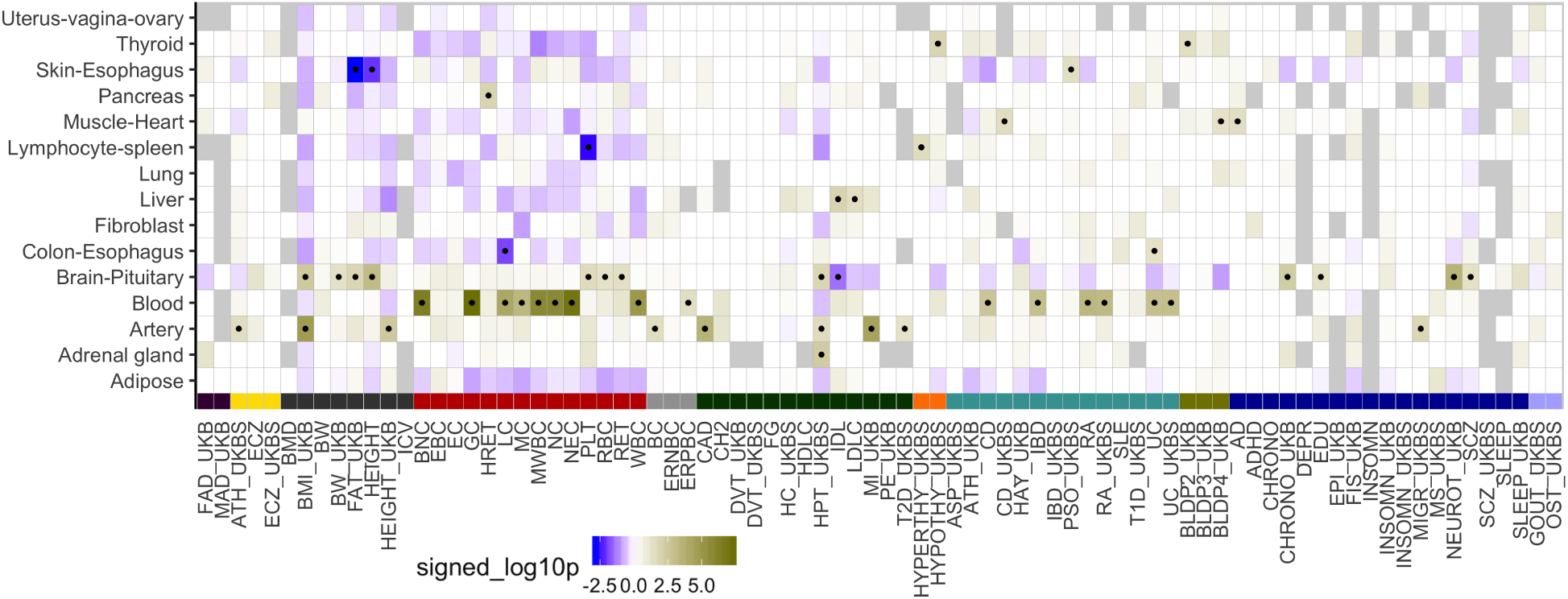
Identifying trait-relevant tissues using tissue-specific enrichment. Enrichment of tisssue-specific association and colocalization compared to the pattern of tissue-specificity of non-colocalized genes. Over-representation of the tissue class for PrediXcan-significant and colocalized genes is indicated by dark yellow while depletion is indicated by blue. Black dots label the tissue class-trait pairs passing the nominal p-value significance threshold of 0.05. Abbreviation: S2. Trait category colors: S1.

## Discussion

We performed in-depth examination of the phenotypic consequences of the genetic regulation of the transcriptome and provide data-driven analytical approaches to benchmark methods that assign function to GWAS loci and best-practice guidelines for using the GTEx resources to interpret GWAS results. We provide a systematic empirical demonstration of the widespread dose-dependent effect of expression and splicing on complex traits, i.e., variants with larger impact at the molecular level have larger impact at the trait level. Furthermore, we found that target genes in GWAS loci identified by *enloc* and PrediXcan were predictive of OMIM genes for matched traits, implying that for a proportion of the genes, the dose-response curve can be extrapolated to the rare and more severe end of the genotype-trait spectrum. The observation that common regulatory variants target genes also implicated by rare coding variants underscores the extent to which these different types of genetic variants converge to mediate a spectrum of similar pathophysiological effects and may provide a powerful approach to drug target discovery.

We implemented association and colocalization methods that leverage the observed allelic heterogeneity of expression traits. After extensive comparison using two independent sets of silver standard gene-trait pairs, we conclude that combining *enloc*, PrediXcan, and proximity ranking outperforms the individual approaches. The significance of the proximity ranking is a sign of the “missing regulability” emphasizing the need to expand the resolution, sample size, and range of contexts of transcriptome studies as well as to examine other molecular mechanisms.

We caution that the increased power offered by this release of the GTEx resources also brings higher risk of false links due to LD contamination and that naive use of eQTL or sQTL association p-values to assign function to a GWAS locus can be misleading. Colocalization approaches can be used to weed out LD contamination but given the lack of LD references from source studies, they can also be overtly conservative. General purpose reference LD from publicly available sources are not sufficient for fine-mapping and colocalization approaches, which can be highly sensitive to LD misspecification when only summary results are used [Benner et al., 2017]. The GWAS community has made great progress in recognizing the need to share summary results, but to take full advantage of these data, improved sharing of LD information from the source study as well as from large sequencing reference datasets, is also required.

Finally, we generated several resources that can open the door for addressing key questions in complex trait genomics. We present a catalog of gene-level associations, including potential target genes for nearly half of the GWAS loci investigated here that provides a rich basis for studies on the functional mechanisms of complex diseases and traits. We provide a database of optimal gene expression imputation models that were built on the fine-mapping probabilities for feature selection and that leverage the global patterns of tissue sharing of regulation to improve the weights. These imputation models of expression and splicing, which to date has been challenging to study, provide a foundation for transcriptome-wide association studies of the human phenome – the collection of all human diseases and traits – to further accelerate discovery of trait-associated genes. Collectively, these data thus represent a valuable resource, enabling novel biological insights and facilitating follow-up studies of causal mechanisms.

## Supporting information

Large Supplementary Tables

## Authors

^∗^ alphabetic order

**Lead Analysts**^∗^ *Equal contribution* Alvaro N Barbeira, Rodrigo Bonazzola, Eric R Gamazon, Yanyu Liang, YoSon Park

**Analysts**^∗^ François Aguet, Lisa Bastarache, Ron Do, Gao Wang, Andrew R Hamel, Farhad Hormozdiari, Zhuoxun Jiang, Daniel Jordan, Sarah Kim-Hellmuth, Boxiang Liu, Milton D Pividori, Abhiram Rao, Marie Verbanck, Dan Zhou

**GTEx GWAS Working Group**^∗^ François Aguet, Kristin Ardlie, Alvaro N Barbeira, Rodrigo Bonazzola, Christopher D Brown, Lin Chen, Eric R Gamazon, Kevin Gleason, Andrew R Hamel, Farhad Hormozdiari, Hae Kyung Im, Sarah Kim-Hellmuth, Tuuli Lappalainen, Yanyu Liang, Boxiang Liu, Dan L Nicolae, Yoson Park, Milton D Pividori, Abhiram Rao, John M. Rouhana, Ayellet V Segrè, Xiaoquan Wen

**Senior Leadership**^∗^ Kristin Ardlie, Christopher D. Brown, Hae Kyung Im, Tuuli Lappalainen, Mark McCarthy, Stephen Montgomery, Ayellet V Segrè, Matthew Stephens, Xiaoquan Wen

**Manuscript Writing Group**^∗^ Eric R Gamazon, Hae Kyung Im, Tuuli Lappalainen, Yanyu Liang, YoSon Park

**Corresponding Author**^∗^ Hae Kyung Im

## GTEx Consortium

### Laboratory and Data Analysis Coordinating Center (LDACC)

François Aguet^1^, Shankara Anand^1^, Kristin G Ardlie^1^, Stacey Gabriel^1^, Gad Getz^1,2^, Aaron Graubert^1^, Kane Hadley^1^, Robert E Handsaker^3,4,5^, Katherine H Huang^1^, Seva Kashin^3,4,5^, Xiao Li^1^, Daniel G MacArthur^4,6^, Samuel R Meier^1^, Jared L Nedzel^1^, Duyen Y Nguyen^1^, Ayellet V Segrè^1,7^, Ellen Todres^1^

### Analysis Working Group (funded by GTEx project grants)

François Aguet^1^, Shankara Anand^1^, Kristin G Ardlie^1^, Brunilda Balliu^8^, Alvaro N Barbeira^9^, Alexis Battle^10,11^, Rodrigo Bonazzola^9^, Andrew Brown^12,13^, Christopher D Brown^14^, Stephane E Castel^15,16^, Don Conrad^17,18^, Daniel J Cotter^19^, Nancy Cox^20^, Sayantan Das^21^, Olivia M de Goede^19^, Emmanouil T Dermitzakis^12,22,23^, Barbara E Engelhardt^24,25^, Eleazar Eskin^26^, Tiffany Y Eulalio^27^, Nicole M Ferraro^27^, Elise Flynn^15,16^, Laure Fresard^28^, Eric R Gamazon^29,30,31,20^, Diego Garrido-Martín^32^, Nicole R Gay^19^, Gad Getz^1,2^, Aaron Graubert^1^, Roderic Guigó^32,33^, Kane Hadley^1^, Andrew R Hamel^7,1^, Robert E Handsaker^3,4,5^, Yuan He^10^, Paul J Hoffman^15^, Farhad Hormozdiari^34,1^, Lei Hou^35,1^, Katherine H Huang^1^, Hae Kyung Im^9^, Brian Jo^24,25^, Silva Kasela^15,16^, Seva Kashin^3,4,5^, Manolis Kellis^35,1^, Sarah Kim-Hellmuth^15,16,36^, Alan Kwong^21^, Tuuli Lappalainen^15,16^, Xiao Li^1^, Xin Li^28^, Yanyu Liang^9^, Daniel G MacArthur^4,6^, Serghei Mangul^26,37^, Samuel R Meier^1^, Pejman Mohammadi^15,16,38,39^, Stephen B Montgomery^28,19^, Manuel Muñoz-Aguirre^32,40^, Daniel C Nachun^28^, Jared L Nedzel^1^, Duyen Y Nguyen^1^, Andrew B Nobel^41^, Meritxell Oliva^9,42^, YoSon Park^14,43^, Yongjin Park^35,1^, Princy Parsana^11^, Ferran Reverter^44^, John M Rouhana^7,1^, Chiara Sabatti^45^, Ashis Saha^11^, Ayellet V Segrè^1,7^, Andrew D Skol^9,46^, Matthew Stephens^47^, Barbara E Stranger^9,48^, Benjamin J Strober^10^, Nicole A Teran^28^, Ellen Todres^1^, Ana Viñuela^49,12,22,23^, Gao Wang^47^, Xiaoquan Wen^21^, Fred Wright^50^, Valentin Wucher^32^, Yuxin Zou^51^

### Analysis Working Group (not funded by GTEx project grants)

Pedro G Ferreira^52,53,54^, Gen Li^55^, Marta Melé^56^, Esti Yeger-Lotem^57,58^

### Leidos Biomedical - Project Management

Mary E Barcus^59^, Debra Bradbury^60^, Tanya Krubit^60^, Jeffrey A McLean^60^, Liqun Qi^60^, Karna Robinson^60^, Nancy V Roche^60^, Anna M Smith^60^, Leslie Sobin^60^, David E Tabor^60^, Anita Undale^60^

### Biospecimen collection source sites

Jason Bridge^61^, Lori E Brigham^62^, Barbara A Foster^63^, Bryan M Gillard^63^, Richard Hasz^64^, Marcus Hunter^65^, Christopher Johns^66^, Mark Johnson^67^, Ellen Karasik^63^, Gene Kopen^68^, William F Leinweber^68^, Alisa McDonald^68^, Michael T Moser^63^, Kevin Myer^65^, Kimberley D Ramsey^63^, Brian Roe^65^, Saboor Shad^68^, Jeffrey A Thomas^68,67^, Gary Walters^67^, Michael Washington^67^, Joseph Wheeler^66^

### Biospecimen core resource

Scott D Jewell^69^, Daniel C Rohrer^69^, Dana R Valley^69^

### Brain bank repository

David A Davis^70^, Deborah C Mash^70^

### Pathology

Mary E Barcus^59^, Philip A Branton^71^, Leslie Sobin^60^

### ELSI study

Laura K Barker^72^, Heather M Gardiner^72^, Maghboeba Mosavel^73^, Laura A Siminoff^72^

### Genome Browser Data Integration & Visualization

Paul Flicek^74^, Maximilian Haeussler^75^, Thomas Juettemann^74^, W James Kentv^75^, Christopher M Lee^75^, Conner C Powell^75^, Kate R Rosenbloom^75^, Magali Ruffier^74^, Dan Sheppard^74^, Kieron Taylor^74^, Stephen J Trevanion^74^, Daniel R Zerbino^74^

### eGTEx groups

Nathan S Abell^19^, Joshua Akey^76^, Lin Chen^42^, Kathryn Demanelis^42^, Jennifer A Doherty^77^, Andrew P Feinberg^78^, Kasper D Hansen^79^, Peter F Hickey^80^, Lei Hou^35,1^, Farzana Jasmine^42^, Lihua Jiang^19^, Rajinder Kaul^81,82^, Manolis Kellis^35,1^, Muhammad G Kibriya^42^, Jin Billy Li^19^, Qin Li^19^, Shin Lin^83^, Sandra E Linder^19^, Stephen B Montgomery^28,19^, Meritxell Oliva^9,42^, Yongjin Park^35,1^, Brandon L Pierce^42^, Lindsay F Rizzardi^84^, Andrew D Skol^9,46^, Kevin S Smith^28^, Michael Snyder^19^, John Stamatoyannopoulos^81,85^, Barbara E Stranger^9,48^, Hua Tang^19^, Meng Wang^19^

### NIH program management

Philip A Branton^71^, Latarsha J Carithers^71,86^, Ping Guan^71^, Susan E Koester^87^, A. Roger Little^88^, Helen M Moore^71^, Concepcion R Nierras^89^, Abhi K Rao^71^, Jimmie B Vaught^71^, Simona Volpi^90^

### Affiliations

1. The Broad Institute of MIT and Harvard, Cambridge, MA, USA

2. Cancer Center and Department of Pathology, Massachusetts General Hospital, Boston, MA, USA

3. Department of Genetics, Harvard Medical School, Boston, MA, USA

4. Program in Medical and Population Genetics, The Broad Institute of Massachusetts Institute of Technology and Harvard University, Cambridge, MA, USA

5. Stanley Center for Psychiatric Research, Broad Institute, Cambridge, MA, USA

6. Analytic and Translational Genetics Unit, Massachusetts General Hospital, Boston, MA, USA

7. Ocular Genomics Institute, Massachusetts Eye and Ear, Harvard Medical School, Boston, MA, USA

8. Department of Biomathematics, University of California, Los Angeles, Los Angeles, CA, USA

9. Section of Genetic Medicine, Department of Medicine, The University of Chicago, Chicago, IL, USA v

10. Department of Biomedical Engineering, Johns Hopkins University, Baltimore, MD, USA

11. Department of Computer Science, Johns Hopkins University, Baltimore, MD, USA

12. Department of Genetic Medicine and Development, University of Geneva Medical School, Geneva, Switzerland

13. Population Health and Genomics, University of Dundee, Dundee, Scotland, UK

14. Department of Genetics, University of Pennsylvania, Perelman School of Medicine, Philadelphia, PA, USA

15. New York Genome Center, New York, NY, USA

16. Department of Systems Biology, Columbia University, New York, NY, USA

17. Department of Genetics, Washington University School of Medicine, St. Louis, Missouri, USA

18. Department of Pathology & Immunology, Washington University School of Medicine, St. Louis, Missouri, USA

19. Department of Genetics, Stanford University, Stanford, CA, USA

20. Division of Genetic Medicine, Department of Medicine, Vanderbilt University Medical Center, Nashville, TN, USA

21. Department of Biostatistics, University of Michigan, Ann Arbor, MI, USA

22. Institute for Genetics and Genomics in Geneva (iGE3), University of Geneva, Geneva, Switzerland

23. Swiss Institute of Bioinformatics, Geneva, Switzerland

24. Department of Computer Science, Princeton University, Princeton, NJ, USA

25. Center for Statistics and Machine Learning, Princeton University, Princeton, NJ, USA

26. Department of Computer Science, University of California, Los Angeles, Los Angeles, CA, USA

27. Program in Biomedical Informatics, Stanford University School of Medicine, Stanford, CA, USA

28. Department of Pathology, Stanford University, Stanford, CA, USA

29. Data Science Institute, Vanderbilt University, Nashville, TN, USA

30. Clare Hall, University of Cambridge, Cambridge, UK

31. MRC Epidemiology Unit, University of Cambridge, Cambridge, UK

32. Centre for Genomic Regulation (CRG), The Barcelona Institute for Science and Technology, Barcelona, Catalonia, Spain

33. Universitat Pompeu Fabra (UPF), Barcelona, Catalonia, Spain

34. Department of Epidemiology, Harvard T.H. Chan School of Public Health, Boston, MA, USA

35. Computer Science and Artificial Intelligence Laboratory, Massachusetts Institute of Technology, Cambridge, MA, USA

36. Statistical Genetics, Max Planck Institute of Psychiatry, Munich, Germany

37. Department of Clinical Pharmacy, School of Pharmacy, University of Southern California, Los Angeles, CA, USA

38. Scripps Research Translational Institute, La Jolla, CA, USA

39. Department of Integrative Structural and Computational Biology, The Scripps Research Institute, La Jolla, CA, USA

40. Department of Statistics and Operations Research, Universitat Politècnica de Catalunya (UPC), Barcelona, Catalonia, Spain

41. Department of Statistics and Operations Research and Department of Biostatistics, University of North Carolina, Chapel Hill, NC, USA

42. Department of Public Health Sciences, The University of Chicago, Chicago, IL, USA

43. Department of Systems Pharmacology and Translational Therapeutics, University of Pennsylvania, Perelman School of Medicine, Philadelphia, PA, USA

44. Department of Genetics, Microbiology and Statistics, University of Barcelona, Barcelona. Spain.

45. Departments of Biomedical Data Science and Statistics, Stanford University, Stanford, CA, USA

46. Department of Pathology and Laboratory Medicine, Ann & Robert H. Lurie Children’s Hospital of Chicago, Chicago, IL, USA

47. Department of Human Genetics, University of Chicago, Chicago, IL, USA

48. Center for Genetic Medicine, Department of Pharmacology, Northwestern University, Feinberg School of Medicine, Chicago, IL, USA

49. Department of Twin Research and Genetic Epidemiology, King’s College London, London, UK

50. Bioinformatics Research Center and Departments of Statistics and Biological Sciences, North Carolina State University, Raleigh, NC, USA

51. Department of Statistics, University of Chicago, Chicago, IL, USA

52. Department of Computer Sciences, Faculty of Sciences, University of Porto, Porto, Portugal

53. Instituto de Investigação e Inovação em Sauúde, Universidade do Porto, Porto, Portugal

54. Institute of Molecular Pathology and Immunology, University of Porto, Porto, Portugal

55. Columbia University Mailman School of Public Health, New York, NY, USA

56. Life Sciences Department, Barcelona Supercomputing Center, Barcelona, Spain

57. Department of Clinical Biochemistry and Pharmacology, Ben-Gurion University of the Negev, Beer-Sheva, Israel

58. National Institute for Biotechnology in the Negev, Beer-Sheva, Israel

59. Leidos Biomedical, Frederick, MD, USA

60. Leidos Biomedical, Rockville, MD, USA

61. UNYTS, Buffalo, NY, USA

62. Washington Regional Transplant Community, Annandale, VA, USA

63. Therapeutics, Roswell Park Comprehensive Cancer Center, Buffalo, NY, USA

64. Gift of Life Donor Program, Philadelphia, PA, USA

65. LifeGift, Houston, TX, USA

66. Center for Organ Recovery and Education, Pittsburgh, PA, USA

67. LifeNet Health, Virginia Beach, VA. USA v

68. National Disease Research Interchange, Philadelphia, PA, USA v

69. Van Andel Research Institute, Grand Rapids, MI, USA

70. Department of Neurology, University of Miami Miller School of Medicine, Miami, FL, USA

71. Biorepositories and Biospecimen Research Branch, Division of Cancer Treatment and Diagnosis, National Cancer Institute, Bethesda, MD, USA

72. Temple University, Philadelphia, PA, USA

73. Virgina Commonwealth University, Richmond, VA, USA

74. European Molecular Biology Laboratory, European Bioinformatics Institute, Hinxton, United Kingdom v 75. Genomics Institute, UC Santa Cruz, Santa Cruz, CA, USA

76. Carl Icahn Laboratory, Princeton University, Princeton, NJ, USA

77. Department of Population Health Sciences, The University of Utah, Salt Lake City, Utah, USA

78. Schools of Medicine, Engineering, and Public Health, Johns Hopkins University, Baltimore, MD, USA

79. Department of Biostatistics, Bloomberg School of Public Health, Johns Hopkins University, Baltimore, MD, USA

80. Department of Medical Biology, The Walter and Eliza Hall Institute of Medical Research, Parkville, Victoria, Australia

81. Altius Institute for Biomedical Sciences, Seattle, WA, USA v

82. Division of Genetics, University of Washington, Seattle, WA, University of Washington, Seattle, WA, USA

83. Department of Cardiology, University of Washington, Seattle, WA, USA

84. HudsonAlpha Institute for Biotechnology, Huntsville, AL, USA

85. Genome Sciences, University of Washington, Seattle, WA, USA

86. National Institute of Dental and Craniofacial Research, Bethesda, MD, USA

87. Division of Neuroscience and Basic Behavioral Science, National Institute of Mental Health, National Institutes of Health, Bethesda, MD, USA

88. National Institute on Drug Abuse, Bethesda, MD, USA

89. Office of Strategic Coordination, Division of Program Coordination, Planning and Strategic Initiatives, Office of the Director, National Institutes of Health, Rockville, MD, USA

90. Division of Genomic Medicine, National Human Genome Research Institute, Bethesda, MD, USA

## Acknowledgements

We thank the donors and their families for their generous gifts of organ donation for transplantation, and tissue donations for the GTEx research project; Mariya Khan and Christopher Stolte for the illustration in Fig. 1; and Laura Vairus for the illustrations in Figs. 2 and 3.

The consortium was funded by GTEx program grants: HHSN268201000029C (F.A., K.G.A., A.V.S., X.Li., E.T., S.G., A.G., S.A., K.H.H., D.Y.N., K.H., S.R.M., J.L.N.), 5U41HG009494 (F.A., K.G.A.), 10XS170 (Subcontract to Leidos Biomedical) (W.F.L., J.A.T., G.K., A.M., S.S., R.H., G.Wa., M.J., M.Wa., L.E.B., C.J., J.W., B.R., M.Hu., K.M., L.A.S., H.M.G., M.Mo., L.K.B.), 10XS171 (Subcontract to Leidos Biomedical) (B.A.F., M.T.M., E.K., B.M.G., K.D.R., J.B.), 10ST1035 (Subcontract to Leidos Biomedical) (S.D.J., D.C.R., D.R.V.), R01DA006227-17 (D.C.M., D.A.D.), Supplement to University of Miami grant DA006227. (D.C.M., D.A.D.), HHSN261200800001E (A.M.S., D.E.T., N.V.R., J.A.M., L.S., M.E.B., L.Q., T.K., D.B., K.R., A.U.), R01MH101814 (M.M-A., V.W., S.B.M., R.G., E.T.D., D.G-M., A.V.), U01HG007593 (S.B.M.), R01MH101822 (C.D.B.), U01HG007598 (M.O., B.E.S.), R01MH107666 (H.K.I.), P30DK020595 (H.K.I.). E.R.G. is supported by the National Human Genome Research Institute (NHGRI) under Award Number 1R35HG010718 and by the National Heart, Lung, and Blood Institute (NHLBI) under Award Number 1R01HL133559. E.R.G. has also significantly benefitted from a Fellowship at Clare Hall, University of Cambridge (UK) and is grateful to the President and Fellows of the college for a stimulating intellectual home. S.K.-H. is supported by the Marie-Sklodowska Curie fellowship H2020 Grant 706636. R.D.: R35GM124836, R01HL139865, 15CVGPSD27130014. D.M.J.: T32HL00782. Y.Pa. is supported by the NHGRI award R01HG10067. A.R.H. was supported by the Massachusetts Lions Eye Research Fund Grant. Computation was performed at the high performance cluster of the Center for Research Informatics at the University of Chicago, funded by the Biological Sciences Division and CTSA UL1TR000430. Additional Computation was performed with resources provided by the University of Chicago Research Computing Center.

We thank the International Genomics of Alzheimer’s Project (IGAP) for providing summary results data for these analyses. The investigators within IGAP contributed to the design and implementation of IGAP and/or provided data but did not participate in analysis or writing of this report. IGAP was made possible by the generous participation of the control subjects, the patients, and their families. The i–Select chips was funded by the French National Foundation on Alzheimer’s disease and related disorders. EADI was supported by the LABEX (laboratory of excellence program investment for the future) DISTALZ grant, Inserm, Institut Pasteur de Lille, Université de Lille 2 and the Lille University Hospital. GERAD was supported by the Medical Research Council (Grant n° 503480), Alzheimer’s Research UK (Grant n° 503176), the Wellcome Trust (Grant n° 082604/2/07/Z) and German Federal Ministry of Education and Research (BMBF): Competence Network Dementia (CND) grant n° 01GI0102, 01GI0711, 01GI0420. CHARGE was partly supported by the NIH/NIA grant R01 AG033193 and the NIA AG081220 and AGES contract N01–AG–12100, the NHLBI grant R01 HL105756, the Icelandic Heart Association, and the Erasmus Medical Center and Erasmus University. ADGC was supported by the NIH/NIA grants: U01 AG032984, U24 AG021886, U01 AG016976, and the Alzheimer’s Association grant ADGC–10–196728.

## Code and data availability

Genotype-Tissue Expression (GTEx) project’s raw whole transcriptome and genome sequencing data are available via dbGaP accession number phs000424.v8.p1. All processed GTEx data are available via GTEx portal. Imputed summary results, *enloc, coloc*, PrediXcan, MultiXcan, *dap-g*, prediction models, and reproducible analysis are available in https://github.com/hakyimlab/gtex-gwas-analysis and links therein.

## URLs

1000 Genomes Project Reference for LDSC,

https://data.broadinstitute.org/alkesgroup/LDSCORE/1000G_Phase3_plinkfiles.tgz;

1000 Genomes Project Reference with regression weights for LDSC,

https://data.broadinstitute.org/alkesgroup/LDSCORE/1000G_Phase3_weights_hm3_no_MHC.tgz;

BioVU, https://victr.vanderbilt.edu/pub/biovu/?sid=194;

eCAVIAR, https://github.com/fhormoz/caviar;

QTLEnrich, https://github.com/segrelabgenomics/eQTLEnrich;

flashr, https://gaow.github.io/mnm-gtex-v8/analysis/mashr_flashr_workflow.html#flashr-prior-covariances;

Gencode, https://www.gencodegenes.org/releases/26.html;

GTEx GWAS subgroup repository, https://github.com/broadinstitute/gtex-v8;

GTEx portal, http://gtexportal.org;

Hail, https://github.com/hail-is/hail;

HapMap Reference for LDSC, https://data.broadinstitute.org/alkesgroup/LDSCORE/w_hm3.snplist.bz2;

LD score regression (LDSD regression), https://github.com/bulik/ldsc;

MetaXcan, https://github.com/hakyimlab/MetaXcan;

Mouse Phenotype Ontology, http://www.informatics.jax.org/vocab/mp_ontology;

NHGRI-EBI GWAS catalog, https://www.ebi.ac.uk/gwas/;

picard, http://picard.sourceforge.net/;

PLINK 1.90, https://www.cog-genomics.org/plink2;

PrediXcan, https://github.com/hakyim/PrediXcan;

pyliftover, https://pypi.org/project/pyliftover/;

Storeys’ qvalue R package, https://github.com/StoreyLab/qvalue;

Summary GWAS imputation, https://github.com/hakyimlab/summary-gwas-imputation;

TORUS, https://github.com/xqwen/torus;

UK Biobank GWAS, http://www.nealelab.is/uk-biobank/;

UK Biobank, http://www.ukbiobank.ac.uk/;

## 1 Supplementary Materials

### 1.1 Terminology

For clarity and to reduce ambiguities, we provide the definition of some of the key terms used in the manuscript.

#### Trait

Here, trait (or complex trait) is used for observable, quantitative trait of individuals, such as presence of a disease or an anthropometric measurement. When speaking about traits, we do not include molecular phenotypes like gene expression or intron splicing quantification.

#### LD block/LD region

Region of the genome containing variants in LD among themselves, as determined from empirical LD patterns observed in 1000 Genomes [Berisa and Pickrell, 2016]. Variants in different LD blocks are unlikely to be correlated.

#### GWAS locus

This term is, in general, used somewhat loosely to refer to a region with a significantly associated variant which may span from tens to hundreds of kilobases depending on the LD of the region. However, here for quantification, we define it as one of the approximately independent LD blocks from [Berisa and Pickrell, 2016] that harbor a GWAS significant association. If multiple traits exist for a GWAS significant association in the block, we count them as distinct.

#### eQTL, eVariant

Here an eVariant is a genetic variant that is associated (FDR*<* 0.05) with the expression of a gene. eQTL refers to the variant-gene pair, in which the variant is an eVariant for the gene.

#### sQTL, sVariant

An sVariant is a genetic variant that is associated (FDR*<* 0.05) with the splicing (quantified as intron excision ratio) of a gene. sQTL refers to the variant-gene pair, in which the variant is an sVariant for the gene.

#### Fine-mapped variant

We call fine-mapped variant to the proxy for causal variant which we selected using *dap-g*’s posterior inclusion probabilities. These variants that are within credible sets with total posterior inclusion probability of at least 0.25 and have variant-level pip*>* 0.01. Within each credible set, one such variant is selected for our analysis.

#### LD contamination

This phenomenon occurs when the variant that alters the expression or splicing is distinct from the one that alters the complex trait, but they are in LD. In these circumstances, the QTL will be associated with the GWAS trait and the GWAS variant will be associated with the molecular trait, but there is no causal relationship between the gene and the complex trait.

#### Posterior inclusion probability (pip)

This is the probability that a variant has a causal effect on a trait. These probabilities are calculated by Bayesian fine-mapping approaches such as *dapg* and *susier*.

#### PrediXcan

This term refers to the family of methods that seeks to identify causal genes by correlating the genetic component of gene expression (mRNA level and splicing) with the trait. This family includes S-PrediXcan (which uses GWAS summary statistics rather than individual level data) and MultiXcan (which aggregates evidence of associations across all tissues leveraging the fact that e/sQTLs are shared across tissues). We use PrediXcan as a generic term to refer to this family of methods.

#### Silver standard genes

To test the ability of colocalization and association methods to identify true causal genes, we curated a set of ‘causal’ genes. To emphasize the imperfect nature, we use the term silver standard genes. In this context, the term OMIM gene is used as the causal gene for the trait.

### 1.2 Genotype-Tissue Expression (GTEx) Project

All processed Genotype-Tissue Expression (GTEx) Project v8 data have been made available on dbGAP (accession ID: phs000424.v8). Primary and extended results generated by consortium members are available on the Google Cloud Platform storage accessible via the GTEx Portal (see URLs).The GTEx Project v8 data, based on 17,382 RNA-sequencing samples from 54 tissues of 948 post-mortem subjects, has established the most comprehensive map of regulatory variation to date. In addition to the larger sample size and greater tissue coverage compared to v6, v8 data also included whole-genome sequencing data, facilitating high resolution QTL map of 838 subjects for 49 tissues with at least 70 samples. The GTEx consortium mapped complex trait associations for 23,268 cis-eGenes and 14,424 cis-sGenes [Aguet et al., 2019]. We did not include trans QTLs in our analyses due to limited power after correcting for confounders and potential pleiotropic effect in complex trait associations. Below, we briefly describe the whole-genome sequencing, RNA-sequencing and QTL data processing protocols. Detailed description of subject ascertainment, sample procurement, and sequencing data processing are available elsewhere [Aguet et al., 2019].

#### 1.2.1 Whole-genome sequence data processing and quality control

Out of 899 WGS samples sequenced at an average coverage of 30x on HiSeq200 (68 samples) and HiSeqX (all other samples), variant call files (VCF) for 866 GTEx donors were included in downstream analyses after excluding one each from 30 duplicate samples and three donors. Of these, 838 subjects with RNA-seq data were included for QTL mapping and subsequent complex trait association analyses in our study. All whole-genome sequencing data were mapped to GRCh38/hg38 reference.

#### 1.2.2 RNA-Seq data processing and quality control

Whole transcriptome RNA-Seq data were aligned using STAR (v2.5.3.a; [Dobin et al., 2013]). For STAR index, GENCODE v26 (GRCh38; see URLs) was used with the sjdbOverhang 75 for 76-bp paired-end sequencing protocol. Default parameters were used for RSEM (see URLs; [Li and Dewey, 2011]) index generation. GTEx utilized Picard (see URLs) to mark and remove potential PCR duplicates and RNA-SeQC [DeLuca et al., 2012] to process post-alignment quality control. RSEM was then used for per-sample transcript quantification. Subsequently, read counts were normalized between samples using TMM [Robinson and Oshlack, 2010]. For eQTL analyses, latent factor covariates were calculated using PEER as follows: 15 factors for N<150 per tissue; 30 factors for 150<=N<250; 45 factors for 250<=N<350; and 60 factors for N>=350. Finally, fastQTL [Ongen et al., 2016] was used for cis-eQTL mapping in each tissue. Only protein-coding, lincRNA, and antisense biotypes as defined by Gencode v26 were considered for further analyses. To study alternative splicing, GTEx applied LeafCutter (version 0.2.8; [Li et al., 2018]) using default parameters to quantify splicing QTLs in cis with intron excision ratios [Aguet et al., 2019].

### 1.3 Genome-wide association studies (GWAS) data

#### 1.3.1 Harmonization of GWAS summary statistics

The process followed for the harmonization and imputation are depicted in fig. S2. For each standardized GWAS summary statistics, we mapped all variants to hg38 (GRCh38) references using *pyliftover* (see URLs). For missing chromosome or genomic position information in the original GWAS summary statistics file, we queried dbSNP build 125 (hg17), dbSNP build 130 (hg18/GRCh36), and dbSNP build 150 (hg19/GRCh37) using the provided variant rsID information and the original reference build of the GWAS summary statistics file. Variants with missing chromosome, genomic position, and rsID information were excluded from further analyses. Only autosomal variants were included in our analyses. Missing allele frequency information was filled using the allele frequencies estimated in the GTEx (v8) individuals of genotype-based European genetic ancestry (here onwards, GTEx-EUR) whenever possible. We excluded variants with discordant reference and alternate allele information between GTEx and the GWAS study. We included only the alleles with the highest MAF among multiple alternate alleles if the variant was reported as multiallelic in GTEx. When more than one GWAS variant mapped to a given GTEx variant (i.e., the same chromosomal location in hg38), only the one with the highest significance was retained. For binary traits, if the sample size was present but the number of cases was missing, we filled the missing count with the sample size and number of cases reported in the paper. For continuous traits, if the file contained the sample size for each variant, the reported number was used. If not, we filled this value using the number reported in the corresponding publication. If only some variants were missing sample size information, we filled the missing value with the median of all reported values.

#### 1.3.2 Imputation of GWAS summary statistics

To standardize the number of variants across tissue-trait pairs, all processed GWAS results were imputed. We implemented the Best Linear Unbiased Prediction (BLUP) approach [Lee et al., 2013; Pasaniuc et al., 2014] in-house (https://github.com/hakyimlab/summary-gwas-imputation) to impute z-scores for those variants reported in GTEx without matching data in the GWAS summary statistics. This algorithm does not impute raw effect sizes (*β* coefficients). The imputation was performed in specific regions assumed to have sufficiently low correlations between them, defined by approximately independent linkage disequilibrium (LD) blocks [Berisa and Pickrell, 2016] lifted over to hg38/GRCh38.

Only GTEx variants with MAF *>* 0.01 in GTEx-EUR subjects were used in downstream analyses. Co-variance matrices (reference LD information) were estimated on these GTEx-EUR subjects. The corresponding (pseudo-)inverse matrices for covariances *C* were calculated via Singular Value Decomposition (SVD) using ridge-like regularization *C* + 0.1*I*. To avoid ambiguous strand issues homogeneously, palindromic variants (i.e. CG) were excluded from the imputation input. Thus, an imputed z-score was generated for palindromic variants available in the original GWAS; for them, we report the absolute value of the original entry with the sign from the imputed z-score. The sample size that we report for the imputed variants is the same as the sample size for the observed ones if it is reported as constant across variants, or their median if it changes across the observed variants, which occurs in the case of meta-analyses.

We initially considered publicly available GWAS summary statistics for 114 complex traits provided by large-scale consortia and the UK Biobank [Bycroft et al., 2018] (table S9). Of these, 27 studies with a relatively small intersection of variants with the GTEx panel (number of variants*<* 2 *×* 10^6^, compared to almost 9 *×* 10^6^ variants available in GTEx) exhibited significant deflation of their association p-values (fig. S4). Thus, all analyses focused on 87 traits where missing variants could be properly imputed unless otherwise stated explicitly (table S2). We observed noteworthy association prediction performance across the selected 87 traits (e.g., with a median *r*^2^ = 0.90 (IQR = 0.0268) between the original and imputed zscores on chromosome 1). The median slope was 0.94 (IQR = 0.0164), as the imputed zscore values tend to be more conservative than the original ones. Imputation quality was consistent across traits, depending strongly on the number of input available variants (fig. S3).

**Supplementary Fig. S1.**
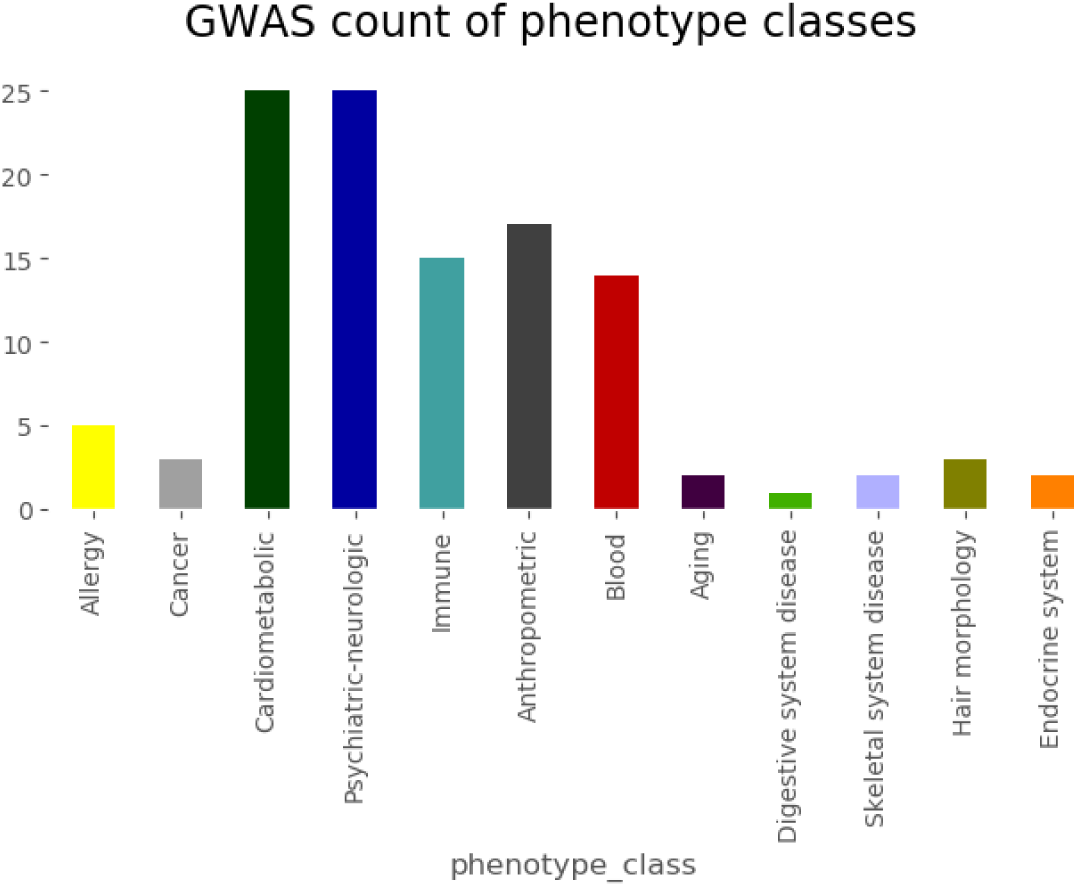
GWAS trait categories. Categories of the traits with full GWAS summary statistics used in the analysis. See list of traits in S2.

**Supplementary Fig. S2.**
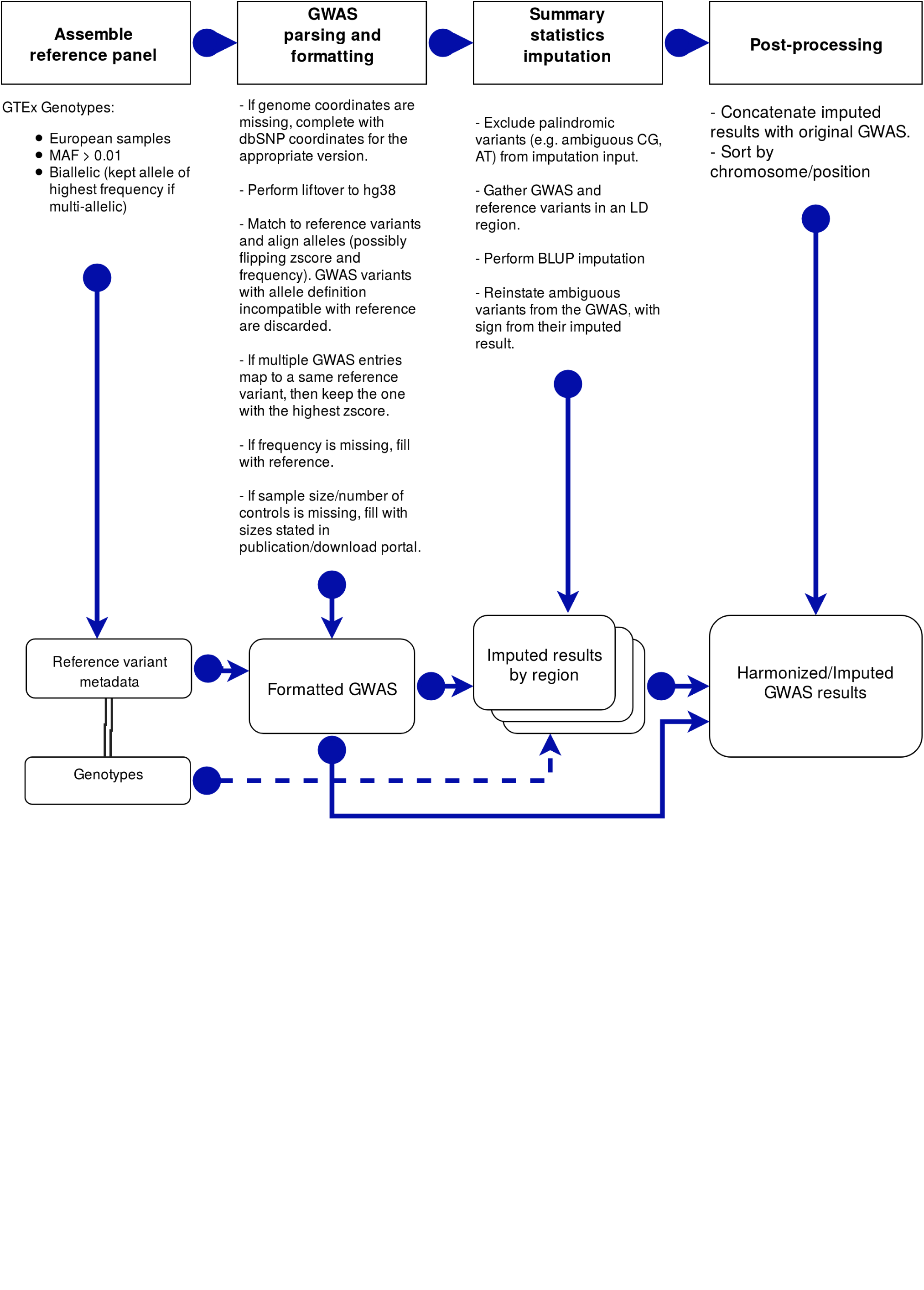
Workflow of GWAS results processing.

**Supplementary Fig. S3.**
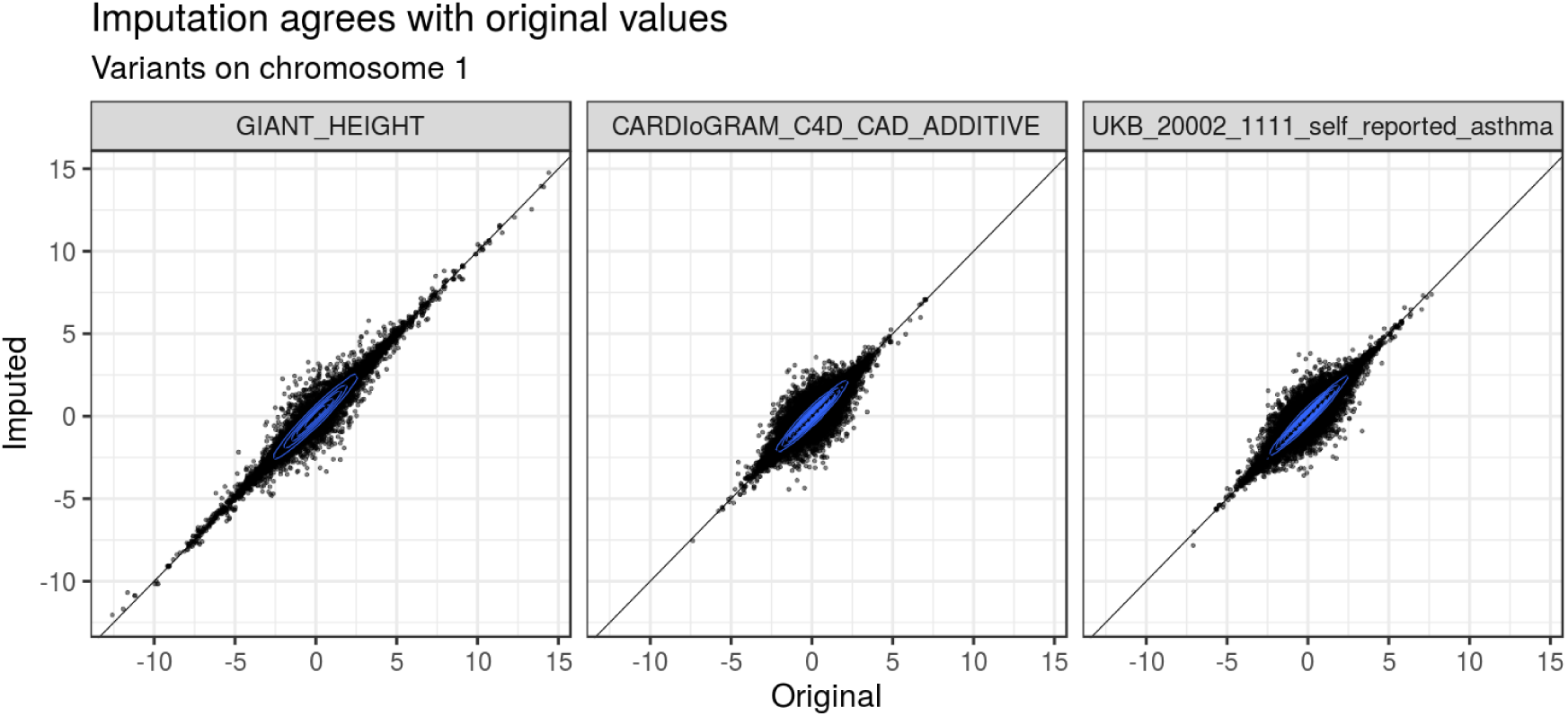
GWAS imputation quality. Original versus imputed zscores for palindromic variants in chromosome 1 for 3 traits.

**Supplementary Fig. S4.**
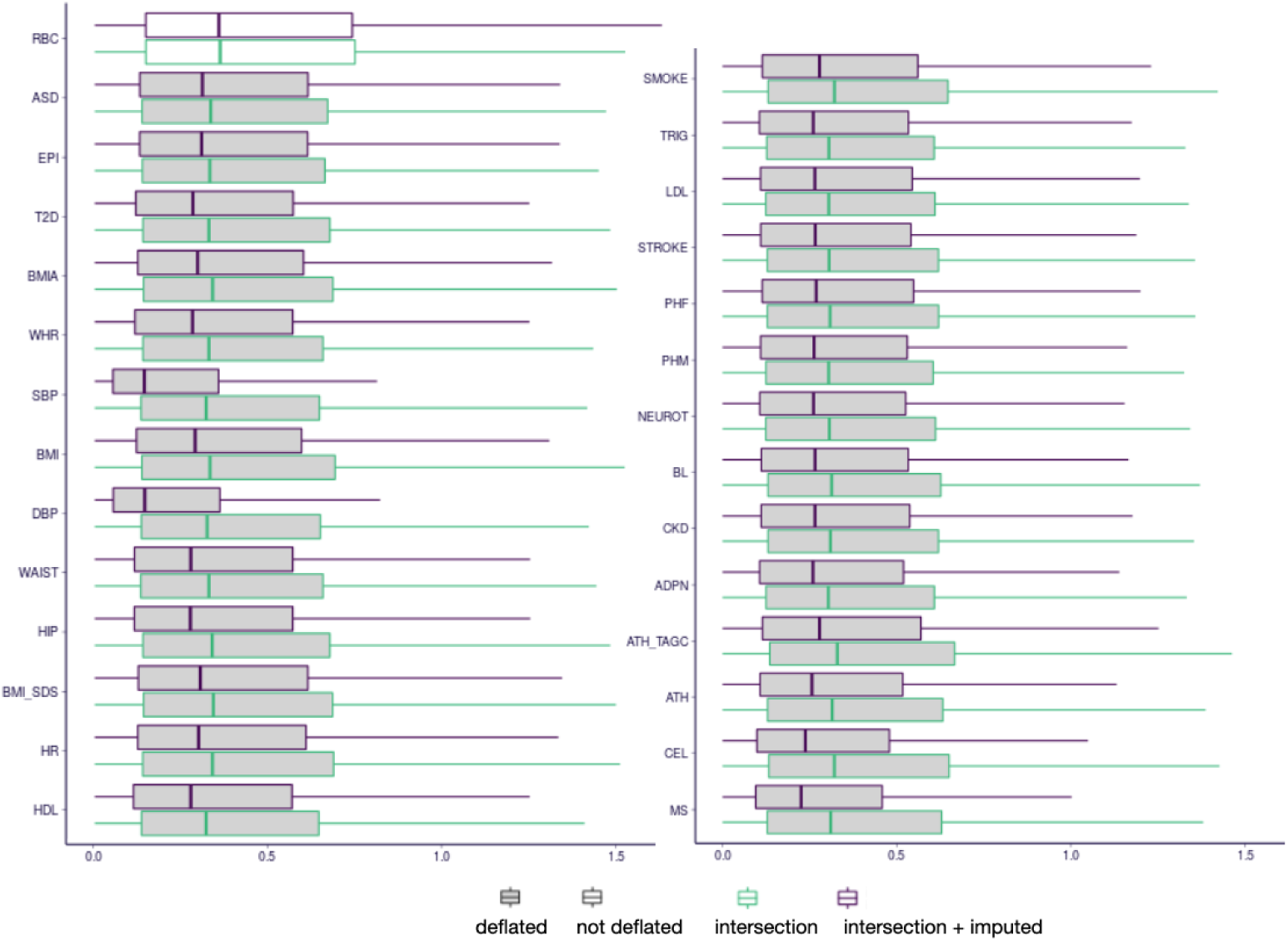
GWAS imputation deflation. This figure compares the distribution of p-values for 28 GWAS traits before and after imputation. Vertical scale shows -log10(p-value) of variant association. The 27 traits that exhibited deflation are filled in gray. An undeflated trait (e.g., Red Blood Cell count) is included for comparison. See trait abbreviation list in table S9.

**Table S1:** List of 87 GWAS datasets

| Category | Trait | Abbreviation | Sample_Size |
| --- | --- | --- | --- |
| Psychiatric-neurologic | Alzheimers Disease | AD | 54162 |
| Psychiatric-neurologic | Attention Deficit Hyperactivity Disorder | ADHD | 53293 |
| Psychiatric-neurologic | Chronotype | CHRONO | 128266 |
| Psychiatric-neurologic | Chronotype UKB | CHRONO_UKB | 337119 |
| Psychiatric-neurologic | Depressive Symptoms | DEPR | 180866 |
| Psychiatric-neurologic | Education Years | EDU | 293723 |
| Psychiatric-neurologic | Epilepsy UKB | EPI_UKB | 337119 |
| Psychiatric-neurologic | Fluid Intelligence Score UKB | FIS_UKB | 337119 |
| Psychiatric-neurologic | Insomnia In Both Sexes | INSOMN | 113006 |
| Psychiatric-neurologic | Insomnia UKB | INSOMN_UKB | 337119 |
| Psychiatric-neurologic | Insomnia UKBS | INSOMN_UKBS | 337119 |
| Psychiatric-neurologic | Migraine UKB | MIGR_UKB | 337119 |
| Psychiatric-neurologic | Migraine UKBS | MIGR_UKBS | 337119 |
| Psychiatric-neurologic | Multiple Sclerosis UKBS | MS_UKBS | 337119 |
| Psychiatric-neurologic | Neuroticism UKB | NEUROT_UKB | 337119 |
| Psychiatric-neurologic | Parkinsons Disease UKBS | PD_UKBS | 337119 |
| Psychiatric-neurologic | Psychological Problem UKBS | PSY_UKBS | 337119 |
| Psychiatric-neurologic | Schizophrenia | SCZ | 150064 |
| Psychiatric-neurologic | Schizophrenia UKBS | SCZ_UKBS | 337119 |
| Psychiatric-neurologic | Sleep Duration | SLEEP | 128266 |
| Psychiatric-neurologic | Sleep Duration UKB | SLEEP_UKB | 337119 |
| Anthropometric | BMI UKB | BMI_UKB | 337119 |
| Anthropometric | Birth Weight | BW | 143677 |
| Anthropometric | Birth Weight UKB | BW_UKB | 337119 |
| Anthropometric | Body Fat Percentage UKB | FAT_UKB | 337119 |
| Anthropometric | Bone Mineral Density | BMD | 49988 |
| Anthropometric | Height | HEIGHT | 253288 |
| Anthropometric | Intracranial Volume | ICV | 30717 |
| Anthropometric | Standing Height UKB | HEIGHT_UKB | 337119 |
| Cardiometabolic | CH2DB NMR | CH2 | 24154 |
| Cardiometabolic | Coronary Artery Disease | CAD | 184305 |
| Cardiometabolic | Deep Venous Thrombosis UKB | DVT_UKB | 337119 |
| Cardiometabolic | Deep Venous Thrombosis UKBS | DVT_UKBS | 337119 |
| Cardiometabolic | Fasting Glucose | FG | 46186 |
| Cardiometabolic | Fasting Insulin | INSUL | 38238 |
| Cardiometabolic | HDL Cholesterol NMR | HDLC | 19270 |
| Cardiometabolic | Heart Attack UKB | MI_UKB | 337119 |
| Cardiometabolic | High Cholesterol UKBS | HC_UKBS | 337119 |
| Cardiometabolic | Hypertension UKBS | HPT_UKBS | 337119 |
| Cardiometabolic | LDL Cholesterol NMR | LDLC | 13527 |
| Cardiometabolic | Pulmonary Embolism UKB | PE_UKB | 337119 |
| Cardiometabolic | Triglycerides NMR | IDL | 21559 |
| Cardiometabolic | Type 2 Diabetes UKBS | T2D_UKBS | 337119 |
| Blood | Eosinophil Count | EC | 173480 |
| Blood | Granulocyte Count | GC | 173480 |
| Blood | High Light Scatter Reticulocyte Count | HRET | 173480 |
| Blood | Lymphocyte Count | LC | 173480 |
| Blood | Monocyte Count | MC | 173480 |
| Blood | Myeloid White Cell Count | MWBC | 173480 |
| Blood | Neutrophil Count | NC | 173480 |
| Blood | Platelet Count | PLT | 173480 |
| Blood | Red Blood Cell Count | RBC | 173480 |
| Blood | Reticulocyte Count | RET | 173480 |
| Blood | Sum Basophil Neutrophil Count | BNC | 173480 |
| Blood | Sum Eosinophil Basophil Count | EBC | 173480 |
| Blood | Sum Neutrophil Eosinophil Count | NEC | 173480 |
| Blood | White Blood Cell Count | WBC | 173480 |
| Cancer | Breast Cancer | BC | 120000 |
| Cancer | ER-negative Breast Cancer | ERNBC | 120000 |
| Cancer | ER-positive Breast Cancer | ERPBC | 120000 |
| Allergy | Asthma UKBS | ATH_UKBS | 337119 |
| Allergy | Eczema | ECZ | 116863 |
| Allergy | Eczema UKBS | ECZ_UKBS | 337119 |
| Immune | Ankylosing Spondylitis UKBS | ASP_UKBS | 337119 |
| Immune | Asthma UKB | ATH_UKB | 337119 |
| Immune | Crohns Disease | CD | 20833 |
| Immune | Crohns Disease UKBS | CD_UKBS | 337119 |
| Immune | Hayfever UKB | HAY_UKB | 337119 |
| Immune | Inflammatory Bowel Disease | IBD | 34652 |
| Immune | Inflammatory Bowel Disease UKBS | IBD_UKBS | 337119 |
| Immune | Psoriasis UKBS | PSO_UKBS | 337119 |
| Immune | Rheumatoid Arthritis | RA | 80799 |
| Immune | Rheumatoid Arthritis UKBS | RA_UKBS | 337119 |
| Immune | Systemic Lupus Erythematosus | SLE | 23210 |
| Immune | Type 1 Diabetes UKBS | T1D_UKBS | 337119 |
| Immune | Ulcerative Colitis | UC | 27432 |
| Immune | Ulcerative Colitis UKBS | UC_UKBS | 337119 |
| Aging | Fathers Age At Death UKB | FAD_UKB | 337119 |
| Aging | Mothers Age At Death UKB | MAD_UKB | 337119 |
| Digestive system disease | Irritable Bowel Syndrome UKBS | IBS_UKBS | 337119 |
| Endocrine system disease | Hyperthyroidism UKBS | HYPERTHY_UKBS | 337119 |
| Endocrine system disease | Hypothyroidism UKBS | HYPOTHY_UKBS | 337119 |
| Skeletal system disease | Gout UKBS | GOUT_UKBS | 337119 |
| Skeletal system disease | Osteoporosis UKBS | OST_UKBS | 337119 |
| Morphology | Balding Pattern 2 UKB | BLDP2_UKB | 337119 |
| Morphology | Balding Pattern 3 UKB | BLDP3_UKB | 337119 |
| Morphology | Balding Pattern 4 UKB | BLDP4_UKB | 337119 |

#### 1.3.3 IGAP GWAS

We used summary results from an Alzheimer’s Disease study from International Genomics of Alzheimer’s Project (IGAP).

IGAP is a large two-stage study based upon genome-wide association studies (GWAS) on individuals of European ancestry. In stage 1, IGAP used genotyped and imputed data for 7,055,881 single nucleotide polymorphisms (SNPs) to meta-analyze four previously-published GWAS datasets consisting of 17,008 Alzheimer’s disease cases and 37,154 controls (The European Alzheimer’s disease Initiative - EADI the Alzheimer Disease Genetics Consortium - ADGC The Cohorts for Heart and Aging Research in Genomic Epidemiology consortium - CHARGE The Genetic and Environmental Risk in AD consortium - GERAD). In stage 2, 11,632 SNPs were genotyped and tested for association in an independent set of 8,572 Alzheimer’s disease cases and 11,312 controls. Finally, a meta-analysis was performed combining results from stages 1 & 2.

#### 1.3.4 NHGRI-EBI GWAS catalog

In addition to the GWAS summary statistics described above, we obtained the list of trait-associated SNPs from the GWAS catalog [Buniello et al., 2019] (downloaded on 9/7/2018), which, at download, contained 80,727 entries. To measure the enrichment of e/sQTL in the GWAS Catalog, we computed the proportion of e/sQTL in the GWAS catalog relative to the proportion of e/sQTL among all GTEx V8 variants. We then obtained a measure of the uncertainty in the proportion and enrichment-fold using block jackknife. See [Aguet et al., 2019] for details.

### 1.4 Correlated t-test to summarize across traits and tissues

Most statistics shown in these analyses are at the tissue-trait level. There are 4,263 statistics, generated from 49 tissues and 87 traits. Typical statistical tests assume the data from which the statistic is computed is sampled independently and identically distributed (IID). Among different tissues, there are wide ranges of standard errors and different patterns of correlation. Because of this, the IID assumption can not be applied to the tissue-trait statistics. Therefore, we describe our derivation of standard errors when statistics are summarized across traits for a given tissue, and when statistics are summarized across tissue and trait pairs. In the following paragraphs, we use *S*_*tp*_ to indicate a statistic estimated in tissue *t* and trait *p*. This statistic has standard error se(*S*_*tp*_).

#### Summarizing across traits for a given tissue

When we have one statistic per tissue-trait pair and summarize across traits in a given tissue, we assume the traits are independent, but we take into account the differences in standard errors. For each tissue *t*, we summarized *S*_*t*1_, …, *S*_*tP*_ by fitting the following linear model:

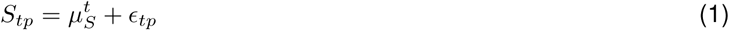

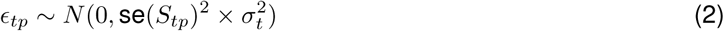

So 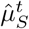 is an estimate for the statistic *S* summarized across all traits in tissue *t*, and this estimate has standard error 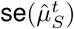). This is essentially a weighted average across traits.

#### Summarizing across trait and tissue pairs

When we summarize across all tissue-trait pairs, *S*_11_, &, *S*_*tp*_, &, *S*_*T P*_, we fit a similar linear model, which allows for correlation between tissues and correlation between traits, and corrects for differences in the standard errors.

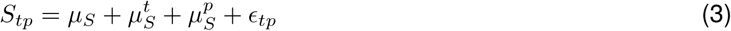

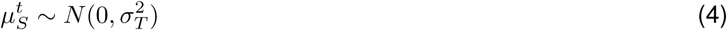

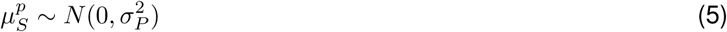

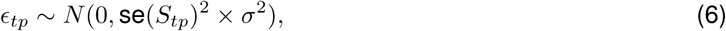

Here, 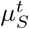 is the tissue-specific random intercept, and 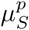 is the trait-specific random intercept. These components account for features common across traits that are specific to tissue *t* and features common across tissues that are specific to trait *p* respectively. The estimate 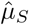 is the weighted average of *S*_*tp*_ across all tissue-trait pairs, and its standard error is 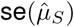.

#### Testing whether two statistics have different mean

We would often like to test whether two statistics are different, *e*.*g*. enrichment signal measured for sQTL as 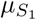 versus enrichment signal measured for eQTL as 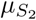. For this, we need to construct a test aggregating pairwise differences across all tissue-trait pairs. For this purpose, we constructed the following paired test. Our test statistic is *T* ^*tp*^ := *S*_1,*tp*_ − *S*_2,*tp*_ with se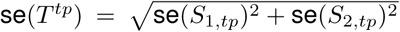. We calculate 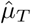 by summarizing across all tissue-trait pairs as described in the previous paragraph. Under the null 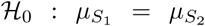 and 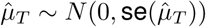.

### 1.5 Enrichment of QTLs among trait-associated variants

To estimate the proportion of SNPs considered as associated with expression (for at least one gene) at various p-value thresholds, we used the most significant p-value (tested using all GTEx individuals) for each SNP from all associations in all tissues (including all genes and variants tested). We observed that the proportion of variants associated with expression and splicing at different significance threshold was much larger for trait-associated variants from the GWAS catalog than for the full set of tested common variants (fig. S5). At a nominal threshold, the proportion of common variants associated with the expression of a gene in some tissue increased from 92.7% in the V6 release [GTEx Consortium et al., 2017] to 97.3% in V8. For splicing, the proportion was 97.7%. These results should serve as a cautionary note that assigning function to a GWAS locus based on QTL association p-value alone, even with a more stringent threshold, could be misleading.

**Supplementary Fig. S5.**
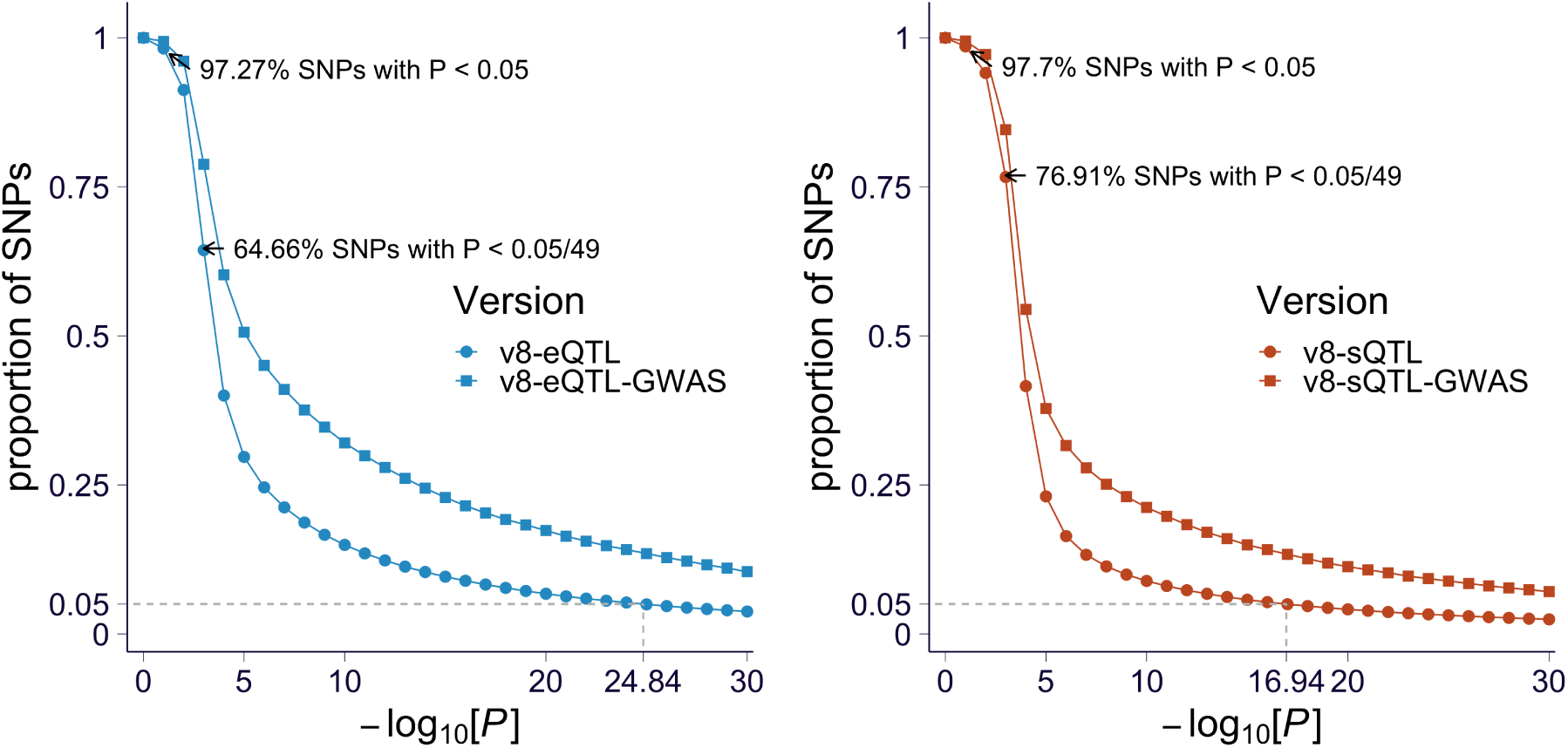
Expression and splicing QTL enrichment among GWAS variants. The proportion of genetic variants associated with gene expression **(A)** and splicing **(B)** of at least one gene in at least one tissue for each p-value cutoff (on x-axis in −log_10_(*p*) scale) is shown. The proportions for all tested variants are shown as squares and the proportions for the GWAS catalog variants are shown as circles.

### 1.6 Cis-region and covariates used in fine-mapping and prediction of expression and splicing traits

For each gene, we considered all variants within the cis-window (1Mbps) with MAF*>*0.01, and used the same covariates as in the GTEx v8 main eQTL analysis: sex, WGS plaform, WGS library preparation protocol, top 5 genetic principal components, and PEER factors. The number of PEER factors was determined from the sample size: 15 for *n <* 150, 30 for 150 ≤ *n <* 250, 45 for 250 ≤ *n <* 350, 60 for 350 ≤ *n*.

### 1.7 Fine-mapping expression and splicing QTLs

We applied *dap-g* [Wen, 2016] to the 49 tissues to estimate the degree to which a variant might exert a causal effect on expression or splicing levels, using default parameter values. First, we selected genes annotated as protein-coding, lincRNA or pseudogenes. We used the covariates listed in the supplement section 1.6. This yielded a list of clusters (variants related by LD), and posterior inclusion probabilities (*pip*) that provide an estimate of the probability of a variant being causal. We repeated this process for splicing ratios from Leafcutter, using a cis-window ranging from 1Mbps upstream of the splicing event start location to 1Mbps downstream of the end location. We used individual-level data for GTEx-EUR subjects both for expression and splicing. We note that the main report of the GTEx v8 included individuals of non-European descent and reported only expression QTL fine-mapping. Sample sizes ranged from 65 in kidney cortex to 602 in skeletal muscle tissues. All results are made publicly available (https://github.com/hakyimlab/gtex-gwas-analysis).

### 1.8 Mediation analysis to quantify the dose-dependent effects of expression and splicing on traits

Enrichment of expression and splicing QTLs suggest a causal role of molecular trait regulation on complex traits. However, confounders such as LD contamination could be inflating these results limiting their interpretation. Here, we sought to gather stronger evidence for a causal link. We tested whether there is a dose-dependent effect of expression and splicing QTLs on complex traits and also whether independent QTLs provided similar measures of the mediated effects.

#### 1.8.1 Selection of fine-mapped variants as instrumental variables and their effect sizes

To investigate the relationship between GWAS and QTL effect sizes in the transcriptome, we generated a set of fine-mapped QTL signals derived from *dap-g* fine-mapping performed in the GTEx-EUR individuals to serve as proxy for causal QTLs. For splicing, we utilized sQTLs at the splicing event/variant level rather than the gene/variant level. We considered only variants within credible sets with at least 25% total probability. Within each credible set, the variant with highest posterior inclusion probability was selected as the fine-mapped variant. Only variants with variant-level *pip* of at least 0.01 were considered.

For each of the selected QTLs, we used the QTL effect size estimated from the marginal test (using the GTEx-EUR individuals) and the GWAS effect size reported by the study or if missing, calculated from the imputed z-score from the GWAS imputation by 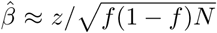, where *f* is the allele frequency and *N* is the GWAS sample size.

#### 1.8.2 Correlation between GWAS and QTL effect sizes

To get a first-order approximation to the mediated effect sizes without imposing any modeling assumptions, we calculated the Pearson correlation of the magnitude of observed GWAS effect size and of cis-eQTL effect size, 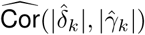, for the list of selected fine-mapped QTLs. This was done for each tissue-trait pair separately. The observed Pearson correlation captures the mediated effect (see details in Section 1.8.5). To obtain a null distribution for the correlation that accounts for the potential confounding effect of different local LD score values, we computed the Pearson correlation under the shuffled data within each LD-score bin defined by quantiles (100 bins were used). The significance of the difference between observed and null distribution was calculated using the correlated t-test method described in Section 1.4.

#### 1.8.3 Modeling effect mediated by regulatory process

We compared the magnitude of GWAS and cis-QTL effect sizes, which is the basis of multi-SNP Mendelian randomization approaches [Bowden et al., 2015].

To formalize the relationship between the GWAS effect size (*δ*) and the QTL effect size (*γ*), we assumed an additive genetic model for the GWAS trait. Specifically, for variant *k*,

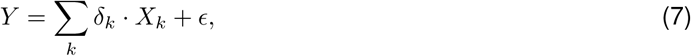

where *X*_*k*_ is the allele count of variant *k, Y* is the trait, and *E* is the un-explained variation. We decomposed GWAS effect size into its mediated and un-mediated components,

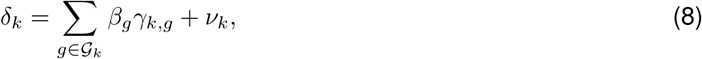

where 𝒢_*k*_ represents the set of genes regulated by variant *k* with corresponding QTL effect size as *γ*_*k,g*_, and *ν*_*k*_ is the un-mediated effect of variant *k* on trait. And *β*_*g*_ is the downstream effect of gene *g* on the trait.

##### 1.8.4 Transcriptome-wide estimation of mediated effects

To estimate the transcriptome-wide contribution of the mediated effects on complex traits, we proposed a mixed-effects model on the basis of Eq. 8,

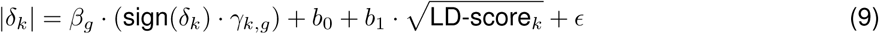

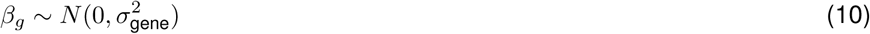

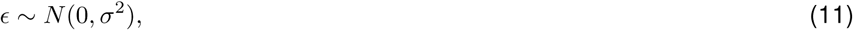

where *b*_0_, *b*_1_ are the fixed effect capturing the un-mediated effect and *β*_*g*_ is the mediated effect of the gene or splicing event *g*. In short, we assumed a random effects model to account for the heterogeneity of the *β*’s and aimed at estimating 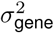 as the transcriptome-wide average of the mediated effect. For each tissue-trait pair, we fitted the model using selected fine-mapped QTLs, as described in Section 1.8.1, along with the corresponding 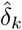 (GWAS effect for variant *k*), 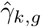 (QTL effect for variant *k*, gene *g*). To obtain the distribution of 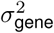 under the null, we performed the same calculation using shuffled GWAS effect sizes. The effect allele choice is arbitrary and we chose them so that all the GWAS effects are positive. This choice made the modeling of the effect of local LD more straightforward since we expect that variants in high LD regions may tag more causal variants and end up with a larger estimated GWAS effect, which would result in a positive *b*_1_. The square root of LD-score represents better the potential effect of LD score on the effect size. Using absolute value of the GWAS effects (| *δ*_*k*_|) and sign(*δ*_*k*_) *· γ*_*k,g*_ in equation (9) is a convenient way to implement the recoding of the effect allele.

#### 1.8.5 Robustness of the estimation of the mediating effect to LD contamination

We illustrate the intuition behind the LD-contamination correction when the average mediated effects are estimated using the approximate method (correlation of absolute values) or the mixed-effects approach.

**Supplementary Fig. S6.**
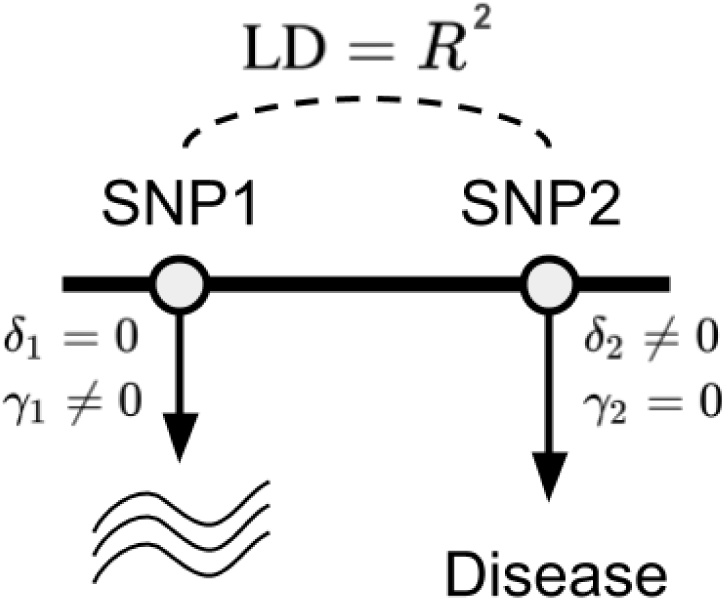
Schematic representation of LD contamination. SNP1 has a causal effect on the expression level of a gene but not on the trait (Disease here), *δ*_1_ = 0 and *γ*_1_ */*= 0. SNP2 has a causal effect on the trait but not on the expression of the gene, *δ*_2_ */*= 0 and *γ*_2_ = 0.

Consider the LD-contamination scenario where SNP 1 and SNP 2 are in LD with correlation *R*^2^ (suppose LD is fixed) and have a non-zero effect on gene expression and trait, respectively (as shown in fig. S6). The marginal effect estimates of SNP 1, *i*.*e*. 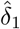 and 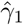, are given by

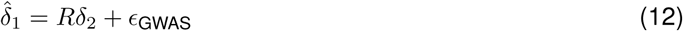

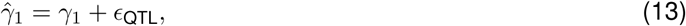

where Eq. 12 holds because the marginal effect size depends on LD. To determine the covariance of the magnitude of the GWAS and QTL estimates for SNP 1, we consider E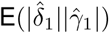.

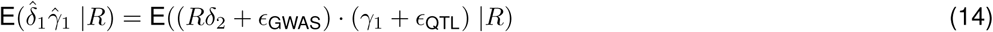

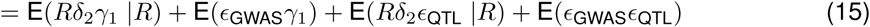

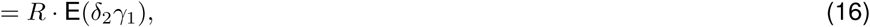

where Eq. 16 holds since the last three terms in the previous line are zeros, due to the independence among *ϵ*_GWAS_, *ϵ*_QTL_, and true effect sizes, *δ* and *γ*.

Hence, the covariance of the GWAS and QTL effect sizes under the LD contamination scenario is

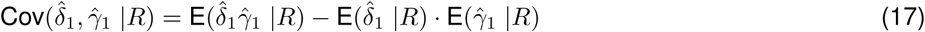

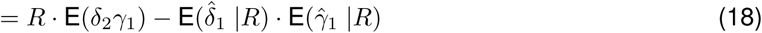

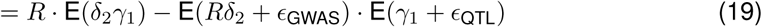

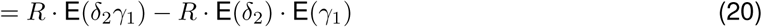

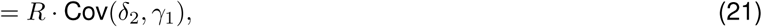

which implies that conditioning on LD, the observed correlation between 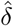 and 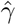 should be very small.

**Supplementary Fig. S7.**
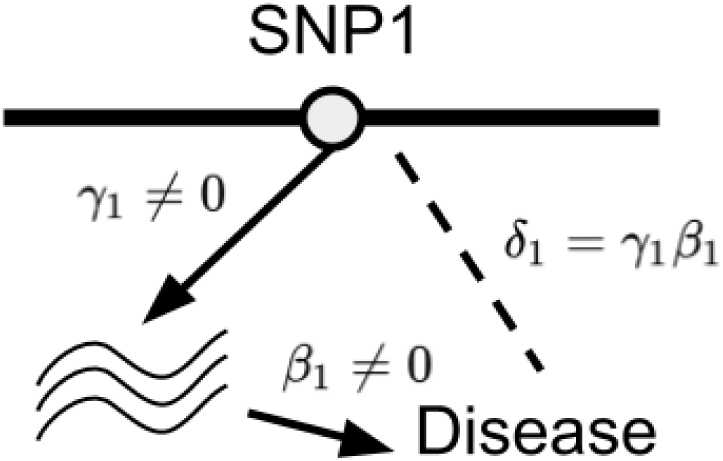
Diagram representation of mediation model.

Similarly, we can derive the correlation between GWAS and QTL effect size estimates under the simple mediation model shown in fig. S7, where we have

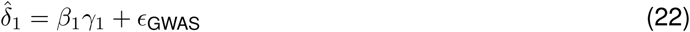

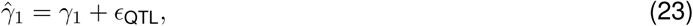

where Eq. 22 follows by definition of the mediation model considering no direct effect. So,

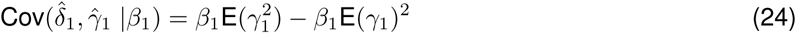

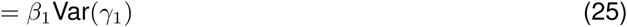

So, if we consider a gene locus, which naturally conditions on local LD and gene-level effect *β*, we can conclude that

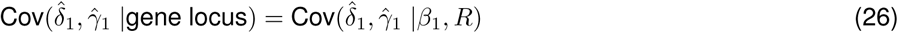

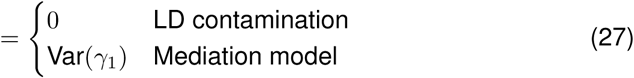

#### 1.8.6 Concordance of mediated effects for allelic series of independent eQTLs

Under the mediation model in Eq. 8, we expect that for a given gene with multiple QTL signals, these signals should share the same downstream effect, *β*_*g*_. Since the number of splicing events with multiple QTL signals was limited, we restricted this analysis to eQTLs only. We tested for concordance of downstream effect size obtained from the primary and secondary eQTL of a gene (ranked by QTL significance or QTL effect size estimate). Specifically, for a given trait and gene *g*, we defined the observed downstream effect for the *k*th variant as 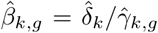. Thus, for each gene, we obtained 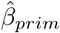 and 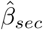 as the observed downstream effect for the primary and secondary eQTLs if more than one eQTL signal was detected by *dap-g*. Ideally, for a mediating gene in a causal tissue (or a good proxy tissue), we would expect that 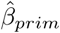 and 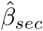 should be similar. We measured the concordance in two ways: 1) correlation between 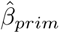 and 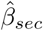; 2) percent concordant, defined as the fraction of eQTL pairs having the same sign in 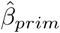 and 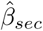. The results of 1) were reported in [Aguet et al., 2019].

##### Visualizing the concordance among colocalized genes

To visualize the concordance of 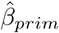 and 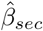, we first scaled 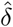 and 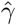 by their standard deviation among all eQTLs selected in Section 1.8.1. Then, we extracted the set of genes with exactly two *dap-g* eQTLs (as described in 1.8.1) and labelled the two eQTLs as primary and secondary based on QTL significance or QTL effect size. We computed 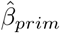 and 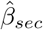 and removed the genes with 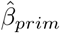 or 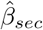 in the top and bottom 5%. As a control, we also simulated random *δ* to compute simulated *β*_*sim*_ for downstream analysis. We further filtered the genes by selecting only those with *enloc* rcp > 0.1.

### 1.9 Identifying patterns of regulation of expression across tissues

We used FLASH Sparse Factor Analysis [Wang and Stephens, 2018] to identify latent factors specific to different tissue clusters. We ran flashr on a set of top eQTLs (obtained from all GTEx individuals) per gene which had been tested in all 49 tissues (around 16,000 eQTLs in total were selected) and shown strong evidence of being active in at least one tissue. Then, for each selected variant-gene pair, the marginal effect size estimates were extracted for all 49 tissues regardless of whether it was significant in that tissue or not. The resulting estimated effect-size matrix (of dimension ∼16, 000 *×* 49) was the input to flashr (with normal prior on loading and uniform with positive support as prior on factor) to obtain the sparse factors. The flashr run yielded 31 FLASH factors (fig. S23), which were used to assign the tissue-specificity of an eQTL.

We defined the eQTL cross-tissue patterns by projecting the estimated effect-size vector across 49 tissues onto the FLASH factors and computed the quality of the projection, PVE, as 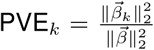. PVE_*k*_ represented the quality score for using FLASH factor *k* to explain the cross-tissue pattern of eQTL. The eQTL was assigned to a FLASH factor *k* if PVE_*k*_ was maximal among all FLASH factors and PVE_*k*_ *>* 0.2 and for those with PVE_*k*_ ≤ 0.2 in all FLASH factors, NA (short for not assigned) was assigned instead. These “not assigned” eQTLs had more complex tissue-sharing pattern than the factors captured in the FLASH analysis. To obtain an interpretable tissue-specificity category, we labeled *Factor1* as the shared factor, *Factor2, Factor13, Factor14, Factor29*, and *Factor30* as brain-specific factors, and the rest of the factor assignment as *other factors*.

We applied the multivariate adaptive shrinkage implemented in *mashr* [Urbut et al., 2018] to smooth cis-eQTL effect size estimates (obtained from all GTEx individuals) by taking advantage of correlation between tissues. To fit the *mashr* model, we used the set of ∼16, 000 cis-eQTLs as stated in Section 1.9 to learn the *mashr* prior, and then fit the *mashr* model using ∼40, 000 randomly selected variant-gene pairs for the same set of eGenes. We learned data-driven *mashr* priors in three ways: 1) FLASH factors as described above; 2) PCA with number of PC = 3; 3) empirical covariance of observed z-scores. The data-driven covariances were further denoised by calling cov_ed in mashr. Furthermore, we included the set of canonical covariances as described in [Urbut et al., 2018] as an additional *mashr* prior. We fit the *mashr* model using the set of randomly selected variant-gene pairs with the error correlation estimated by applying estimate_null_correlation function in mashr and the priors obtained above. The resulting *mashr* model was used to compute the posterior mean, standard deviation, and local false sign rate (LFSR) for any variant-trait pair.

### 1.10 Causal gene prioritization

Two classes of methods can be used to identify the target genes of GWAS loci. One class is based on the colocalization of GWAS and QTL loci, which seeks to determine whether the causal variant for the trait is the same as the causal variant for the molecular phenotype. The other class is based on the association between the genetically regulated component of gene expression (or splicing) with the trait. We applied representative examples of each class of methods.

#### 1.10.1 Colocalization

For a given variant associated with multiple traits such as gene expression (eQTL) and complex disease (trait-associated variant), extensive LD makes it challenging to identify the underlying true causal mechanisms. Colocalization approaches attempt to address this problem. Here, we conducted colocalization analysis using two independent approaches: *coloc* [Giambartolomei et al., 2014] and *enloc* [Wen et al., 2017]), to estimate whether a gene’s expression or a splicing event shares a causal variant with a trait.

#### 1.10.2 enloc

We computed Bayesian regional colocalization probability (rcp) using *enloc*, to estimate the probability of a GWAS region and a gene’s cis window sharing causal variants. We used the *dap-g* results described in 1.7, which was based on EUR individuals only. We split the GWAS summary statistics into approximately LD-independent regions [Berisa and Pickrell, 2016], each region defining a GWAS locus. For each tissue-trait combination, we computed the rcp of every overlapping GWAS locus to a gene’s or splicing event’s cis window with *enloc*’s default execution mode.

For each trait, we counted the number of GWAS loci that contain a GWAS significant hit, and among these, the number of loci that additionally contain a gene with *enloc* colocalization *rcp >* 0.5. As shown in fig. S12C, across traits, a median 29% of loci with a GWAS signal contain an *enloc* colocalized signal. Given *enloc*’s conservative nature, we caution that *rcp <* 0.5 does not mean that there is no causal relationship between the molecular phenotype and the complex trait; rather, it should be interpreted as lack of sufficient evidence with current data. We summarize the findings in fig. S13. We observed a smaller proportion of GWAS loci containing a colocalized splicing event (median 11% across traits).

#### 1.10.3 coloc

We computed *coloc* on all cis-windows with at least one eVariant (cis-eQTL per-tissue q-value*<* 0.05) or sVariant. For each gene’s cis-window, we used summary statistics from the GWAS traits and the main GTEx eQTL/sQTL analysis. For binary traits, case proportion and ‘cc’ trait type parameters were used. For continuous traits, sample size and ‘quant’ trait type parameters were used. In both cases, imputed or calculated z-scores were used as effect coefficients in Bayes factor calculations.

*Coloc* is very sensitive to the choice of priors. We used *enloc*’s enrichment estimates to define data-based priors in a consistent manner. First, we defined likely LD-independent blocks of variants using definitions provided previously [Berisa and Pickrell, 2016]. The probability of eQTL signal, Pr(*d*_*i*_ = 1), was estimated using *dap-g* [Wen, 2016]. Subsequently, we calculated priors *p*_1_, *p*_2_, and *p*_12_ for colocalization analyses as follows:

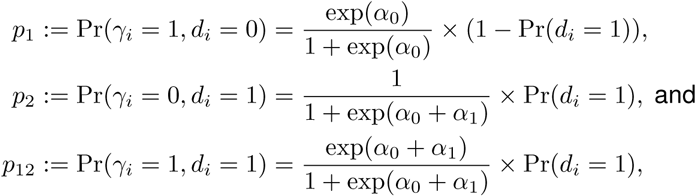

where *α*_0_ and *α*_1_ indicate intercept effect estimate and log odds ratio estimate for the enrichment using *enloc*, respectively.

We ran *coloc* using variants in the cis-window for each gene and the intersection with each GWAS trait, obtaining five probabilities for each gene-tissue-trait tuple: **P0** for the probability of neither expression nor GWAS having a causal variant; **P1** for the probability of only expression having a causal variant; **P2** for only the GWAS having a causal variant; **P3** for the GWAS and expression traits to have distinct causal variants; **P4** for the GWAS and expression traits to have a shared causal variant. We repeated this process using sQTL results.

### 1.11 Fine-mapping of height GWAS using summary statistics

To investigate the robustness of fine-mapping, we fine-mapped “height” from the GIANT GWAS meta-analysis and “standing height” from the UK Biobank using susieR [Wang et al., 2018]. We performed fine-mapping using susie_bhat within each LD block [Berisa and Pickrell, 2016]. We used GWAS effect sizes 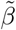 imputed from z-scores by 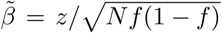 and 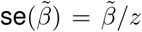, where *f* is allele frequency and *N* is GWAS sample size. The GTEx-EUR individuals were used to calculate the reference LD panel. We recorded 95% credible set which has posterior probability 95% to capture a causal signal. To compare the fine-mapping results of two GWASs, we defined their 95% credible sets as “overlapped” if they shared at least one variant. To see how 95% in GIANT GWAS is colocalized with UK Biobank GWAS, we calculated colocalization probability as 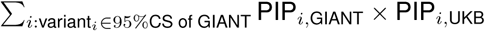.

### 1.12 Association to predicted expression or splicing

#### 1.12.1 Predicting the genetically regulated components of expression and splicing

To predict expression, we constructed linear prediction models [Barbeira et al., 2020], using only individuals of European ancestry, and variants with MAF *>* 0.01, for genes annotated as protein-coding, pseudo-gene, or lncRNA. For each gene-tissue pair, we selected the variants with highest *pip* in their cluster, and kept those achieving pip *>* 0.01 in *dap-g* [Wen, 2016]. We used *mashr* [Urbut et al., 2018] effect sizes (as computed in 1.9) for each selected variant. For each model, we computed the covariance matrix between variants using only individuals of European ancestries, with sample sizes ranging from 65 (kidney - cortex) to 602 (skeletal muscle). This allowed us to build LD panels for every tissue. For every gene, we also computed the covariance of all the variants present across the different tissue models, compiling a cross-tissue LD panel to compute the correlation between predicted expression levels across tissues. We refer to these models as *mashr* models. We compared the number of *mashr* models to the number of Elastic Net models from GTEx version 7 (fig. S8). We generated analogous prediction models for splicing ratios, as computed by Leafcutter [Li et al., 2018], applying the same model-building methodology to the data from the sQTL analysis.

Expression phenotypes were adjusted for unwanted variation using the following covariates: sex, sequencing plaform, the top 3 principal components from genotype data, and PEER factors. The number of PEER factors was determined from the sample size: 15 for *n <* 150, 30 for 150 ≤ *n <* 250, 45 for 250 ≤ *n <* 350, 60 for 350 ≤ *n*. We obtained 686,241 models for different (gene, tissue) pairs.

We also generated analogous prediction models for splicing ratios, with the same model-building methodology applied to the data from the sQTL analysis, obtaining 1,816,703 (splicing event, tissue) pairs.

**Supplementary Fig. S8.**
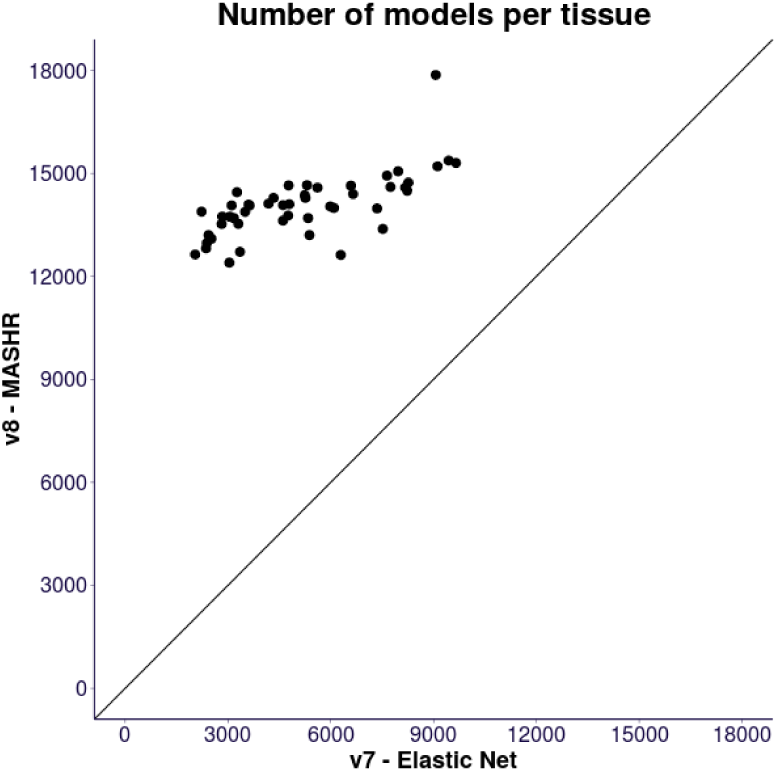
Number of models available in v8 MASHR family of models, compared to v7 Elastic Net family. The point with 17,867 models is Testis, consistently with the high levels of expression observed in the eQTL analysis Aguet et al. [2019].

#### 1.12.2 PrediXcan

We performed PrediXcan analysis [Barbeira et al., 2018] on the 87 complex traits, using the GWAS summary statistics described in 1.3.2, to identify trait-associated genes (typically *p <* 2.5 *×* 10^−7^). We used the 49 models and LD panels described in 1.12.1, separately on each trait, to obtain 59,485,548 gene-tissue-trait tuples. Repeating this process to generate splicing event ratio models, we obtained 154,891,730 splicing event-tissue-trait tuples; for each trait, the Bonferroni-significance threshold was *p <* 9.5 *×* 10^−8^.

#### 1.12.3 Colocalized and significantly associated genes

We assessed how many genes present evidence of trait association and colocalization, using both expression and splicing event. First, we counted the proportion of genes that showed a colocalized expression signal with any trait in any tissue, and observed 15% such genes at rcp*>* 0.5. Then, for each gene, we considered the splicing event with highest colocalization value in any trait or tissue, and found evidence for 5% at rcp*>* 0.5.

Then we repeated this process for PrediXcan associations at different signifcance thresholds. About 30% of genes showed a significant PrediXcan association to any trait, and only 8% when filtered for associations with rcp*>* 0.5. When using the highest splicing association and colocalization value for a gene, these proportions were 20% and 3%, respectively.

These proportions gauge our power to predict causal genes affecting complex traits on the GTEx resource, with expression yielding more findings than splicing.

**Supplementary Fig. S9.**
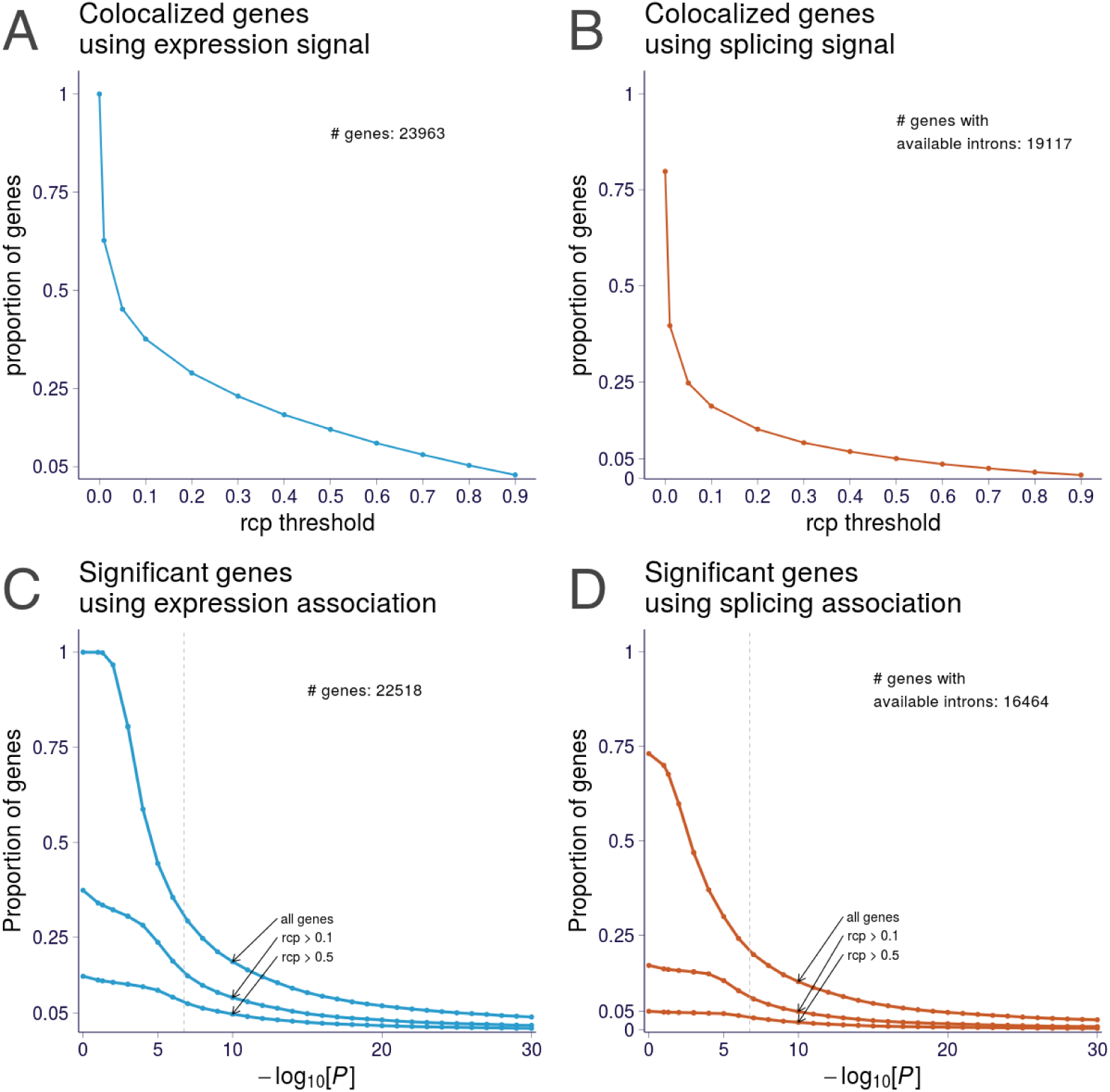
Proportion of genes with a colocalized or associated signal using expression or splicing event. **A** shows the proportion of genes with colocalization evidence in expression data, for different rcp thresholds. 3,477 genes show evidence at rcp*>* 0.5 (15% out of 23,963 genes with *enloc* results). **B** shows the proportion of genes with colocalization evidence in splicing data; 1,277 genes (5% of all 23,963) show evidence at rcp*>* 0.5. **C** shows the proportion of genes with association evidence in expression data, additionally filtered by colocalization on different thresholds. About 30% of genes show associations at the bonferroni threshold (*p <* 0.05*/*686, 241), while 8% also show colocalization evidence. **D** shows the proportion with association and colocalization evidence in splicing data; about 20% show association evidence (*p <* 0.05*/*1, 816, 703) and 3% are also colocalized.

#### 1.12.4 S-MultiXcan

Given the substantial sharing of eQTLs across tissues [GTEx Consortium et al., 2017], we aggregated PrediXcan results across tissues using S-MultiXcan [Barbeira et al., 2019]. MultiXcan has been shown to exploit the tissue sharing of regulatory variation, to improve our ability to identify trait-associated genes. The method extends the single-tissue PrediXcan approach, leveraging GWAS summary statistics and taking into account the correlation between tissues. We obtained association statistics for 1,958,220 gene-trait pairs and 11,986,329 splicing event-trait pairs.

#### 1.12.5 PrediXcan replication in BioVU

We replicated the significant gene-level associations for a prioritized list of traits (table S16) using BioVU [Denny et al., 2013], Vanderbilt University’s DNA Biobank tied to a large-scale Electronic Health Records (EHR) database. We sought BioVU replication in the exact discovery tissues for the significant gene-trait associations. We restricted our analysis to subjects of European ancestries, using principal component analysis as implemented in EIGENSOFT (version 7.1.2; [Price et al., 2006]). First, we estimated the genetically determined component of gene expression in the BioVU individuals using the PrediXcan imputation models. We then conducted association analysis for the prioritized traits using logistic regression, with sex and age as covariates.

Among replicated loci are *SORT1* (liver, coronary artery disease rcp = 0.952; dicovery p = 2.041 *×*10^−19^ BioVU p = 3.475 *×* 10^−4^), which has a well-established associations to lipid metabolism and cardiovascular traits [Musunuru et al., 2010]. Chromosome 6p24 region, which contains *PHACTR1*, has been previously associated with a constellation of vascular diseases, including coronary artery disease [Nikpay et al., 2015] and migraine headache [Anttila et al., 2013]. Notably, *PHACTR1* was significant in three different arteries (aorta artery, coronary artery and tibial artery) in two traits (coronary artery disease and migraine) in the replication analysis. In all six tissue-trait pairs, *PHACTR1* showed very high posterior probabilities in discovery analyses (rcp = 0.992 to 1.00). In our replication analysis, *PHACTR1* remained significant only for coronary artery disease associations (table S16, aorta artery, discovery p = 2.246 *×* 10^−39^, BioVU p = 7.484 *×* 10^−8^; coronary artery, discovery p = 1.952 *×* 10^−37^, BioVU p = 2.047 *×* 10^−7^; tibial artery, discovery p = 1.559 *×* 10^−33^, BioVU p = 9.880 *×* 10^−7^).

**Supplementary Fig. S10.**
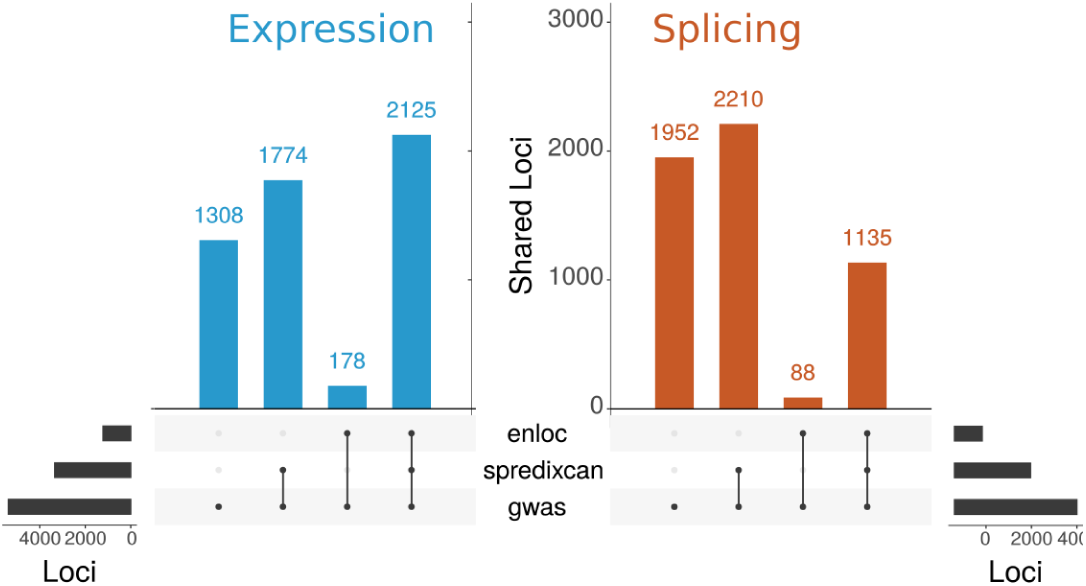
Causal gene prioritization using PrediXcan and *enloc*. Summary of GWAS loci that also contain an associated PrediXcan or colocalized signal, for expression (left) and splicing (right), using MASHR models. Significance was defined at Bonferroni-adjusted threshold for number of tests in each trait: *p <* 0.05*/*(gene-tissue pairs) = 7.28 *×* 10^−8^ for expression, *p <* 0.05*/*(intro-tissue pairs) = 2.75 *×* 10^−8^ for splicing. Colocalization status was defined as *enloc* rcp*>* 0.5.

**Supplementary Fig. S11.**
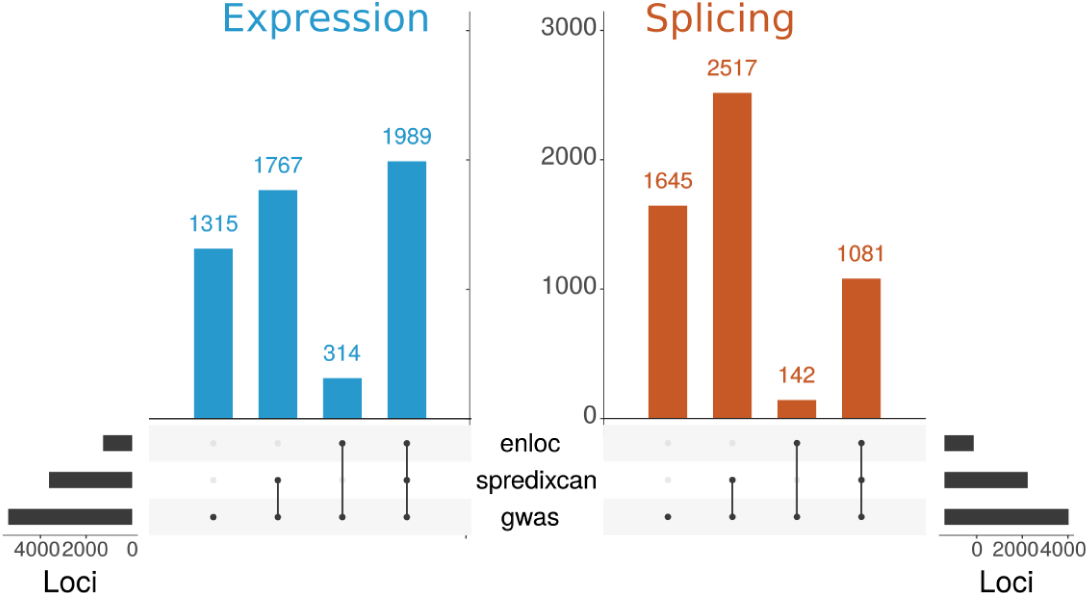
Causal gene prioritization using PrediXcan and *enloc*. Summary of GWAS loci that also contain an associated PrediXcan or *enloc* signal, for expression (left) and splicing (right), using Elastic Net models. Significance was defined at Bonferroni-adjusted threshold for number of tests in each trait: *p <* 0.05*/*(gene-tissue pairs) = 1.77 *×* 10^−7^ for expression, *p <* 0.05*/*(intro-tissue pairs) = 9.51 *×* 10^−8^ for splicing. Colocalization status was defined as *enloc* rcp*>* 0.5.

**Table S2:**
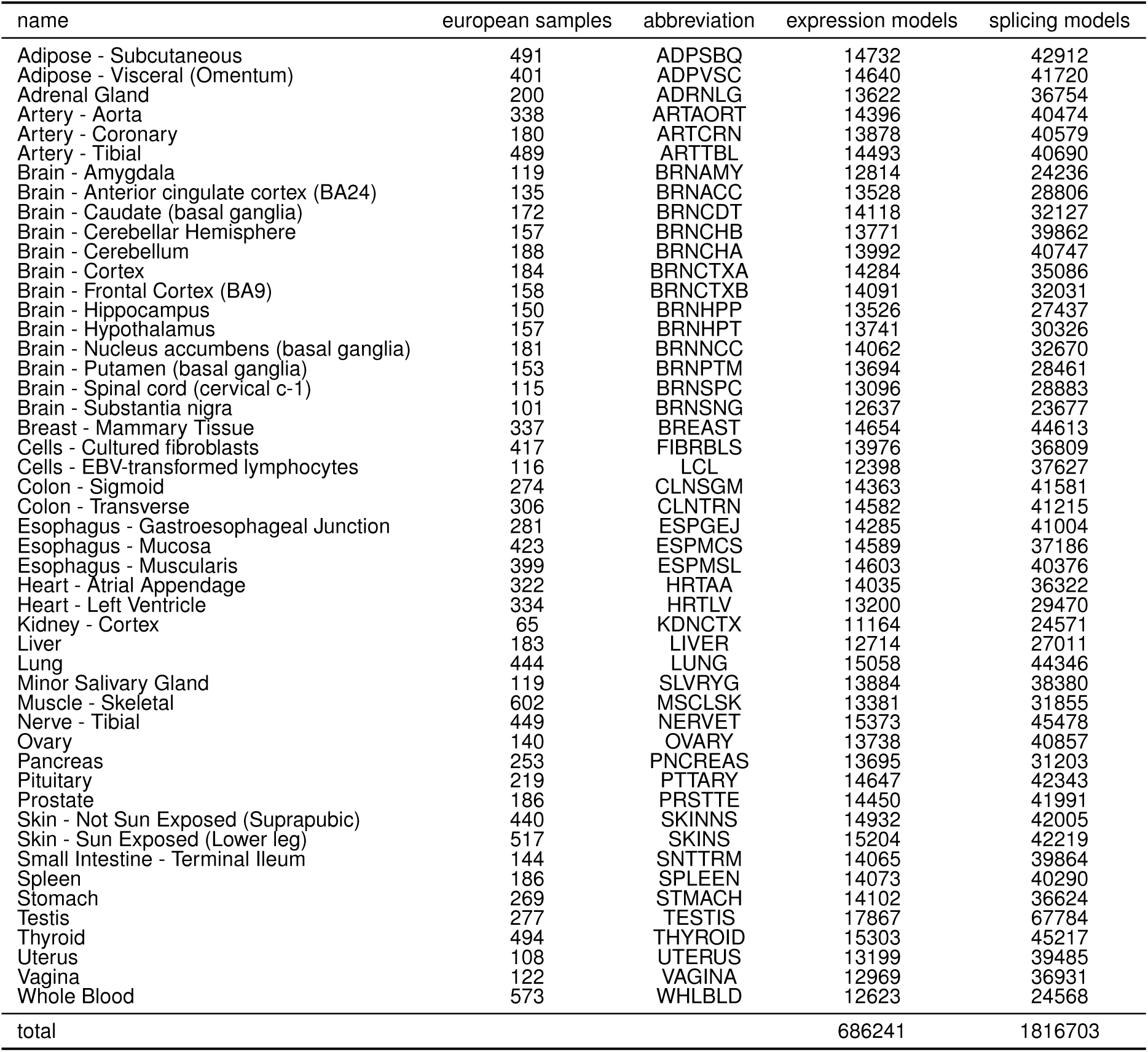
Expression and splicing prediction models using mashr-based models. Training sample size and number of genes predicted for expression and splicing traits.

**Table S3:**
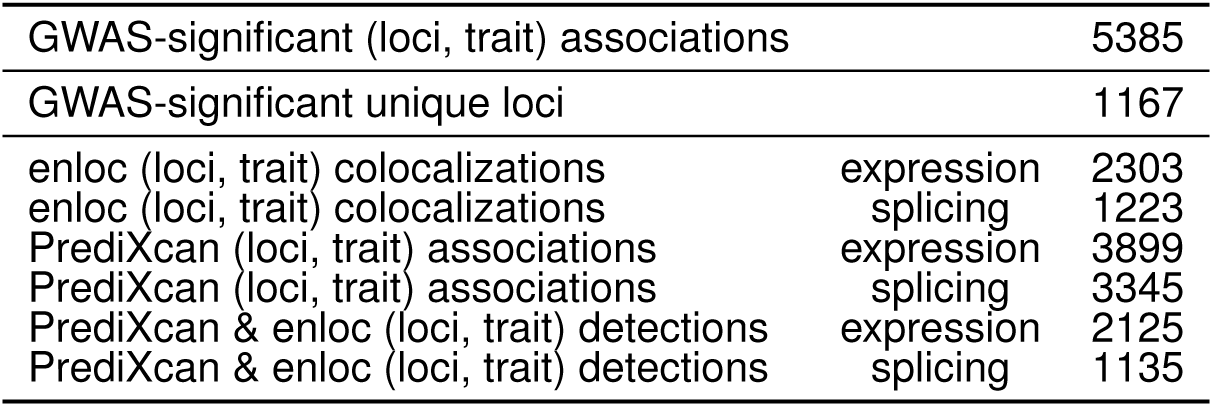
GWAS loci with colocalized or significant genes assigned. Numbers of loci-trait associations with associated/colocalized genes/splicing event detected by each method. A locus is said to have a GWAS association to a trait if it contains at least one variant with *p <* 0.05*/*(variants tested) = 5.7 *×* 10^−9^. We list here how many such loci-trait associations have an S-PrediXcan association or *enloc* signal. Significant S-PrediXcan associations were defined at Bonferroni-adjusted threshold for number of tests in each trait: *p <* 0.05*/*(gene-tissue pairs) = 7.28 *×* 10^−8^ for expression, *p <* 0.05*/*(intro-tissue pairs) = 2.75 *×* 10^−8^ for splicing. Colocalization status was defined as *enloc* rcp*>* 0.5.

#### 1.12.6 Summary-data-based Mendelian Randomization (SMR) and HEIDI

For comparison, we also performed top-eQTL based Summary-data-based Mendelian Randomization (SMR) [Zhu et al., 2016] analysis of the 4,263 tissue-trait pairs. SMR, which integrates summary statistics from GWAS and eQTL data, has been used to prioritize genes underlying GWAS associations.

**Supplementary Fig. S12.**
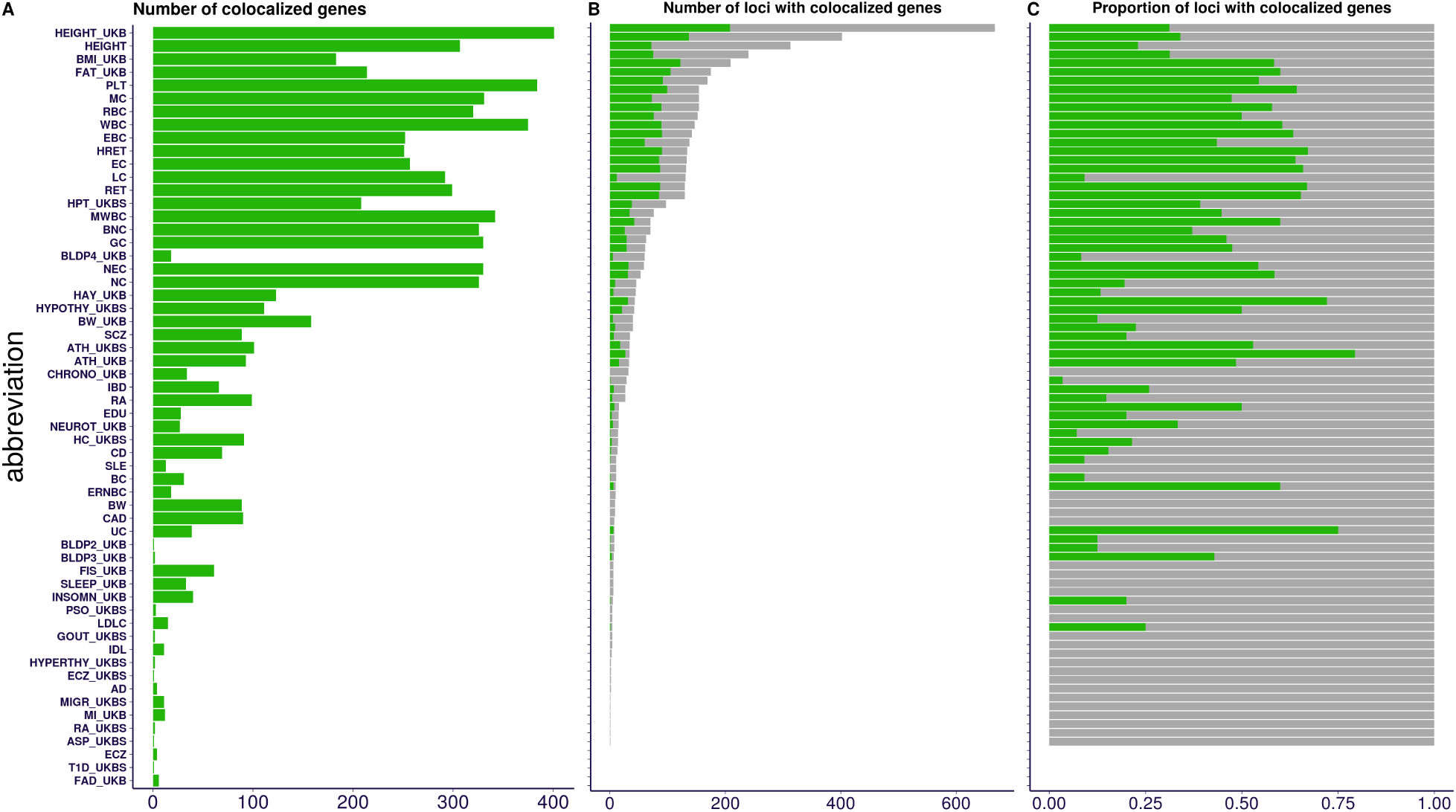
Colocalization of expression QTLs Colocalization for each of the 87 GWAS traits aggregated across the 49 tissues. GWAS loci are shown in gray, colocalized results are shown in dark green. The traits are ordered by number of GWAS-significant variants. **Panel A** shows the number of colocalized genes, achieving *enloc rcp >* 0.5 in at least one tissue, for each GWAS trait. The number of colocalized results tends to increase with the number of GWAS-significant variants. **Panel B** shows the number of loci (approximately independent LD regions from [Berisa and Pickrell, 2016]) with at least one GWAS-significant variant (dark gray), and among them those with at least one gene reaching *rcp >* 0.5 (dark green). **Panel C** shows the proportion of loci with at least one GWAS-significant hit that contain at least one colocalized gene. Across traits, a median of 21% of the GWAS loci contain colocalized results. See trait abbreviation list in table S2. These results were also presented in [Aguet et al., 2019] and are shown here for completeness.

**Supplementary Fig. S13.**
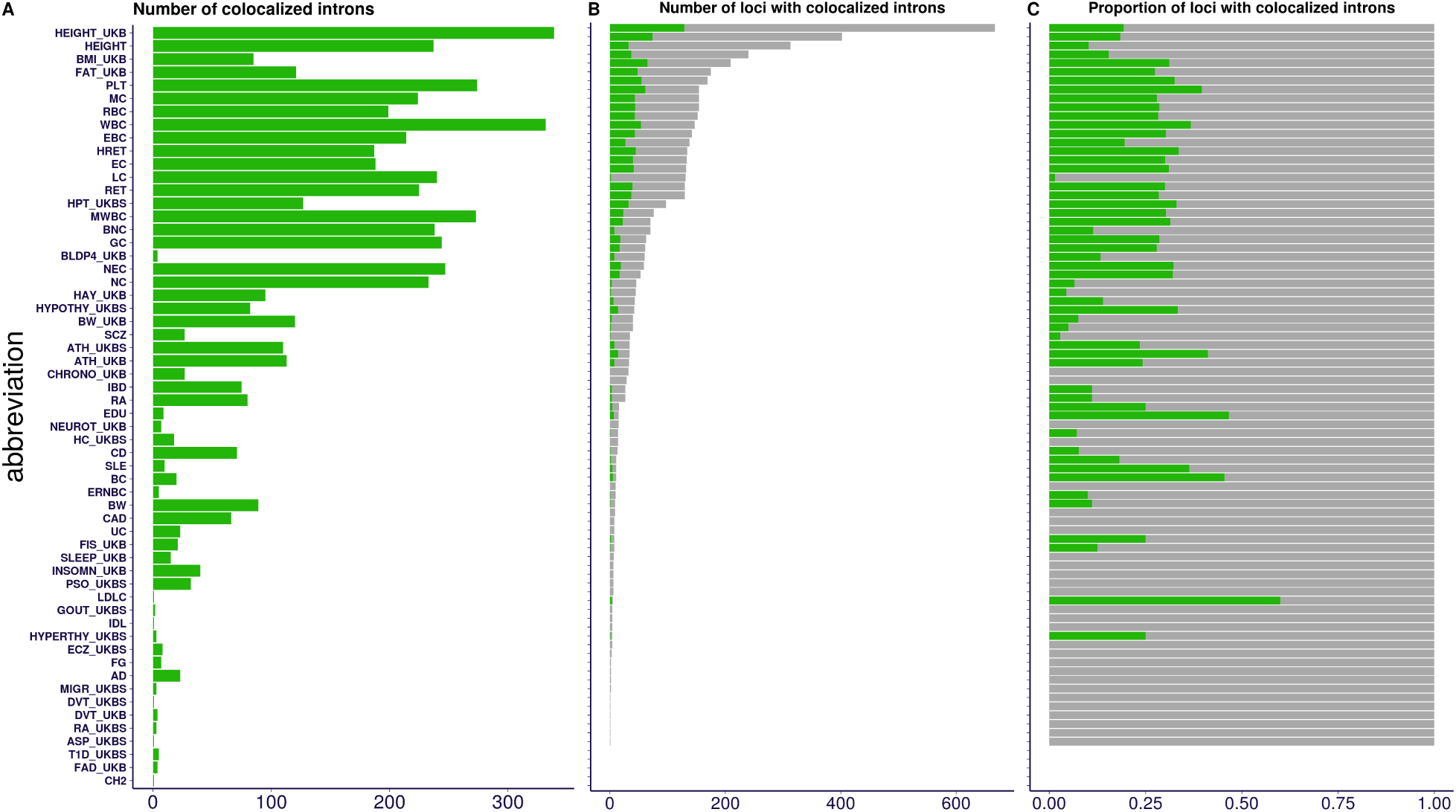
Colocalization of splicing QTLs for each of the 87 GWAS traits aggregated across the 49 tissues. The traits are ordered by number of GWAS-significant variants. GWAS loci are shown in gray, colocalized results are shown in dark green. **Panel A** shows the number of colocalized splicing event, achieving *enloc rcp >* 0.5 in at least one tissue, for each GWAS trait. As with gene expression results, the number of colocalized results tends to increase with the number of GWAS-significant variants. **Panel B** shows the number of loci (approximately independent LD regions from [Berisa and Pickrell, 2016]) with at least one GWAS-significant variant (dark gray), and among them those with one splicing event achieving *rcp >* 0.5 (dark green). **Panel C** shows the proportion of loci with at least one GWAS-significant hit loci with at least one colocalized splicing event. Across traits, a median of 11% of the GWAS loci contain a colocalized result, lower than the gene expression counterpart (29%), indicating a decreased power in the sQTL study. See trait abbreviation list in table S2.

**Supplementary Fig. S14.**
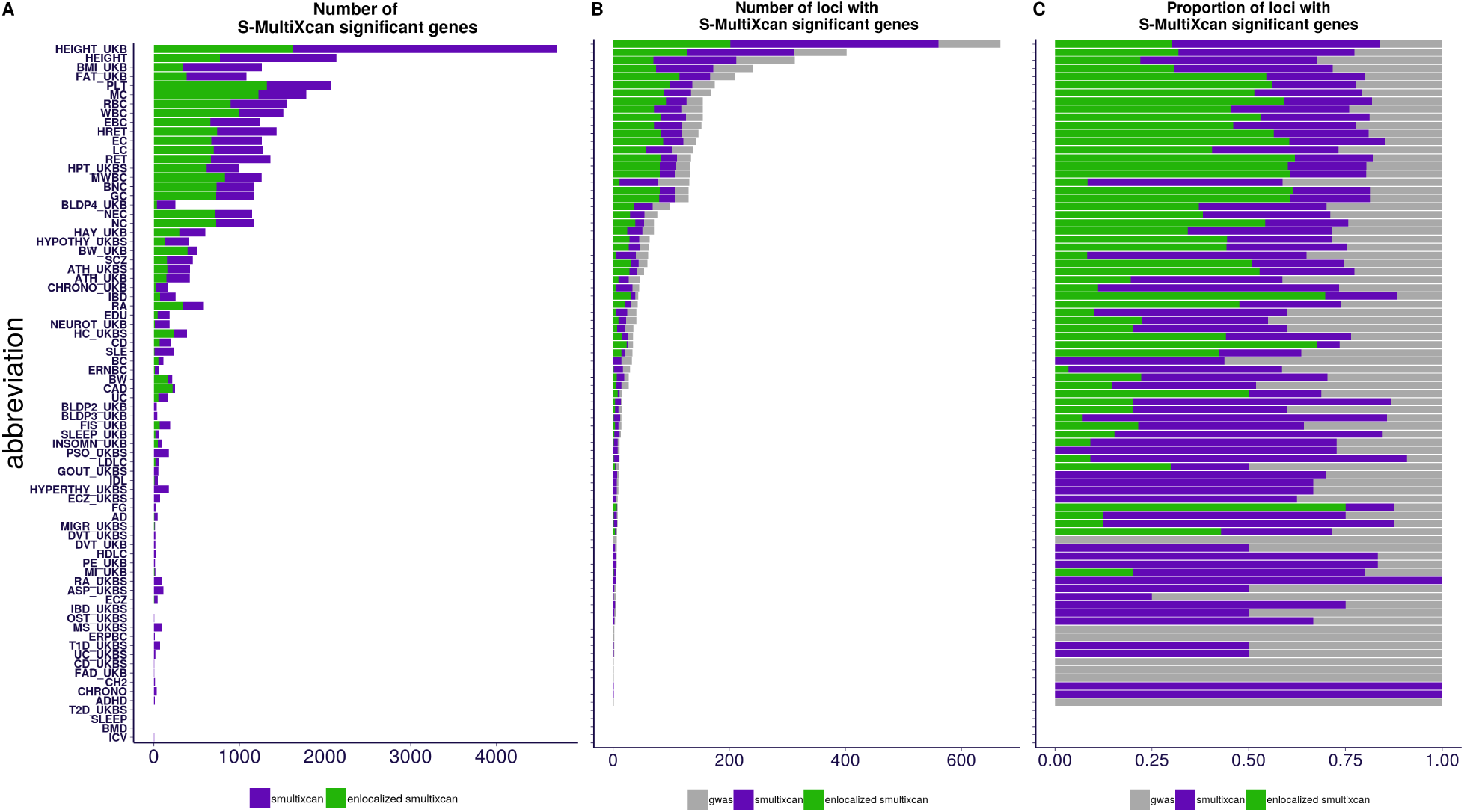
PrediXcan expression associations aggregated across tissues. This figure summarizes S-MultiXcan associations for each of the 87 traits using the gene expression models. The traits are ordered by number of GWAS-significant variants. **Panel A)** shows in purple the number of S-MultiXcan significant genes, and in dark green the subset also achieving *enloc rcp >* 0.5 in any tissue. S-MultiXcan has a high power for detecting associations, but 12% (median across traits) of these genes show evidence of colocalization. **Panel B)** shows the number of loci (approximately independent LD regions [Berisa and Pickrell, 2016]) with a significant GWAS association (gray), a significant S-MultiXcan association (purple), and a significant S-MultiXcan association that is colocalized (dark green). Anthropometric and Blood traits tend to present the largest number of associated loci, with Height from two independent studies leading the number of associations. **Panel C)** shows the proportion of loci with significant GWAS associations (gray) that contain S-Multixcan (purple) and colocalized S-MultiXcan associations (dark green). Across traits, a median of 70% of GWAS-associated loci show a S-MultiXcan detection, while 19% show a colocalized S-MultiXcan detection. See trait abbreviation list in table S2.

**Supplementary Fig. S15.**
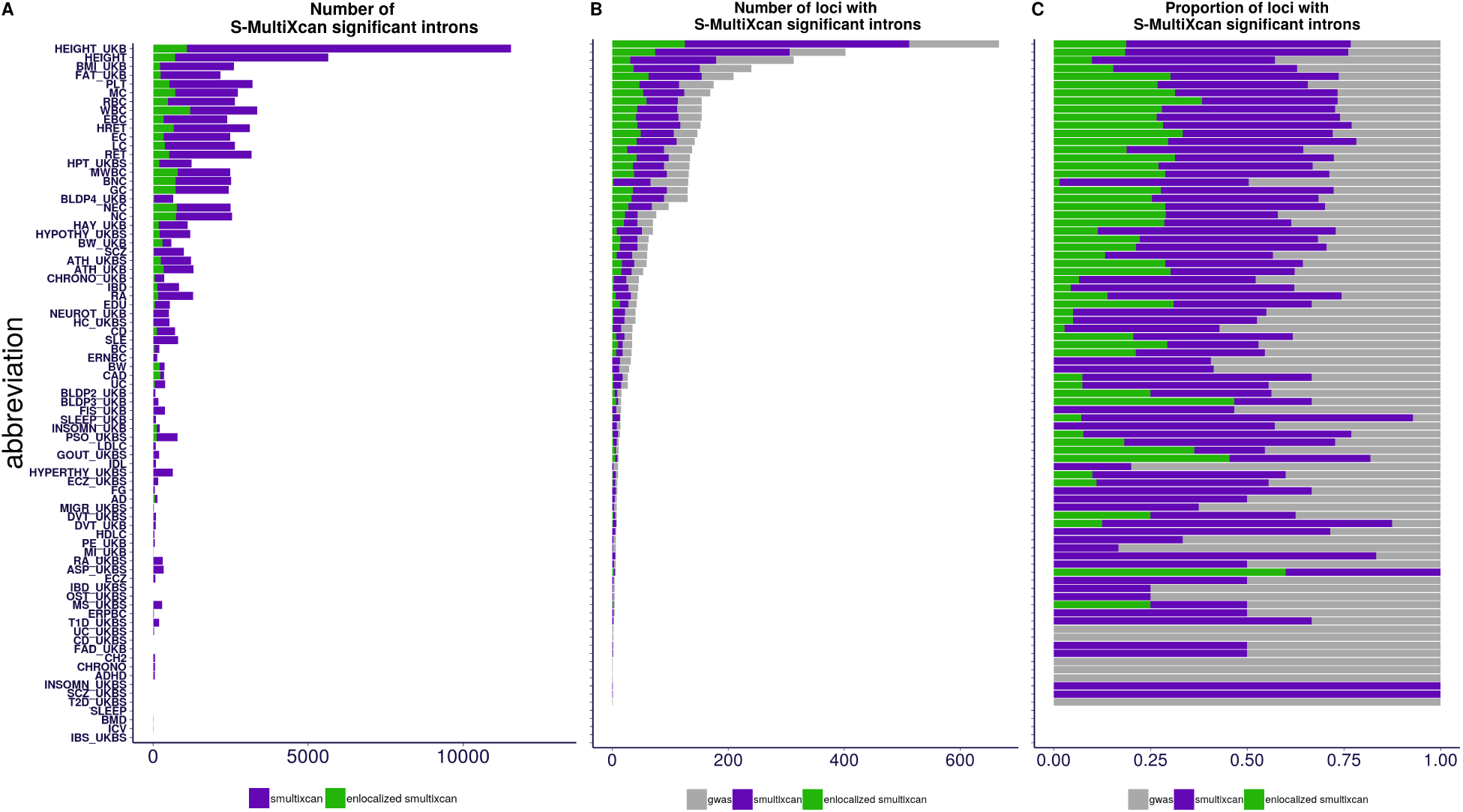
PrediXcan splicing associations aggregated across tissues. This figure summarizes S-MultiXcan associations for each of the 87 traits using splicing models. The traits are ordered by number of GWAS-significant variants. Panel A) shows in purple the number of S-MultiXcan significant splicing events, and in dark green the subset also achieving *enloc rcp >* 0.5 in any tissue. The proportion of colocalized, significantly associated splicing events is typically 2%, much lower than the proportion from gene expression (12%). Panel B) shows the number of loci (approximately independent LD regions [Berisa and Pickrell, 2016]) with a significant GWAS association (gray), a significant S-MultiXcan association (purple), and a significant S-MultiXcan association that is colocalized (dark green). As in the case of expression models, Anthropometric and Blood traits tend to present the largest number of associated loci. Panel C) shows the proportion of loci with significant GWAS associations (gray) that contain S-Multixcan (purple) and colocalized S-MultiXcan associations (dark green). Across traits, a median of 63% of GWAS-associated loci show an S-MultiXcan association, while 11% show a colocalized S-MultiXcan association. These proportions are lower than the corresponding ones for expression (70% and 19% respectively). See trait abbreviation list in table S2.

### 1.13 Assessing the performance of association and colocalization methods to identify causal genes

To assess the performance of colocalization and association methods to identify causal genes, we curated two sets of ‘causal’ gene-trait pairs. One set is based on the OMIM database and the other one is based on rare variant association results from exome-wide association studies. To quantify the performance, we framed the causal gene identification problem as one of classification and used the standard tools such as ROC and precision recall curves, which have the advantage of not needing ad-hoc thresholds and show the full trade-off between true positives and false positives as well as precision vs. power. Throughout this section, we limited our scope to only the protein-coding genes.

#### 1.13.1 OMIM-based curation of causal genes

To obtain a curated set of trait-gene pairs from the OMIM database [Hamosh et al., 2005], we mapped our GWAS traits to the OMIM traits and linked them to the corresponding genes in the OMIM database. The mapping process is illustrated in fig.S16 for a specific example GWAS trait, fasting glucose by the MAGIC consortium. First, the GWAS trait was mapped to the GWAS catalog trait names by searching for relevant keywords (defined manually S5) in the description field of the GWAS catalog. Second, the GWAS catalog trait names were linked to phecodes using the mapping in the phewas catalog [Denny et al., 2013]. Third, we mapped phecodes to OMIM traits ids (MIM) as described in [Bastarache et al., 2018]. Finally, in step 4, we mapped OMIM traits to OMIM genes using the

The keywords used for the each of the initial selected set of 114 GWAS traits is listed in tab S5). For a subset of datasets with GWAS results from more than one source (public GWAS vs UKB) in our collection, we kept the dataset with higher number of GWAS loci to avoid double counting. The number of GWAS loci was determined based on counting the lead variants, using the PLINK V1.9 command `–clump-r2 0.2 –clump-p1 5e-8` at genome-wide significance 5 *×* 10^−8^) for each trait. Furthermore, for this analysis, we excluded GWAS traits with fewer than 50 GWAS loci. The full list of OMIM based trait-gene pairs is listed in S10.

With this procedure, we curated a list of 1,592 gene-trait pairs with evidence of causal associations in the OMIM database (hereafter, **OMIM genes**), which was downloaded from omim.org/downloads (accessed on Aug 12th 2019). After matching traits, we retained 29 unique traits and 631 unique genes that were within the same LD block [Berisa and Pickrell, 2016] as the GWAS hit (table S10).

**Supplementary Fig. S16.**
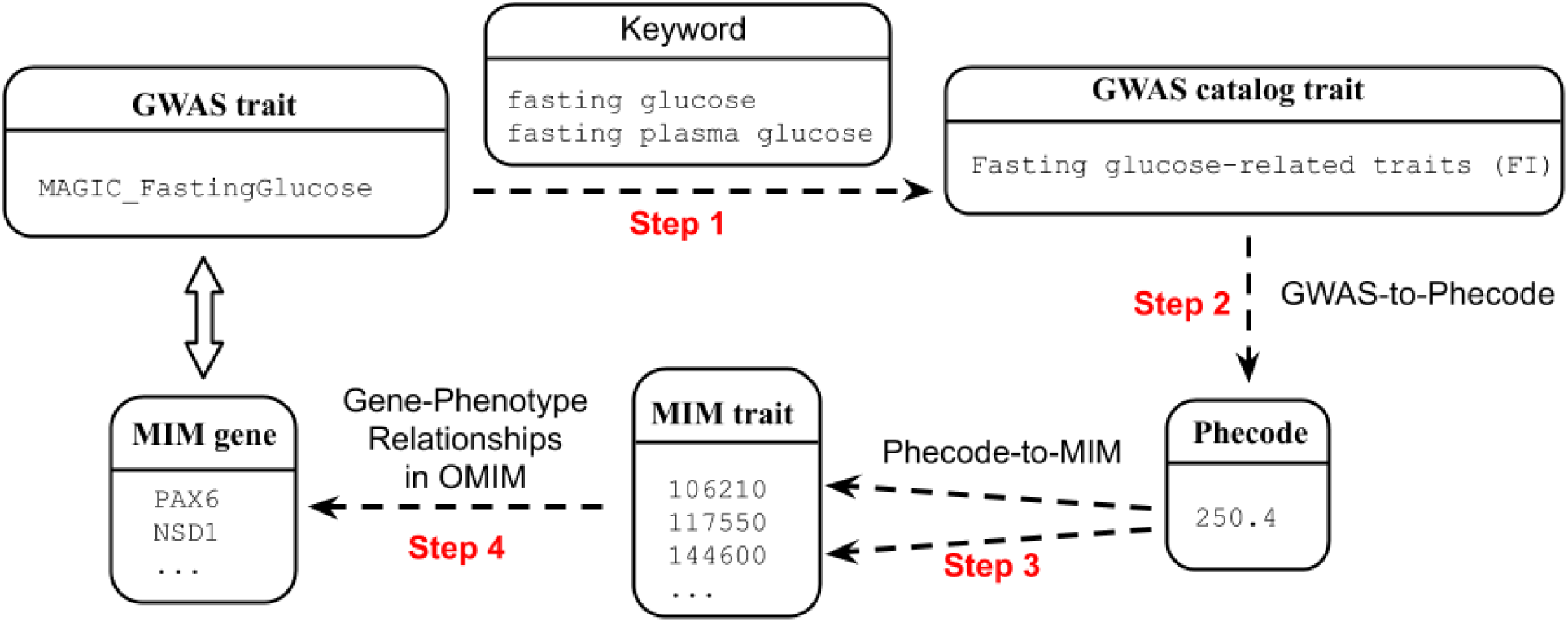
Workflow of OMIM-based curation of causal genes. The workflow of OMIM-based causal gene curation is shown where each box represents the trait description/identifier in different databases. The steps to obtain OMIM genes for MAGIC_FastingGlucose, one of our GWAS traits, is shown as a concrete application of the workflow.

**Table S4:**
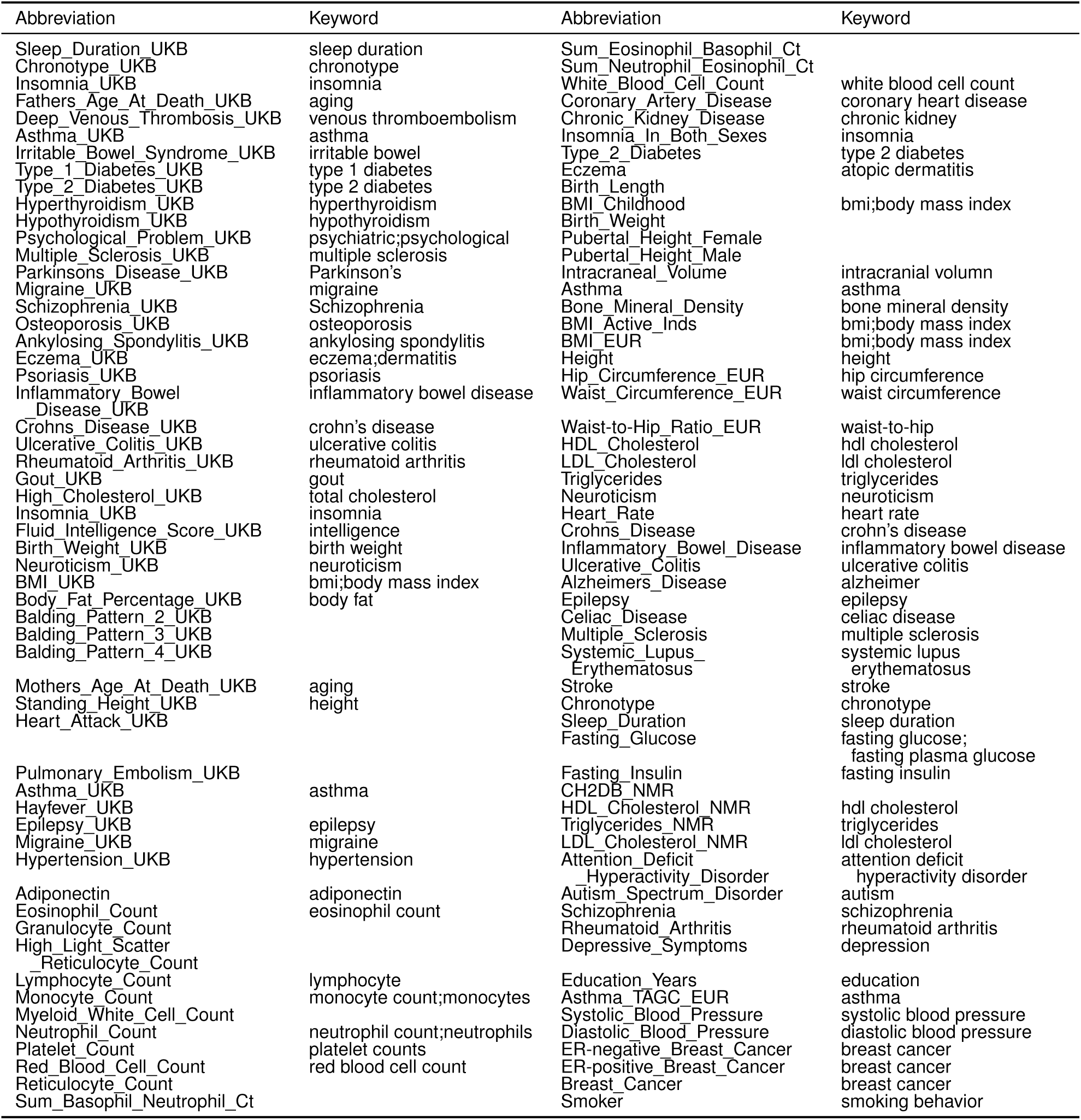
Keywords of GWAS traits used for mapping with the GWAS catalog. Keywords of all 114 GWAS traits used for OMIM-based curation and analyses are listed.

#### 1.13.2 Rare variant association-based curation of causal genes

In addition to the OMIM-based curation, we collected a set of genes in which rare protein-coding variants were reported to be significantly associated with our list of complex traits. Given the power of existing rare variant association studies, we focused on height and lipid traits (low-density lipid cholesterol, high-density lipid cholesterol, triglycerides, and total cholesterol levels) [Marouli et al., 2017; Liu et al., 2017; Locke et al., 2019].

We collected significant coding/splicing variants reported previously [Marouli et al., 2017] and kept variants with effect allele frequency *<* 0.01 (table S6 therein: ExomeChip variants with Pdiscovery <2e-07 in the European-ancestry meta-analysis (N=381,625)). Similarly, we collected significant variants reported by [Liu et al., 2017] (table S12 therein: Association Results for 444 independently associated variants with lipid traits) and filtered out variants with minor allele frequency *<* 0.01. For the whole-exome sequencing study conducted in Finnish isolates [Locke et al., 2019], we extracted significant genes identified by a gene-based test using protein truncating variants (table S9 therein: Gene-based associations from aggregate testing with EMMAX SKAT-O with P<3.88E-6) and significant variants (table S7 therein: A review of all variants that pass unconditional threshold of P<5E-07 for at least one trait) with gnomAD MAF *<* 0.01. The full list of trait-gene pairs constructed from the process is available in table S13.

#### 1.13.3 Setting up the classification problem to quantify performance for identifying causal genes

We partitioned the genome into approximately independent LD blocks [Berisa and Pickrell, 2016] and for each GWAS trait, we kept only genes located in LD blocks where there were at least one silver standard gene and a GWAS significant hit for the trait as illustrated in fig. S18. Then, we labelled the silver standard genes as 1 and all the others were labelled as 0, as represented schematically in fig. S19. We calculated the ROC and precision recall curves for classifying the silver standard gene correctly.

In more detail, for each of tested gene-trait pairs, we obtained the gene-level statistics for the corresponding trait from the application of various methods, *i*.*e. enloc, coloc*, SMR, and PrediXcan-*mashr*. Since we had results across tissues, we selected the ‘best’ scores (highest regional colocalization probability (rcp) in *enloc*; highest posterior probability under hypothesis 4 in *coloc*; smallest p-value in SMR and PrediXcan-*mashr*) to build the table S19. For splicing (with statistics reported at the intron excision event level), we obtained gene-level statistics by taking the ‘best’ score among all splicing events of the gene, across all tissues.

The full list of silver standard genes can be found in tables S14 and S15. The number of GWAS loci and silver standard genes that remained after the above filtering steps can be found in table S6. The number of genes tested per LD block is shown in fig. S17.

**Table S5:** Count of GWAS loci with predicted causal effects overlapping likely functional genes. The number of GWAS loci and the number of silver standard genes included for analysis after taking the intersection between GWAS loci and silver standard genes are shown.

| silver standard | trait | nloci | ngene | silver standard | trait | nloci | ngene |
| --- | --- | --- | --- | --- | --- | --- | --- |
| rare variant | Standing_Height_UKB | 29 | 35 | OMIM | Monocyte_Count | 1 | 1 |
| rare variant | LDL_Cholesterol | 7 | 10 | OMIM | Neutrophil_Count | 14 | 17 |
| rare variant | High_Cholesterol_UKBS | 6 | 8 | OMIM | White_Blood_Cell_Count | 16 | 17 |
| rare variant | HDL_Cholesterol | 12 | 18 | OMIM | Coronary_Artery_Disease | 12 | 13 |
| rare variant | Triglycerides | 6 | 9 | OMIM | Type_2_Diabetes | 11 | 12 |
| OMIM | Deep_Venous_Thrombosis | 2 | 2 | OMIM | Waist_Circumference_EUR | 6 | 6 |
| OMIM | Asthma_UKBS | 10 | 12 | OMIM | LDL_Cholesterol | 7 | 9 |
| OMIM | Type_1_Diabetes_UKBS | 1 | 2 | OMIM | Triglycerides | 11 | 11 |
| OMIM | Hypothyroidism_UKBS | 14 | 14 | OMIM | Inflammatory_Bowel_Disease | 7 | 8 |
| OMIM | Eczema_UKBS | 4 | 5 | OMIM | Ulcerative_Colitis | 4 | 4 |
| OMIM | Psoriasis_UKBS | 2 | 2 | OMIM | Alzheimers_Disease | 2 | 2 |
| OMIM | Gout_UKBS | 1 | 1 | OMIM | Systemic_Lupus_Erythematosus | 3 | 5 |
| OMIM | High_Cholesterol_UKBS | 6 | 8 | OMIM | Schizophrenia | 1 | 1 |
| OMIM | BMI_UKB | 35 | 35 | OMIM | Rheumatoid_Arthritis | 3 | 3 |
| OMIM | Hypertension_UKBS | 19 | 24 | OMIM | Systolic_Blood_Pressure | 2 | 2 |
| OMIM | Eosinophil_Count | 7 | 7 | OMIM | Diastolic_Blood_Pressure | 3 | 3 |
| OMIM | Lymphocyte_Count | 2 | 2 |  |  |  |  |

**Supplementary Fig. S17.**
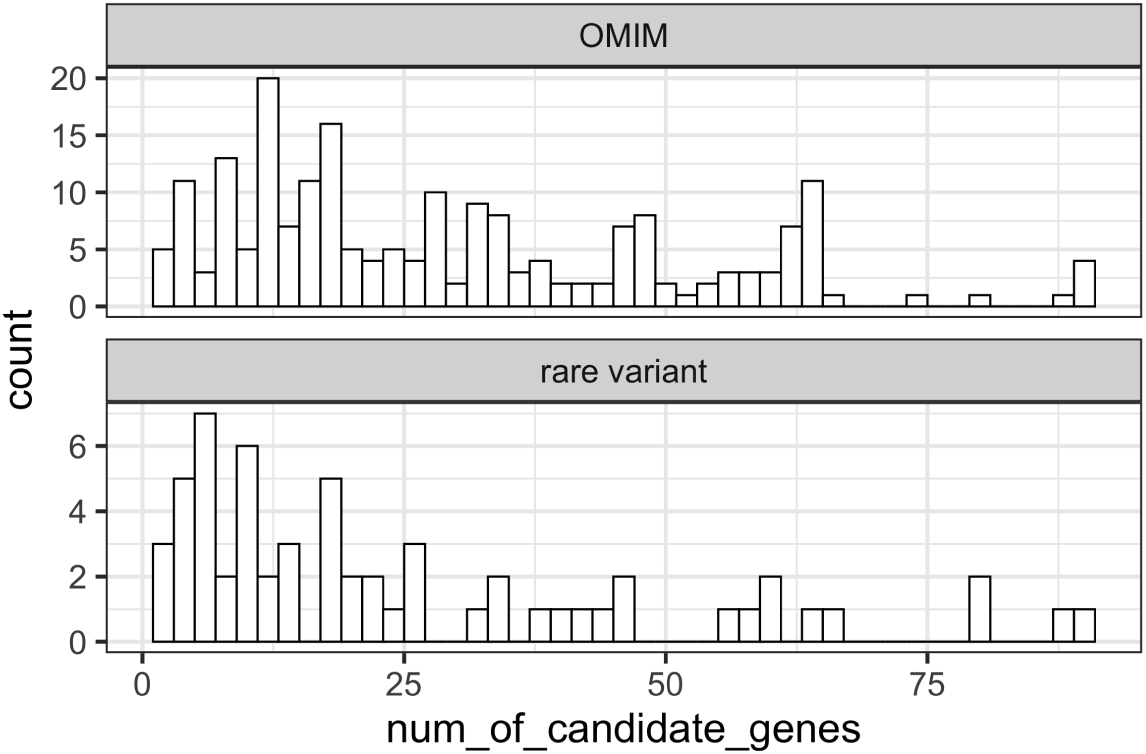
Distribution of the number of tested genes per GWAS locus overlapping OMIM- and rare variant-based silver standard. The distributions of the number of candidate genes per GWAS locus are shown for OMIM-based curation (top) and rare variant association-based curation (bottom).

**Supplementary Fig. S18.**
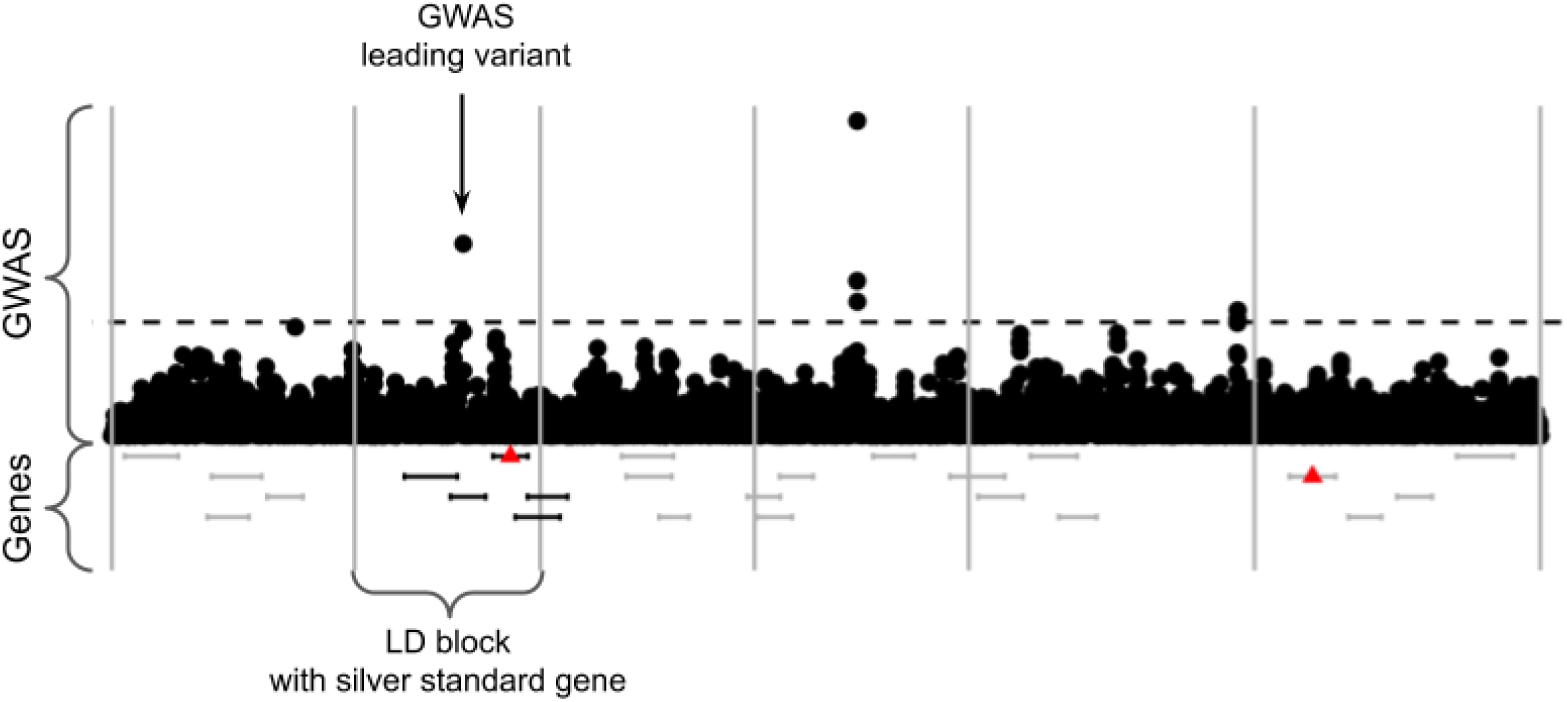
Selection of genes to assess ability to identify silver standard genes. The GWAS summary statistics were binned into independent LD blocks (boundaries of LD block are shown as gray vertical lines). Only genes within LD blocks that contain both a silver standard gene (red triangle) and a GWAS significant variant (points above −log10(*p*) *>* −log10(5 10^−8^)) were used in the calculation of performance (ROC and PR curves).

**Supplementary Fig. S19.**
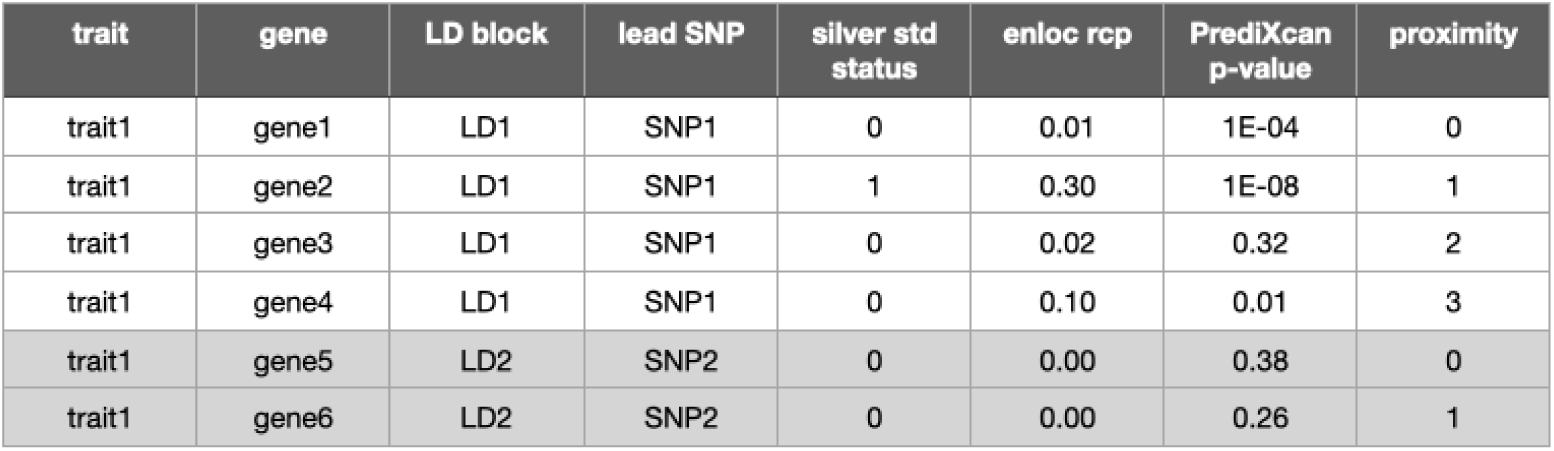
Schematic representation of data used for classification.

#### 1.13.4 AUC of the ROC curves

For expression, the areas under the curve (AUC) of were, in increasing performance, 0.553, 0.591, 0.669, and 0.672 for *coloc*, SMR, *enloc*, and PrediXcan using the OMIM silver standard 3C. AUC were higher when using the rare variant silver standard with SMR at the bottom of the ranking followed by *coloc*, PrediXcan, and enloc at the top S7. For splicing *enloc* had higher 0.650 vs. 0.632 for PrediXcan using OMIM silver standard and 0.714 and 0.686 using the rare variant silver standard.

**Table S6:** Enrichment and AUC fo *coloc, enloc*, SMR, and PrediXcan

| Regulation | Dataset | Method | ROC AUC |
| --- | --- | --- | --- |
| expression | OMIM | *coloc* | 0.553 |
| expression | OMIM | *enloc* | 0.669 |
| expression | OMIM | PrediXcan | 0.672 |
| expression | OMIM | SMR | 0.591 |
| expression | Rare variant | *coloc* | 0.661 |
| expression | Rare variant | *enloc* | 0.755 |
| expression | Rare variant | PrediXcan | 0.743 |
| expression | Rare variant | SMR | 0.629 |
| splicing | OMIM | *enloc* | 0.650 |
| splicing | OMIM | PrediXcan | 0.632 |
| splicing | Rare variant | *enloc* | 0.714 |
| splicing | Rare variant | PrediXcan | 0.686 |

#### 1.13.5 Precision-recall curves of PrediXcan and *enloc* on silver standard gene sets

**Supplementary Fig. S20.**
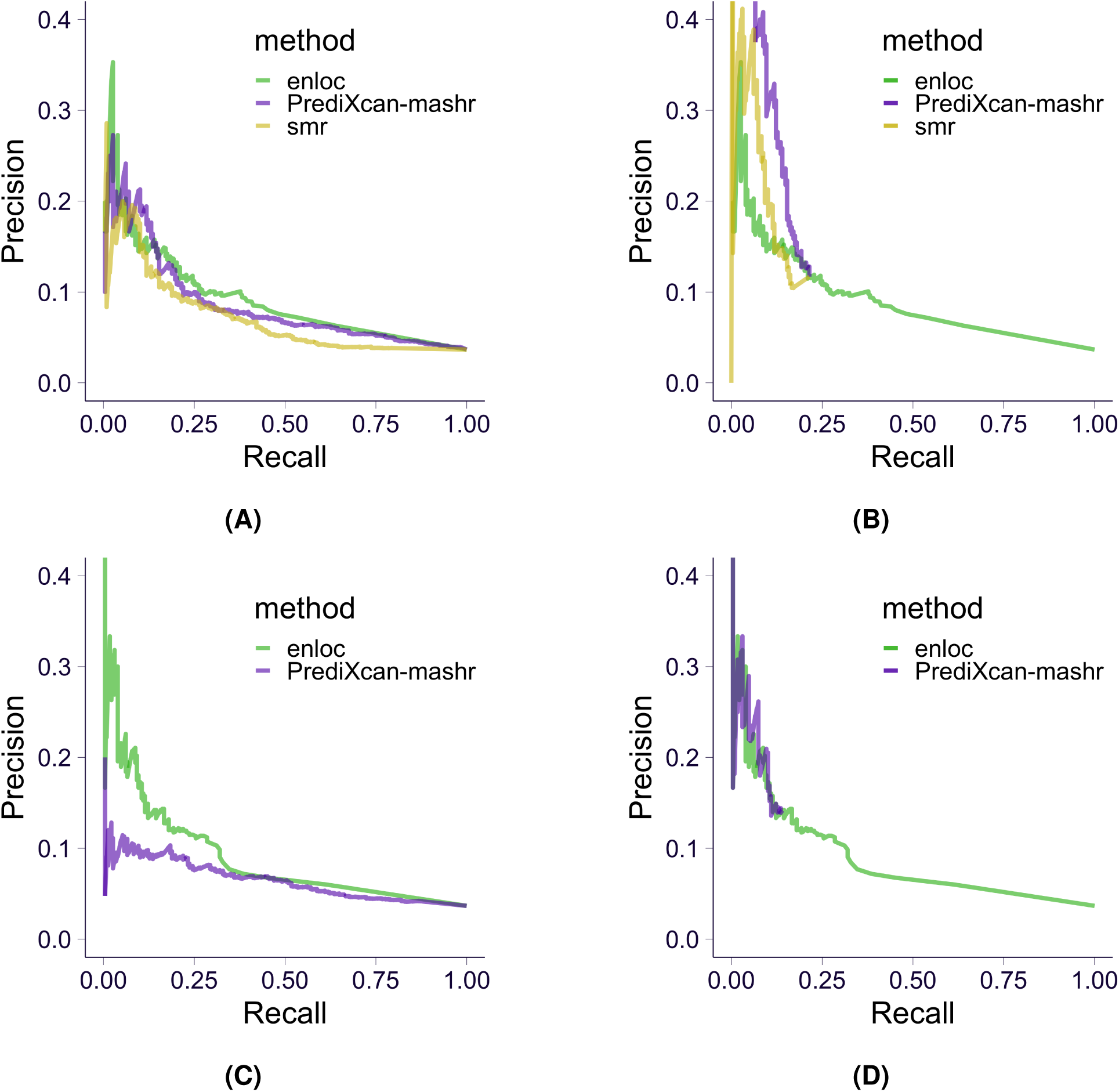
Precision-recall curves of colocalization/association based methods on OMIM silver standard. The results on expression data are shown in top row and the ones on splicing data are shown in bottom row. **(A**,**C)** Precision-recall curve of colocalization/association based methods. **(B**,**D)** Precision-recall curve of association based methods when pre-filtering with *enloc* rcp > 0.1.

**Supplementary Fig. S21.**
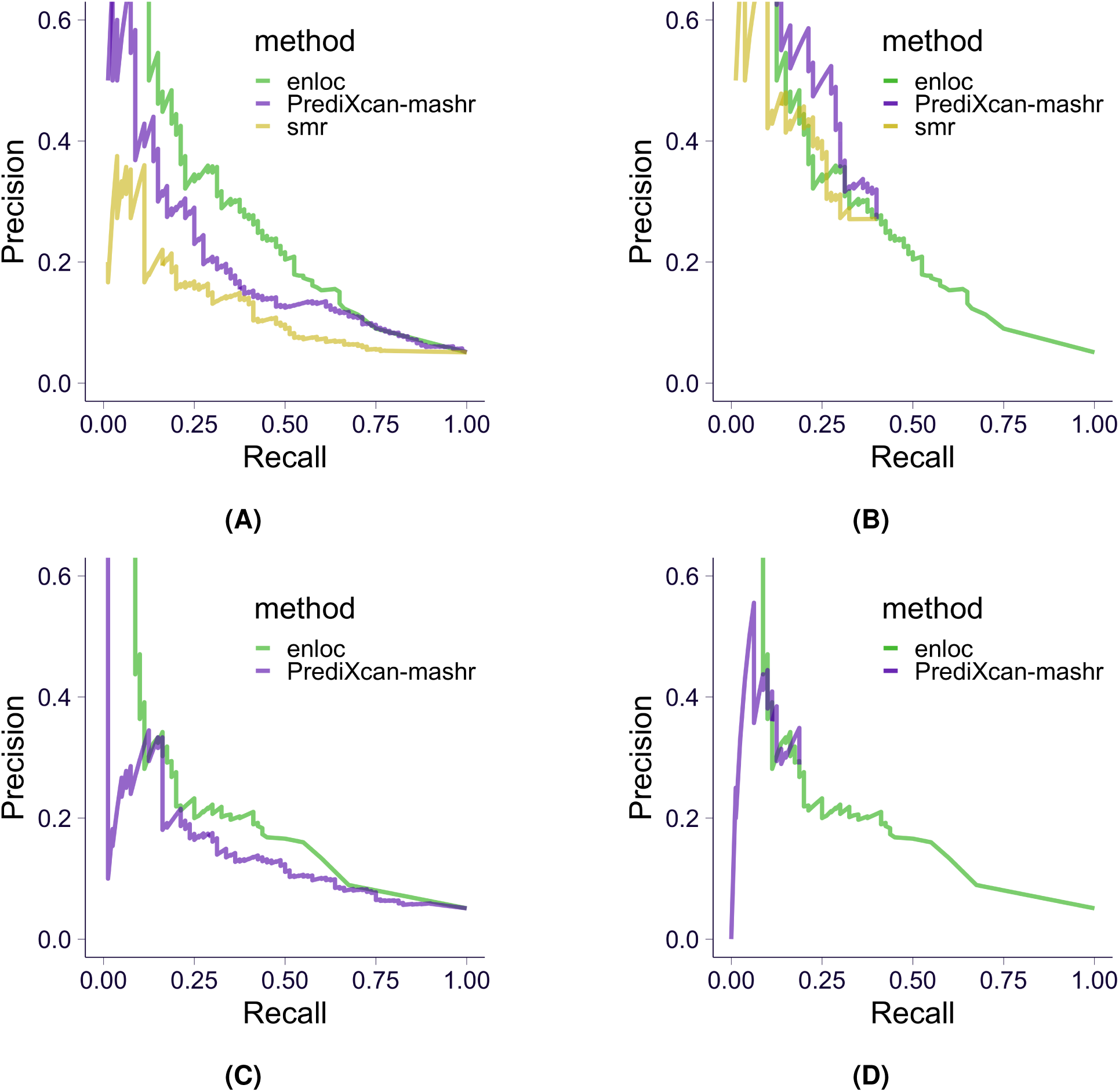
Precision-recall curves of colocalization/association based methods on rare variant-based silver standard. The results on expression data are shown in top row and the ones on splicing data are shown in bottom row. **(A**,**C)** Precision-recall curve of colocalization/association based methods. **(B**,**D)** Precision-recall curve of association based methods when pre-filtering with *enloc* rcp > 0.1.

**Supplementary Fig. S22.**
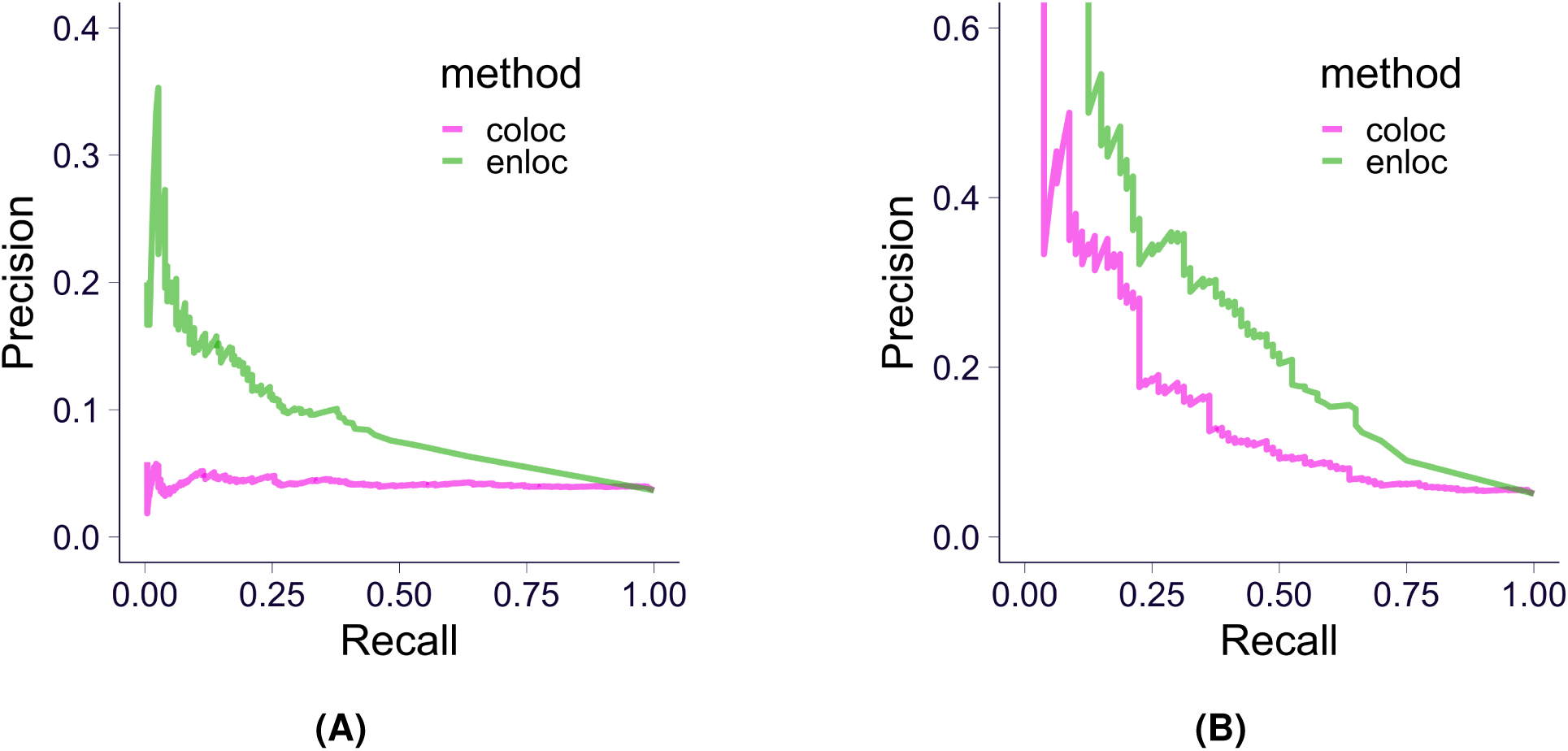
Precision-recall curves of *enloc* vs *coloc*. Precision recall curve of *enloc* (blue) and *coloc* (green) with expression using OMIM silver standard (in **(A)**) and rare variant-based silver standard (in **(B)**).

#### 1.13.6 Assessing the contribution of proximity, colocalization, and association significance

To investigate the usefulness of the colocalization and association statistics reported by *enloc* and PrediXcan respectively, we performed logistic regression, as described in Eq. 28, to fit log odds of being a ‘causal’ gene against the ranking of: 1) proximity to GWAS lead variant (from close to distal), 2) rcp from *enloc* (from high to low), and 3) gene-level association p-value from PrediXcan-*mashr* or SMR (from significant to non-significant).

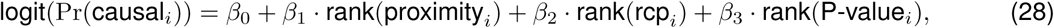

in which non-zero *β*_*k*_ meant that the *k*th variable contributed independently on predicting whether a gene was causal. Moreover, negative *β*_*k*_ indicated that the direction of contribution of the variable was as expected.

We note that here the analysis is performed by LD blocks rather than genome-wide as was done for calculating the ROC and precision recall curves. More specifically, the ranking within each LD block is used rather than genome-wide.

**Table S7:**
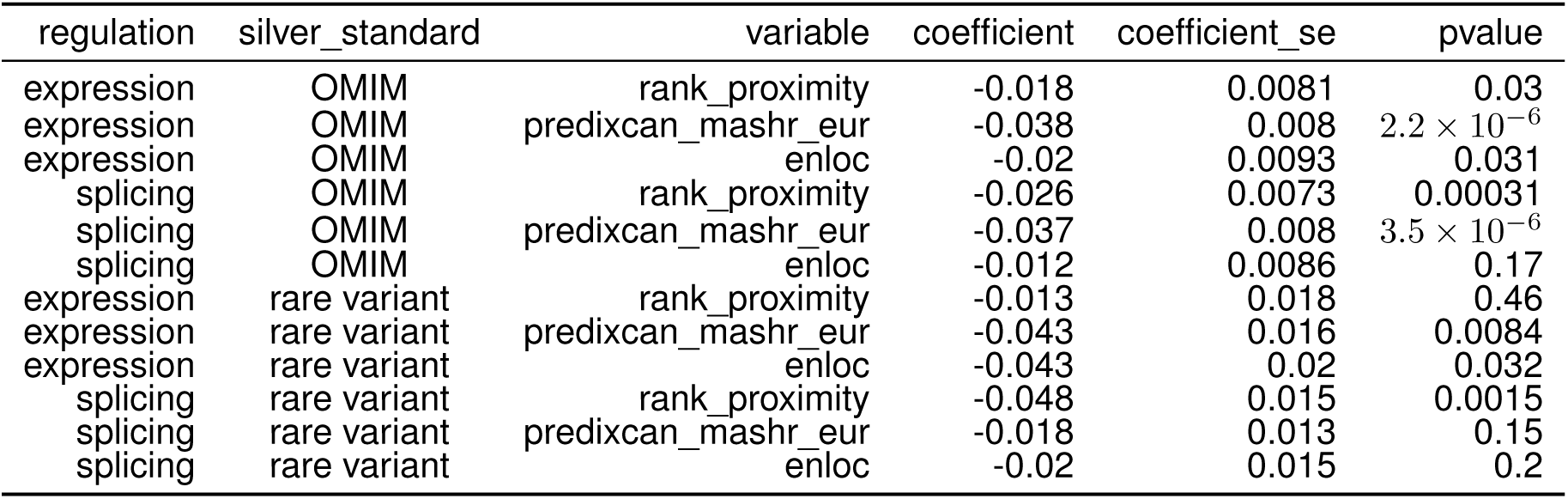
Predictive value of different per-locus prioritization methods. Results on regression-based test (logistic regression) in per-locus analysis are shown. The estimated log odds ratio of the rank of proximity (distance between GWAS leading variant and gene body), PrediXcan significance, and *enloc* rcp are shown in rows **rank_proximity, predix-can_mashr_eur**, and **enloc**.

### 1.14 Causal tissue analysis

To identify tissues of relevance for the etiology of complex traits, we investigated the patterns of tissue specificity and tissue sharing of PrediXcan association results across 49 tissues. For each trait-gene pair, the PrediXcan z-score can be represented as a 49 *×*1 vector with each entry being the gene-level z-score in the corresponding tissue (if the prediction model of the gene is not available in that tissue, we filled in zero). To explore the tissue-specificity of the PrediXcan z-score vector, we proceeded by assigning the z-score vector to a tissue-pattern category and tested whether certain tissue-pattern categories were over-represented among colocalized PrediXcan genes as compared to non-colocalized genes. We used the FLASH factors identified from matrix factorization applied to the cis-eQTL effect size matrix, as described in Section 1.9 (as PrediXcan and cis-eQTL shared similar tissue-sharing pattern, data not shown). To obtain a set of detailed and biologically interpretable tissue-pattern categories from the 31 FLASH factors, we manually merged them into 18 categories as shown in fig. S23. For each trait, we projected the z-score vector of each gene to one of the 31 FLASH factors (as described in Section 1.9) so that the gene was assigned to the corresponding tissue-pattern category. We defined a ‘positive’ set of genes as the ones that met Bonferroni significance at *α* = 0.05 in at least one tissue and *enloc* rcp > 0.01 in at least one tissue, which could be thought as a set of candidate genes affecting the trait through expression level. We chose a rather low threshold used for the rcp due to the stringent conservative nature of colocalization probabilities. We also constructed a ‘negative’ set of genes with *enloc* rcp = 0, which could be thought as a set of genes whose expressions were unlikely to affect the trait. We proceeded to test whether certain tissue-pattern categories were enriched in ‘positive’ set as compared to ‘negative’ set. Since the main focus of this analysis was tissue-specific patterns, we excluded *Factor1* (the cross-tissue factor) and *Factor25* (likely to be a tissue-shared factor capturing tissues with large sample size). Additionally, we excluded *Factor7* (testis), as it was unlikely to be the mediating tissue but might introduce false positives. We tested the enrichment of each tissue-pattern category by Fisher’s exact test (‘positive’/’negative’ sets and in/not in tissue-patter category). Among 87 traits, 82 traits had *enloc* signal and the enrichment of these was calculated accordingly.

**Supplementary Fig. S23.**
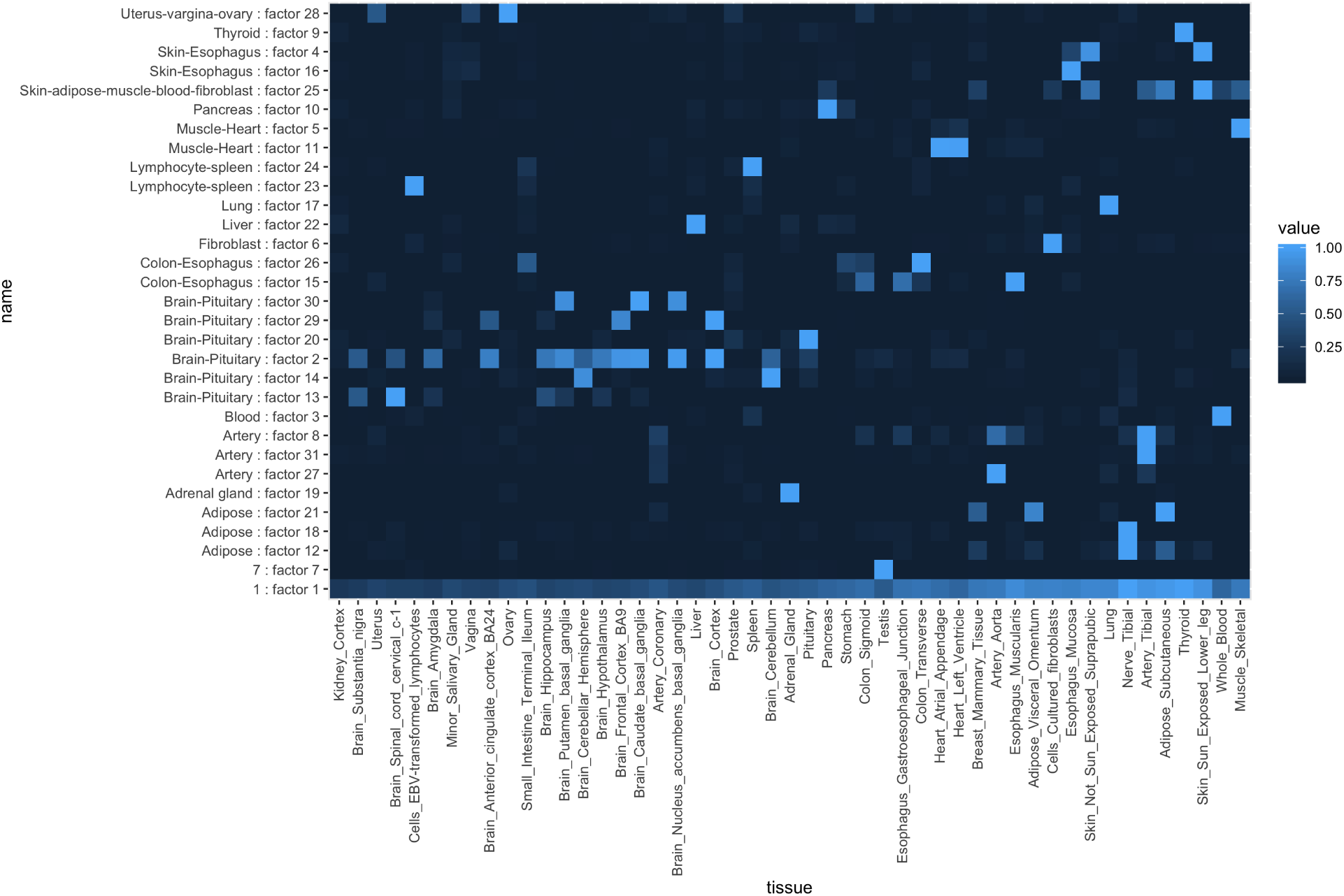
Patterns of tissue sharing identified via factor analysis using flashr. Tissue-pattern categories generated from FLASH applied to the cis-eQTLs are shown. Factor 1 represents cross tissue category covering all tissues, with higher weight for larger sample size tissues. These tissue categories (on y-axis) were used in the analysis of causal tissue identification. Tissues are ordered by sample size.

### 1.15 Supplementary tables in spreadsheet

**Table S9: Presumed causal genes included in the OMIM database**. Columns are: **trait**: Tag used for the trait, **pheno_mim**: MIM ID of the phenotype mapped to GWAS trait, **mim**: MIM ID of the corresponding gene, **entry_type**: Entry type in the OMIM database, **entrez_gene_id**: Gene ID based on Entrez database, **gene_name**: Official gene symbol, **ensembl_gene_id**: Gene ID based on Ensembl database, **gene_type**: Gene type based on Gencode, **gene**: Trimmed Gene ID based on Ensembl database.

**Table S8: GWAS Metadata** contains relevant information concerning each GWAS study used. Full table available in Supplementary Material. Analyses used the 87 traits with deflation=0 unless explictly said otherwise. Columns are: **Tag**: Internal name to identify the study, **Deflation**: Deflation status after imputation (0 for no deflation, 1 for moderate deflation, 2 for extreme deflation), **PUBMED_Paper_Link**: PUBMED entry, **Pheno_File**: name of downloaded file, **Source_File**: actual name of GWAS summary statistics (i.e. downloaded files might contain several traits), **Portal**: URL to GWAS study portal, **Consortium**: Name of Consortium if any, **Link**: download link for the file, **Notes**: any special comment on the GWAS trait, **Header**: GWAS summary statistics header in case the file is malformed, **EFO**: Experimental Factor Ontology [Malone et al., 2010] entry if applicable, **HPO**: Human Phenotype Ontology [Köhler et al., 2013] entry if applicable, **Description**: optional description of the study, **Trait**: trait name, **Sample_Size**: number of individuals included in the study, **Population**: types of populations present (EUR for European, AFR for African, EAS for East Asian, etc), **Date**: Date the file was downloaded, **Declared_Effect_Allele**: column specifying effect allele, **Genome_Reference**: Human Genome release used as reference (i.e. hg19, hg38), **Binary**: wether the trait is dichotomous, **Cases**: number of cases if binary trait, **abbreviation**: short string for figure and table display, **new_abbreviation**: additional abbreviation, **new_Trait**: additional trait name, **Category**: type of trait, **Color**: Hexadecimal color code for display

**Table S10: PrediXcan and enloc results for predicted causal genes selected based on OMIM**. Columns are: **lead_var**: the most significant variant within the LD block, **trait**: trait name, **gene**: Ensembl ID for the gene, **is_omim**: Is included in the OMIM database. TRUE if included, FALSE if not, **proximity**: 0 if variant is in the gene, otherwise BPS from the gene boundary, **rank_proximity**: ranking by proximity within LD block (rank starts from 0 and the closer the lower rank), **percentage_proximity**: rank_proximity / number of genes in the locus, **predixcan_mashr_eur_score**: -log10 p-value (most significant across tissues is used) of PrediXcan-MASH trained on European data, **enloc_score**: rcp (max across tissues), **predixcan_mashr_eur_rank**: PrediXcan significance ranking within LD block (rank starts from 0 and the higher significance the lower rank), **enloc_rank**: enloc rcp ranking within LD block (rank starts from 0 and the higher rcp the lower rank), **predixcan_mashr_eur_percentage**: predixcan_mashr_eur_rank / number of genes in the locus, **enloc_percentage**: enloc_rank / number of genes in the locus, **gene_name**: Official gene symbol, **gene_type**: Gencode annotsted gene type, **chromosome**: Chromosome for the gene, **start**: Gencode annotated gene start position. All isoforms are combined, **end:** Gencode annotated gene end position. All isoforms are combined, **strand**: Gencode annotated gene strand.

**Table S11: PrediXcan and enloc results for presumed causal genes in the rare variant based silver standard**. Columns are: **lead_var**: the most significant variant within the LD block, **trait**: trait name, **gene**: Ensembl ID for the gene, **is_ewas**: Is included in the EWAS. TRUE if included, FALSE if not, **proximity**: 0 if variant is in the gene, otherwise BPS from the gene boundary, **rank_proximity**: ranking by proximity within LD block (rank starts from 0 and the closer the lower rank), **percentage_proximity**: rank_proximity / number of genes in the locus, **predixcan_mashr_score**: -log10 p-value (most significant across tissues is used) of PrediXcan-MASH trained on European data, **enloc_score**: rcp (max across tissues), **predixcan_mashr_rank**: PrediXcan significance ranking within LD block (rank starts from 0 and the higher significance the lower rank), **enloc_rank**: enloc rcp ranking within LD block (rank starts from 0 and the higher rcp the lower rank), **predixcan_mashr_percentage**: predixcan_mashr_eur_rank / number of genes in the locus, **enloc_percentage**: enloc_rank / number of genes in the locus, **gene_name**: Official gene symbol, **gene_type**: Gencode annotsted gene type, **chromosome**: Chromosome for the gene, **start**: Gencode annotated gene start position. All isoforms are combined, **end**: Gencode annotated gene end position. All isoforms are combined, **strand**: Gencode annotated gene strand.

**Table S12: Genes suggested as causal by rare variant association studies**. Columns are: **gene**: Trimmed gene ID based on Ensembl database, **nobs**: Number of times gene has been observed in the trait, **trait**: Tag for the trait name.

**Table S13: OMIM genes included in the analysis**. Columns are: **gene, trait**.

**Table S14: Rare variant silver standard genes included in the analysis**. Columns are: **gene, trait**.

**Table S15: BioVU**. Columns are: **gene, tissue, trait_map**: mapped trait, **pheno**: trait, **gene_name, p_discovery, rcp_discovery, beta_biovu, p_biovu, z_biovu**.

